# Single-component multilayered self-assembling nanoparticles presenting rationally designed glycoprotein trimers as Ebola virus vaccines

**DOI:** 10.1101/2020.08.22.262634

**Authors:** Linling He, Anshul Chaudhary, Xiaohe Lin, Cindy Sou, Tanwee Alkutkar, Sonu Kumar, Timothy Ngo, Ezra Kosviner, Gabriel Ozorowski, Robyn L. Stanfield, Andrew B. Ward, Ian A. Wilson, Jiang Zhu

## Abstract

Ebola virus (EBOV) glycoprotein (GP) can be recognized by neutralizing antibodies (NAbs) and is the main target for vaccine design. Here, we first investigate the contribution of the stalk and heptad repeat 1-C (HR1_C_) regions to GP metastability. Specific stalk and HR1_C_ modifications in a mucin-deleted form (GPΔmuc) increase trimer yield, whereas alterations of HR1_C_ exert a more complex effect on thermostability. Crystal structures are determined to validate two rationally designed GPΔmuc trimers in their unliganded state. We then display a modified GPΔmuc trimer on reengineered nanoparticles that encapsulate a layer of locking domains (LD) and a cluster of helper T-cell epitopes. In mice and rabbits, GP trimers and nanoparticles elicit cross-ebolavirus NAbs, as well as non-NAbs that enhance pseudovirus infection. Repertoire sequencing reveals quantitative profiles of vaccine-induced B-cell responses. This study demonstrates a promising vaccine strategy for filoviruses, such as EBOV, based on GP stabilization and nanoparticle display.

## Introduction

Ebola virus (EBOV), a member of the *Ebolavirus* genus in the *Filoviridae* family^1^, can cause a severe human disease known as viral hemorrhagic fever (VHF)^2,3^. EBOV was solely responsible for the largest filovirus outbreak in history in 2013-2016 that caused 11,325 deaths^4^. The EBOV outbreak in 2019 led to 2,103 deaths^5^ and was declared an international emergency on July 17, 2019, by the World Health Organization (WHO). In recent years, significant progress has been made to counter this deadly virus. Neutralizing antibodies (NAbs) provided effective therapeutics for EBOV infection^6–9^, as demonstrated by the ZMapp cocktail of murine chimeric antibodies^10,11^, as well as human antibodies^12,13^. Vaccines based on different delivery systems have been tested in humans^14–17^, of which rVSV-ZEBOV (Ervebo^®^) – a replication-competent recombinant vesicular stomatitis virus (VSV) expressing a *Zaire* EBOV glycoprotein (GP)^18–21^ – was recently approved by the U.S. Food and Drug Administration (FDA) for human use. However, GP-specific antibody titers did not noticeably increase seven days after rVSV-ZEBOV vaccination in humans^15,22^, in contrast to prior findings in nonhuman primates (NHPs)^23^. Additionally, a recent study reported the neurotropism of rVSV-ZEBOV that resulted in damage to the eye and brain in neonatal mice^24^. Antibody-dependent enhancement (ADE) of infection was also found for antibodies isolated from human survivors^25^, suggesting that weak or non-NAbs induced by a suboptimal vaccine may cause adverse effects. Currently, no protein-based subunit vaccines are available but may be needed to boost the NAb response in the rVSV-ZEBOV-vaccinated population.

EBOV GP, a trimer of GP1-GP2 heterodimers responsible for cell entry^26^, is recognized by the humoral immune response during natural infection^27–29^. The outbreak in 2013-2016 led to an enduring campaign to identify and characterize NAbs for EBOV^30^ and other filoviruses, such as Marburg virus (MARV)^31–33^. As a result, panels of NAbs were isolated from human survivors, vaccinated humans, and immunized animals^12,34–39^. Crystallography^40–43^ and electron microscopy (EM)^44–47^ revealed multiple sites of vulnerability on EBOV GP. A systematic study of 171 monoclonal antibodies (mAbs) defined eight epitopes^48^, six of which can be recognized by broadly neutralizing antibodies (bNAbs)^9^. Meanwhile, over the last decade, HIV-1 vaccine research has been driven largely by a strategy that focuses on bNAb isolation, the structural analysis of bNAb-envelope glycoprotein (Env) interactions, and immunogen design^49,50^. An important milestone in recent HIV-1 research was the development of native-like Env trimers, which have emerged as a promising vaccine platform^51,52^. In contrast to the growing number of EBOV (b)NAbs and their structures with GP, little attention has been given to the rational design of EBOV GP. As class-I viral fusion proteins^53,54^, HIV-1 Env and EBOV GP are inherently metastable, which is a property that has been studied in depth for HIV-1 Env^55–57^, but not yet for EBOV GP. Another advance in the HIV-1 vaccine field was to display Env trimers on self-assembling nanoparticles (NPs)^58,59^. Recombinant virus-like particles (VLPs) can protect against EBOV challenge in animals^60–62^, but manufacturing difficulties have hindered their development as human vaccines^63^. Therefore, the multivalent display of stabilized EBOV GP trimers on NPs may provide a promising solution for developing VLP-type protein subunit vaccines, but this possibility has yet to be explored.

Here, we investigated the causes of EBOV GP metastability and designed multilayered NP immunogens for in vivo evaluation. To facilitate GP purification, we developed an immunoaffinity column based on mAb100^12,42^, which is specific to native-like, trimeric GP. We first examined the contribution of two regions in GP2, namely the stalk and heptad repeat 1-C (HR1_C_) regions, to GP metastability in a mucin-deleted *Zaire* EBOV GP construct (GPΔmuc). We extended the soluble GP ectodomain (GP_ECTO_) in the stalk region from residue 632 (C terminus of HR2) to 637 and introduced a W615L mutation based on a comparison of EBOV and MARV GPs. We also assessed eight proline mutations in HR1_C_, a region equivalent to the HR1 bend that is essential to HIV-1 Env metastability^55–57^, for their ability to prevent the GP transition from pre- to postfusion states. Both the stalk and HR1_C_-proline mutations increased trimer yield, whereas the latter exhibited a complex effect on GP thermostability. In addition, newly engineered inter-protomer disulfide bonds (SS) were tested for their ability to increase trimer stability. Crystal structures were solved for two redesigned GPΔmuc trimer constructs to validate the stalk and HR1_C_-proline mutations at the atomic level. We then displayed a redesigned GPΔmuc trimer on ferritin (FR), E2p, and I3-01 NPs. Locking domains (LDs) and helper T-cell epitopes were incorporated into E2p and I3-01 60-mers^55,64^ to stabilize the NP shell from the inside and create multilayered NP carriers. In mice and rabbits, GP trimer and NP vaccines induced distinct antibody responses, but enhanced pseudoviral infection was observed for some constructs. The next-generation sequencing (NGS) of GP-specific B cells revealed different patterns for NPs that presented large trimeric spikes versus smaller antigens, such as hepatitis C virus (HCV) E2 core^65^. Our study thus reports on critical factors for EBOV GP metastability, single-component multilayered self-assembling NPs for the development of VLP-type vaccines, and a subunit vaccine strategy that is applicable to other filoviruses.

## Results

### Tag-free immunoaffinity purification of EBOV GP trimers

EBOV GP contains a heavily glycosylated mucin-like domain (MLD) that shields the glycan cap and neutralizing epitopes in GP1 and GP2 (**Fig. 1a**). A soluble, mucin-deleted form of *Zaire* EBOV GP (GPΔmuc) that produced the first^40^ and subsequent high-resolution^66^ crystal structures was used as a basic construct to investigate GP metastability (**Fig. 1a**). In HIV-1 vaccine research, immunoaffinity chromatography (IAC) columns based on bNAbs 2G12 and PGT145^67,68^ have been widely used to purify native-like Env trimers. 2G12 targets a glycan patch on a single gp120, whereas PGT145 binds the trimer apex and can separate closed trimers from partially open and misfolded Envs. These two bNAb columns have also been used to purify HIV-1 gp140-presenting NPs^55,64,69^. Likewise, GP-specific IAC columns may provide useful tools for EBOV vaccine research. Corti et al. recently identified two potent NAbs, mAb114 and mAb100, from a human survivor^12^. Misasi et al. elucidated the structural basis for neutralization by mAb114, which targets the receptor binding site (RBS), and mAb100, which interacts with the GP1/GP2 interface and internal fusion loop (IFL) of two GP protomers^42^. Here, we examined the utility of mAb114 and mAb100 in IAC columns. The GPΔmuc constructs with and without a C-terminal foldon motif were transiently expressed in 250 ml HEK293F cells and purified on an antibody column prior to size-exclusion chromatography (SEC) using a Superdex 200 10/300 GL column and blue native polyacrylamide gel electrophoresis (BN-PAGE). With mAb114, both GPΔmuc samples showed aggregate (∼9 ml), dimer (∼12 ml), and monomer (∼13.5-14 ml) peaks in the SEC profiles, but only GPΔmuc-foldon showed a visible trimer peak (∼10.5-11 ml) in SEC and a band of slightly less than 440 kD on the BN gel (**Fig. 1b**). Following mAb100 purification, GPΔmuc produced a low overall yield, whereas GPΔmuc-foldon demonstrated high trimer purity without any monomer or dimer peaks. Consistently, GPΔmuc-foldon registered a single trimer band on the BN gel (**Fig. 1c**). Altogether, both mAb114 and mAb100 can be used to purify EBOV GP, but mAb100 offers a more effective IAC method for purifying native-like trimers due to its recognition of a quaternary epitope. The stark difference in trimer yield between the two GPΔmuc constructs after mAb100 purification also suggests a strong tendency for trimer dissociation without foldon.

**Fig. 1.**
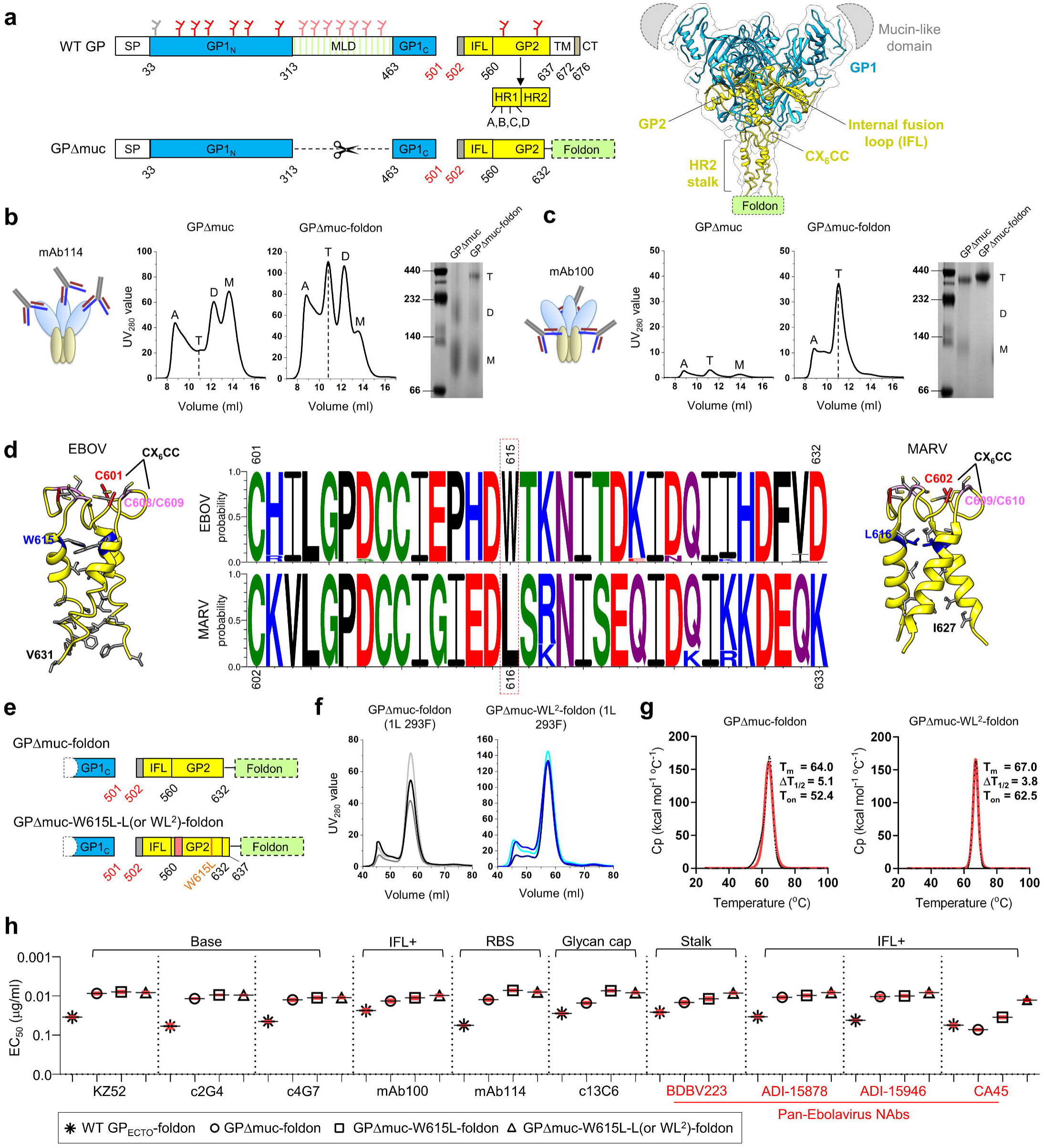
Design and characterization of EBOV GPΔmuc trimers with modified HR2 stalk. (**a**) Left: Schematic representation of EBOV GP and GPΔmuc. GP1 N/C termini (GP1_N/C_), mucin- like domain (MLD), internal fusion loop (IFL), heptad repeat regions 1/2 (HR1/HR2), transmembrane region (TM), and cytoplasmic tail (CT) are labeled with *N*-linked glycans, which are indicated as gray (mutated), red, and pink (within MLD) branches. Right: Ribbon representation of EBOV GP (PDB: 5JQ3) in transparent molecular surface, with GP1 in dodger blue and GP2 in yellow. The MLD and foldon are shown as a gray half oval and a green rectangle, respectively. **(b)** Schematic representation of mAb114 bound to EBOV GP (left), SEC profiles of mAb114-purified GPΔmuc and GPΔmuc-foldon (middle) from a Superdex 200 10/300 column, and BN-PAGE gel (right). **(c)** Schematic representation of mAb100 bound to EBOV GP (left), SEC profiles of mAb100-purified GPΔmuc and GPΔmuc-foldon (middle) from a Superdex 200 10/300 column, and BN-PAGE gel (right). In (b) and (c), GP species (A: aggregate; T: trimer; D: dimer; M: monomer) are labeled on the SEC profile and BN-PAGE gel. **(d)** Ribbon representation of EBOV HR2 stalk (left) and MARV HR2 stalk (right) with CX_6_CC motif and residues of interest indicated. Middle: Logo analysis of EBOV and MARV HR2 sequences. **(e)** Schematic representation of GPΔmuc-W615L-L (or WL^2^)-foldon. **(f)** SEC profiles of 293F-expressed, mAb100-purified GPΔmuc-foldon and GPΔmuc-WL^2^-foldon from a HiLoad Superdex 200 16/600 column for three production runs. **(g)** Thermostability of GPΔmuc-foldon and GPΔmuc-WL^2^-foldon, with T_m_, ΔT_1/2_, and T_on_ measured by DSC. **(h)** EC_50_ (μg/ml) values of EBOV GP-foldon, GPΔmuc-foldon, GPΔmuc-W615L-foldon, and GPΔmuc-WL^2^-foldon binding to 10 representative antibodies. Four pan-Ebolavirus NAbs are colored in red. Antibody binding was measured by ELISA in duplicates, with mean value and standard deviation (SD) shown as black and red lines, respectively. Source data are provided as a Source Data file.

### Effect of HR2 stalk on EBOV GP metastability

Previously, we demonstrated that the gp41 ectodomain (gp41_ECTO_) is the source of HIV-1 Env metastability^55^. Unlike HIV-1 Env^70,71^, in which the gp41 HR2 helix is packed against the bottom of the gp41 HR1 helix and gp120 C1/C5 loops and forms extensive interactions at the gp140 trimer base^70,71^, EBOV GP has a long, extended HR2 stalk (**Fig. 1a**, right). Even in the high-resolution GPΔmuc structures^66,72,73^, the HR2 stalk still contains less helical content than most coiled-coils in the Protein Data Bank (PDB), ∼15 versus ∼30 aa, and becomes less regular and unwound toward the C terminus, suggesting an inherent instability in HR2 (**Fig. 1d**, left). Recently, King et al. solved a 3.17 Å-resolution structure for MARV GPΔmuc bound to a human mAb, MR191^32^. Surprisingly, the MARV HR2 stalk adopted a well-formed coiled-coil with canonical sidechain packing along the three-fold axis (**Fig. 1d**, right). To identify the cause of this difference in HR2, we obtained EBOV and MARV GP sequences from the Virus Pathogen Database and Analysis Resource (ViPR, https://www.viprbrc.org). A total of 274 EBOV GPs and 87 MARV GPs were used for sequence conservation analysis of the region spanning the CX_6_CC motif and HR2 stalk, residues 601-632 for EBOV and residues 602-633 for MARV, respectively (**Fig. 1d**, middle). Most inward-facing amino acids were conserved except for W615 in EBOV and the equivalent L616 in MARV. Indeed, structural analysis revealed a critical difference at this position: the three W615s in EBOV GP (PDB: 5JQ3) formed a wide triangle at the neck of the coiled-coil with a Cα distance of 11.1 Å and Cβ distance of 9.0 Å; in contrast, with the smaller and more hydrophobic L616, a Cα distance of 10.5 Å and Cβ distance of 8.3 Å were observed in MARV GP (PDB: 6BP2). This finding suggests that a W615L mutation may improve the stability of EBOV GP.

To further examine the effect of the HR2 stalk on GP trimer stability, we created three GPΔmuc constructs by replacing residues 617-632 with the coiled-coil from a GCN4 leucine zipper (PDB: 2WPZ, residues 3-33) and by extending the C terminus to D637 and N643 to include a newly identified bNAb epitope^74^ that spans HR2 and the membrane-proximal external region (MPER), termed “L” and “Ext”, respectively (**Fig. S1a**). Notably, D637 is also important for proteolytic cleavage by tumor necrosis factor α-converting enzyme (TACE), which enables GP to be shed from the virus surface^75^. These three constructs were characterized by SEC and BN-PAGE following transient expression in 250-ml HEK293F cells and purification on an antibody column. With mAb114 purification, all three HR2 stalk variants showed a greater trimer yield than wildtype GPΔmuc in SEC (**Fig. S1b**), with trimer bands observed only for the stalk variants on the BN gel (**Fig. S1c**, left). Upon mAb100 purification, all three HR2 stalk variants showed more visible trimer bands than wildtype GPΔmuc on the BN gel (**Fig. S1c**, right). Of the three designs, “2WPZ” improved GP stability at the cost of replacing the entire HR2 stalk in GP2 but provided supporting evidence for the W615L mutation, which presumably increases coiled-coil content in the HR2 stalk (**Fig. 1d**). Overall, the “L” extension appeared to be a more practical solution as it modestly improved trimer stability with the inclusion of a recently identified bNAb epitope^74^.

We next combined the W615L mutation and “L” extension in a single construct named GPΔmuc-W615L-L-foldon, or simply GPΔmuc-WL^2^-foldon (**Fig. 1e**). This construct, along with GPΔmuc-foldon, was transiently expressed in 1-liter HEK293F cells and purified by an mAb100 column prior to SEC on a HiLoad Superdex 200 16/600 GL column (**Fig. 1f**). In three production runs, GPΔmuc-WL^2^-foldon consistently outperformed the wildtype construct, showing a twofold higher trimer peak in the SEC profile and a ∼2.6-fold greater trimer yield after SEC (1.3 mg versus 0.5 mg). Thermostability was assessed by differential scanning calorimetry (DSC) for two purified GP trimers (**Fig. 1g**). The thermal denaturation midpoint (T_m_) value for the stalk-stabilized trimer was 3 °C higher than the wildtype trimer (67 °C versus 64 °C). Stalk stabilization also increased the onset temperature (T_on_) from 52.4 °C to 62.5 °C, with a narrower half width of the peak (ΔT_1/2_) than the wildtype trimer (3.8 °C versus 5.1 °C). Antigenicity was assessed for four mAb100/SEC-purified EBOV GP trimers in the enzyme-linked immunosorbent assay (ELISA) (**Fig. 1h, Fig. S1d** and **S1e**). Ten antibodies were tested, including three NAbs targeting the base (KZ52^76^, c2G4, and c4G7^10^), two human NAbs^12^ – mAb100 (IFL) and mAb114 (RBS), a non-NAb directed to the glycan cap (c13C6^10^), and four pan-ebolavirus bNAbs targeting the conserved HR2-MPER epitope (BDBV223^74^) and IFL (ADI-15878, ADI-15946^37,38^, and CA45^36,77^). The GPΔmuc trimer showed notably improved antibody binding with respect to the GP_ECTO_ trimer, with an up to 7.6-fold difference in the half maximal effective concentration (EC_50_), indicating that the MLD can shield much of the GP from antibody recognition. The two stalk modifications only modestly increased recognition of the RBS, stalk, and IFL epitopes by their respective (b)NAbs except for CA45^36,77^, for which the WL^2^ mutation led to a 5.6-fold change in EC_50_ compared with GPΔmuc-foldon. A 50% reduction in EC_50_ observed for GPΔmuc-WL^2^-foldon binding to BDBV223 confirmed that the L extension into MPER can improve the recognition of this conserved bNAb epitope at HR2-MPER^74^. King et al. proposed two scenarios for ebolavirus GP to expose the BDBV223 epitope: one of the HR2 helices is splayed apart from the coiled-coil, or the whole GP is lifted or tilted with respect to the membrane^74^. It is perhaps more plausible that ebolavirus GP may adopt an open stalk conformation similar to the parainfluenza virus 5 (PIV5) fusion (F) protein, which has a long HR-B stalk^78^. Altogether, WL^2^ considerably improved the trimer yield and thermostability for EBOV GP but only exerted a modest effect on antigenicity, because the C-terminal foldon motif in these constructs could retain GPΔmuc in a trimeric form, which is required for (b)NAb binding.

### Effect of the HR1_C_ bend on EBOV GP metastability

Structures of the prefusion glycoprotein and the postfusion six-helix-bundle have been determined for HIV-1 Env^70,71,79^ and EBOV/MARV GP^40,80–82^. Previously, we identified the N terminus of HR1 (HR1_N_, residues 547-569) as a major cause of HIV-1 Env metastability^56^, because this region undergoes a drastic conformational change during fusion. We and others stabilized diverse HIV-1 Envs by replacing the 22-aa HR1_N_ with an optimized 8-aa loop^55–57,83^. The equivalent region in EBOV GP (residues 576-583), HR1_C_, forms a 21-nm triangular interior at around two-thirds of the height of the chalice that spans 94-Å wide at the rim (**Fig. 2a**). This prefusion HR1_C_ bend will refold to two helical turns in the center of a long helix in the postfusion state (**Fig. 2b**, left). Here, we hypothesized that HR1_C_ is essential to EBOV GP metastability. Since HR1_C_ in wildtype EBOV GP is equivalent in length (8 aa) to a truncated engineered HR1_N_ in the prefusion-optimized HIV-1 Env^55,56^, metastability in EBOV GP may not be as sensitive to HR1_C_ length and likely requires a different solution. We thus hypothesized that a proline mutation in one of the eight residues in HR1_C_ can rigidify the HR1_C_ bend and improve EBOV GP trimer stability (**Fig. 2b**, right). Similar proline mutations in HR1_N_ have effectively stabilized HIV-1 Env trimers^56,68^. To examine this possibility, eight GPΔmuc-W615L variants, each with a proline mutation in HR1_C_ (termed P^1^ to P^8^) but without the L extension and C-terminal foldon motif, were characterized by SEC after 250-ml 293 F expression and IAC. After mAb114 purification, most proline mutations had little effect on the distribution of GP species except for T577P (P^2^) and L579P (P^4^), which appeared to have a trimer peak at ∼11 ml in their SEC profiles (**Fig. 2c**). After mAb100 purification, only P^2^ and P^4^ produced any trimer yield, with a higher SEC peak observed for P^2^ that corresponded to well-formed trimers (**Fig. 2c**). The mAb100-purified GP was analyzed by BN-PAGE, which showed a trimer band for P^2^ and P^4^ (**Fig. 2d**). Overall, the T577P mutation (P^2^) can considerably increase trimer yield, whereas the L579P mutation (P^4^) has a less pronounced effect.

**Fig. 2.**
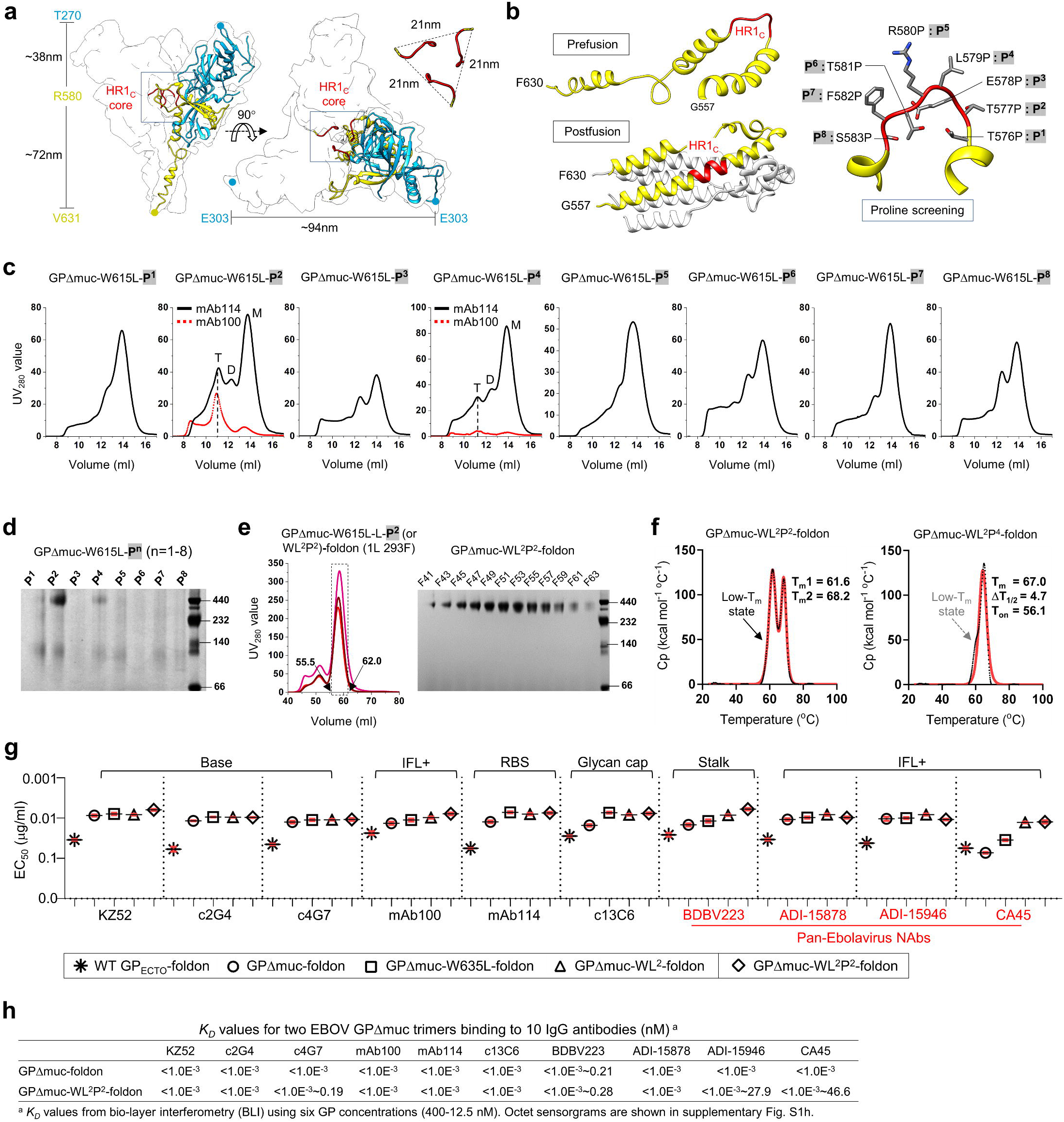
Design and characterization of EBOV GPΔmuc trimers with modified HR1_C_ bend. **(a)** Ribbon representation of EBOV GPΔmuc protomer (PDB: 5JQ3) in transparent molecular surface with GP1 in dodger blue, GP2 in yellow, and HR1_C_ bends from three protomers in red. Left: side view. Right: top view. **(b)** Left: Ribbon representation of the HR1 region in the prefusion (top, PDB ID: 5JQ3) and postfusion (bottom, PDB ID: 2EBO) states with the HR1 region in yellow and the HR1_C_ bend in red. Right: Zoomed-in view of the HR1_C_ bend with the eight residues in this region shown as sticks and labeled with the proline mutations, P^1^-P^8^. **(c)** SEC profiles of mAb114-purified GPΔmuc-W615L-P^n^ variants (n=1 to 8) from a Superdex 200 10/300 column. SEC profiles of mAb100-purified GPΔmuc-W615L-P^2^ and -P^4^ are shown in the red dotted line. Trimer (T), dimer (D), and monomer (M) peaks are labeled on the SEC profiles for P^2^ and P^4^, with the trimer peak marked with a black dashed line. **(d)** BN-PAGE gel of mAb114-purified GPΔmuc-W615L-P^n^ variants (n=1 to 8). **(e)** SEC profiles of 293F-expressed, mAb100-purified GPΔmuc-W615L-L-P^2^ (or WL^2^P^2^)-foldon from a HiLoad Superdex 200 16/600 column (left) and BN-PAGE gel of SEC fractions 41-63 (55.5-62.0 ml) (right). SEC profiles were from three production runs. **(f)** Thermostability of GPΔmuc-WL^2^P^2^-foldon and GPΔmuc-WL^2^P^4^-foldon, with Tm, ΔT_1/2_, and Ton measured by DSC. (**g**) EC_50_ (μg/ml) values of EBOV GP-foldon, GPΔmuc-foldon, GPΔmuc-W615L-foldon, GPΔmuc-WL^2^-foldon, and GPΔmuc-WL^2^P^2^-foldon binding to 10 representative antibodies. Four pan-Ebolavirus NAbs are colored in red. Antibody binding was measured by ELISA in duplicates, with mean value and standard deviation (SD) shown as black and red lines, respectively. (**h**) *K_D_* values of GPΔMuc-foldon and GPΔmuc-WL^2^P^2^-foldon binding to 10 representative antibodies. BLI was performed on an Octet RED96 instrument using a trimer titration series of six concentrations (400-12.5 nM by twofold dilution) and kinetics (AHC) biosensors. The *K_D_* values were calculated using a global fit 2:1 model. Source data are provided as a Source Data file.

Next, the T577P mutation (P^2^) was incorporated into the GPΔmuc-WL^2^-foldon construct, resulting in a construct named GPΔmuc-WL^2^P^2^-foldon. This construct was transiently expressed in 1-liter 293 F cells and purified using an mAb100 column for SEC characterization on a HiLoad Superdex 200 16/600 GL column. In three production runs, GPΔmuc-WL^2^P^2^-foldon generated a trimer peak that was two- and fourfold higher than GPΔmuc-WL^2^-foldon and wildtype GPΔmuc-foldon, respectively, with an average yield of 2.6 mg after SEC (**Fig. 2e**, left). Protein collected in the range of 55.5-62.0 ml was analyzed by BN-PAGE, which displayed a trimer band across all fractions without any detectable impurity (**Fig. 2e**, right). The thermostability of GPΔmuc-WL^2^P^2^-foldon was determined by DSC after mAb100 and SEC purification. Unexpectedly, two transition peaks were observed in the thermogram, one registered at a lower T_m_ of 61.6 °C and the other at a higher T_m_ of 68.2 °C (**Fig. 2f**, left). To this end, a second construct containing the L579P mutation (P^4^), termed GPΔmuc-WL^2^P^4^-foldon, was also assessed by DSC (**Fig. 2f**, right). Although only one peak was observed in the thermogram with a T_m_ of 67.0 °C, a slight widening at the onset of the peak suggested a similar unfolding behavior upon heating. Thus, DSC revealed the complexity associated with a proline-rigidified HR1_C_ bend, which may increase the trimer yield at the cost of reducing or modifying the GP thermostability profile. The antigenicity of GPΔmuc-WL^2^P^2^-foldon was first assessed by ELISA using the same panel of 10 antibodies (**Fig. 2g, Fig. S1f** and **S1g**). GPΔmuc-WL^2^P^2^-foldon appeared to bind more favorably to bNAb BDBV223 than GPΔmuc-WL^2^-foldon, with a twofold reduction in EC_50_. In the bio-layer interferometry (BLI) analysis (**Fig. 2h** and **Fig. S1h**), the GPΔmuc-WL^2^P^2^-foldon trimer and wildtype GPΔmuc-foldon trimer yielded nearly indistinguishable kinetic profiles, with nano- to picomolar equilibrium dissociation constant (*K_D_*) values, consistent with the fast on-rates and slow off-rates in antibody binding.

Our results demonstrated the importance of HR1_C_ to EBOV GP metastability and the perhaps unexpected sensitivity of HR1_C_ to proline mutation. Recently, Rutten et al. tested proline mutations in HR1_C_ along with a K588F mutation to stabilize filovirus GP trimers^84^. A similar increase in trimer yield was observed for the T577P mutant, but the reported thermostability data appeared to be inconsistent with our DSC measurements. Further investigations are thus warranted to understand the role of HR1_C_ in filovirus-cell fusion and GP stability. Nevertheless, the combined stalk/HR1_C_ modification, WL^2^P^2^, appeared to have no effect on GP binding to diverse (b)NAbs.

### GP stabilization with inter-protomer disulfide bonds

Engineered disulfide (SS) bonds have been used to stabilize viral glycoproteins, as demonstrated for HIV-1 Env^68^, respiratory syncytial virus (RSV) F^85^, and Lassa virus (LASV) GP complex (GPC)^86^. EBOV GP already contains an endogenous SS bond linking GP1 and GP2 (C53-C609). Thus, we examined whether inter-protomer SS bonds can be used to stabilize trimer formation and lock the GP in a “closed” trimer. Based on a high-resolution EBOV GPΔmuc structure (PDB ID: 5JQ3), we identified inter-protomer amino acid pairs whose C_β_-C_β_ distances are within a cutoff of 4.72 Å defined by Rosetta^87^ (**Fig. S2a**). Of the nine pairs that were identified, three were removed after visual inspection, because they may interfere with an existing SS bond or a hydrophobic cluster. The remaining six pairs were divided into three groups: IFL-head group (SS1/2/4), IFL-NHR group (SS/5), and HR2 group (SS6) (**Fig. S2b**). Six GPΔmuc-SS constructs without C-terminal foldon were designed and characterized by SEC following transient expression in 250-ml 293 F cells and purification using an mAb114 column or mAb100 column (**Fig. S2c**). Diverse SEC profiles were observed for the mAb114-purified SS variants, with SS2 showing a notable trimer fraction. Upon mAb100 purification, only SS2, SS3, and SS5 resulted in a measurable protein yield, with SS2 showing a clean trimer peak without dimer/monomer species. The BN-PAGE analysis of mAb114 and mAb100-purified SS variants exhibited consistent patterns, with S22 showing a trimer band slightly below 440 kD on the gel (**Fig. S2d**). However, the incorporation of SS2, SS3, or SS5 into the GPΔmuc-foldon construct led to abnormal SEC profiles regardless of the antibody column used for purification, suggesting their incompatibility with the foldon trimerization motif (**Fig. S2e**). To this end, antigenicity was assessed only for the mAb100/SEC-purified GPΔmuc-SS2 trimer. In the ELISA, GPΔmuc-SS2 showed identical or improved antibody binding compared with GPΔmuc-foldon, with a 3.7-fold reduction in EC_50_ for bNAb CA45 (**Fig. S2f** and **S2g**). In summary, a well-positioned inter-protomer SS bond (e.g., between the GP1 head and GP2 IFL) can stabilize the EBOV GP trimer, with no adverse effect on antigenicity.

### Crystallographic analysis of redesigned EBOV GPΔmuc trimers

To understand how the HR2 stalk and HR1_C_ mutations affect EBOV GP, we determined crystal structures for the unliganded GPΔmuc-foldon with WL^2^ and WL^2^P^2^ at 2.3 Å and 3.2 Å, respectively (**Fig. S3, Fig. 3**). Both proteins showed a three-lobed, chalice-like trimer structure^40,66^. WL^2^/WL^2^P^2^ yielded Cα root-mean-square deviations (RMSDs) of 0.9/1.6 Å (for 367/382 residues) at the single protomer level and 1.25/1.23 Å (for 1103/1145 residues) at the trimer level, respectively, relative to wildtype GP (PDB ID: 5JQ3)^66^. WL^2^P^2^ yielded a more complete structure than WL^2^ at the glycan cap (R302-V310) and HR2 stalk (I627-D637) (**Fig. S3, Fig. 3**). In the WL^2^P^2^ structure, the glycan cap covers the RBS with glycan moieties visible for N238/N257/N268 in GP1 and N563 in GP2. In the WL^2^ structure, the glycan cap covers the RBS with glycan moieties visible for N238/N257 in GP1 and N563/N618 in GP2 (**Fig. S3b**). GP1 consists mainly of β-strands, which form a broad semicircular groove that clamps the α3 helix and β19-β20 strands in GP2 (**Fig. 3a**). The T577P mutation appeared to have minimal effect on conformation of the HR1_C_ bend, as indicated by a Cα RMSD of 0.2 Å for this 8-aa segment (**Fig. 3b**, left). In the WL^2^P^2^ structure, the backbone carbonyl (CO) groups of R574, A575, and T576 in one protomer formed moderate-to-strong hydrogen bonds with the guanidinium moiety of R164 in an adjacent protomer, whereas only one CO-NH distance was within the 3.5 Å cutoff in wildtype GPΔmuc^66^. We previously hypothesized that a bulky, inward-facing W615 at the top of the coiled-coil destabilizes the HR2 stalk, whereas the W615L mutation would improve its packing (**Fig. 1d**). Indeed, W615L exerted a positive effect on the stalk structure (**Fig. 3b**, bottom center and right). The Cα-Cα/Cβ-Cβ distances between two W615s of adjacent protomers in wildtype GPΔmuc^66^, 11.1/9.0 Å, were reduced to 10.1/8.0 Å and 10.6/8.2 Å in WL^2^ and WL^2^P^2^, respectively (**Fig. S3, Fig. 3b**, bottom center and right). As a result, the coiled-coil region in the EBOV HR2 stalk added one more helical turn (D624-V631), thereby resembling the MARV HR2 stalk (**Fig. 1d**, right). The L extension in the WL^2^P^2^ structure could be fully modeled to D637 as a well-ordered loop anchored to the C-terminal foldon motif (**Fig. 3b**, bottom center), rendering a complete HR2 stalk and partial MPER. The superposition of HR2 stalks, from R596 up to D637, yielded Cα RMSDs of 1.5 Å, 2.1 Å, and 1.9 Å relative to EBOV-Mayinga (PDB ID: 5JQ3), SUDV (PDB ID: 3S88), and BDBV (PDB ID: 6EA5) GPs, respectively (**Fig. 3b**, right), suggesting inherent structural variability in this region.

**Fig. 3.**
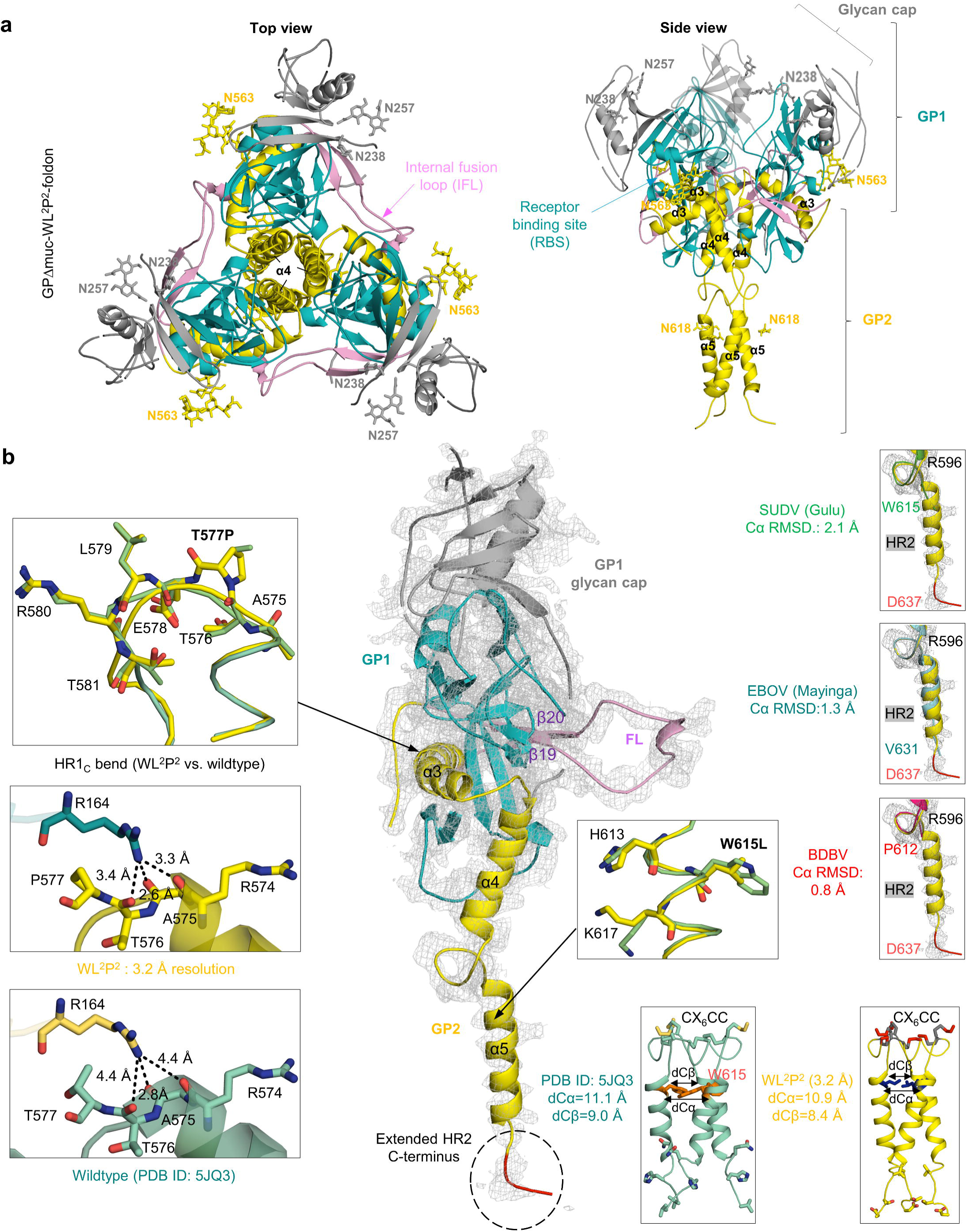
Structural characterization of EBOV GP with stalk and HR1_C_ mutations. **(a)** The 3.2Å-resolution crystal structure of EBOV GPΔmuc-WL^2^P^2^-foldon in a ribbon representation (top view and side view). GP1 is shown in dodger blue, except for the glycan cap, which is in gray. GP2 is shown in yellow with the internal fusion loop (IFL) in pink. *N*-linked glycans at N238, N257, and N563 are shown as sticks. **(b)** Ribbon representation of a GPΔmuc-WL^2^P^2^ protomer in the center with inset images showing structural comparison for the HR1_C_ bend (left), W615-sorrounding HR2 region (bottom right), and C terminus of the HR2 stalk (right). For the HR1_C_ bend, WL^2^P^2^ is superimposed onto GPΔmuc (PDB ID: 5JQ3) (top) with three hydrogen bonds labeled for WL^2^P^2^ (middle) and GPΔmuc (bottom). For the W615-sorrounding HR2 region, WL^2^P^2^ is superimposed onto GPΔmuc (PDB ID: 5JQ3) (top) with the coiled-coil structure shown for GPΔmuc (left) and WL^2^P^2^ (right). Cα and Cβ distances for residue 615 around the threefold axis are labeled. For the HR2 C terminus, WL^2^P^2^ is superimposed onto GP structures of SUDV (top), EBOV (middle), and BDBV (bottom) with Cα RMSDs calculated after fitting. The 2Fo – Fc electron density map contoured at 1σ is shown as a gray mesh for the WL^2^P^2^ protomer (center) and HR2 stalks (right).

The WL^2^P^2^ structure was then compared to a recently reported *Makona* GPΔmuc structure (PDB ID: 6VKM, 3.5 Å) that contained the T577P/K588F mutation (**Fig. S4**). In total, 353 of 398 residues in the WL^2^P^2^ structure matched the double mutant with a Cα RMSD of 0.9 Å. A more complete cathepsin cleavage loop (between β13 and β14, residues 190-210) was observed in WL^2^P^2^ than in the double mutant, showing 10 more flanking residues, five on each side, of the loop that bridges over the IFL and interacts with IFL-directed NAbs such as mAb100^42^. In addition, more electron density was observed for the β18 loop of the glycan cap (residues 294-310) and the stalk in WL^2^P^2^ than in the double mutant (**Fig. S4b**, top right). For the HR1 bend, WL^2^P^2^ showed a Cα RMSD of 0.3 Å and more favorable hydrogen bonding patterns (**Fig. S4b**, bottom left). A Cα RMSD of 1.7 Å was obtained for the IFL region between the two structures (**Fig. S4b**, bottom right). Lastly, the WL^2^P^2^ structure was docked into a panel of known GP/antibody complexes (**Fig. S5a**). Overall, WL^2^P^2^ preserved all critical GP-antibody interactions (**Fig. S5b**). The mAb100/GP complex is of most interest, because mAb100 was used to purify GP trimers. Cryo-EM revealed additional density near the mAb100 light chain that likely corresponds to portions of the cathepsin cleavage loop^42^, but this density was not observed in a 6.7 Å crystal structure of the same complex (PDB ID: 5FHC) ^42^. In the WL^2^P^2^ structure, the flanking region on the β13 side extended to H197 (**Fig. S4**), which would be in proximity to the mAb100 light chain in the WL^2^P^2^/mAb100 complex.

The crystal structures validated the stalk mutation WL^2^ and its combination with the HR1_C_ mutation, WL^2^P^2^, in addition to providing atomic details for regions that were absent in previously reported GP structures. The WL^2^P^2^ structure also provides an explanation for the higher trimer yield (the formation of more favorable inter-protomer hydrogen bonds), although the cause of the two-peak thermogram remains unclear. Notably, GPΔmuc-SS2 was not structurally characterized in this study, because its incompatibility with foldon posed a challenge to crystallization.

### Display of EBOV GPΔmuc trimers on multilayered hyperstable nanoparticles

VLPs are intrinsically immunogenic due to their large size and dense antigen display^88^. Compared with small antigens, VLPs are more suitable for direct uptake by dendritic cells (DC) and clustering of B-cell receptors (BCRs)^88^. Although recombinant VLPs can protect against EBOV challenge^60–62^, they may not be optimal vaccine solutions because of abnormal filament structures (up to 14 μm long) and manufacturing challenges^63^. Recently, self-assembling protein NPs were considered an alternative platform for developing VLP vaccines^58,59^. Previously, we displayed gp41-stabilized HIV-1 Env trimers on protein NPs of various sizes, which elicited robust NAb responses in mice and rabbits^55,64^. We also reported protein NPs that present an optimized HCV E2 core, which induced cross-genotype NAb responses in mice^65^. In this study, we displayed rationally redesigned GPΔmuc trimers on 24- and 60-meric protein NPs for in vivo assessment.

To explore this possibility, we modeled the EBOV GPΔmuc trimer on FR, E2p, and I3-01, resulting in GP-presenting NPs with diameters of 34.5 nm, 45.9 nm, and 49.2 nm, respectively (**Fig. 4a**). Superposition of GPΔmuc C-termini onto FR and E2p N-termini yielded Cα RMSDs of 7.0 Å and 5.5 Å, suggesting that GPΔmuc can be fused to FR with a short G_4_S linker and to E2p without a linker, respectively. However, the large spacing between the N termini of I3-01 subunits (∼50.5 Å) requires a long linker to connect with the C-termini of a GPΔmuc trimer, which form a long, narrow stalk. Computational modeling suggested a 10-aa (G_4_S)_2_ linker, which would result in a Cα RMSD of 0.8 Å. Here, we first displayed two GPΔmuc trimers, wildtype and WL^2^P^2^, on FR, E2p, and I3-01 with a 5-aa linker, no linker, and a 10-aa linker, respectively. WL^2^P^2^, instead of WL^2^, was selected for NP display for its high trimer propensity and atomic structure. All six GP-NP fusion constructs were transiently expressed in 100-ml ExpiCHO cells followed by mAb100 purification and SEC on a Superose 6 10/300 GL column (**Fig. 4b**). WL^2^P^2^ outperformed wildtype GPΔmuc with greater NP yield and purity. Based on molecular weight (m.w.), the SEC peaks at ∼15 ml correspond to the unassembled GP-NP species, suggesting an inherent instability for wildtype E2p and I3-01. The mAb100-purified GPΔmuc-WL^2^P^2^-presenting NP samples were further analyzed by negative stain EM (**Fig. 4c**), showing NPs mixed with impurities.

**Fig. 4.**
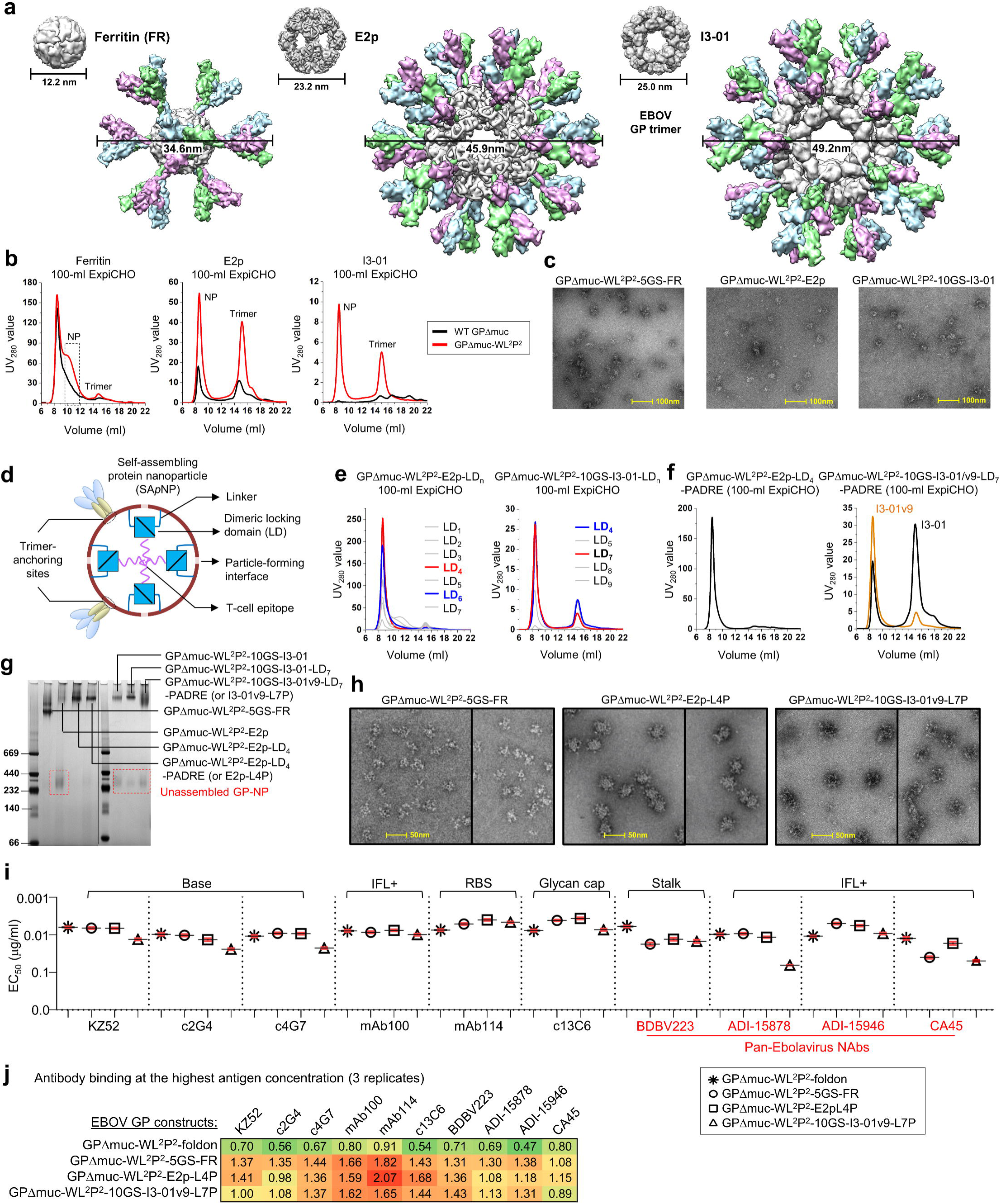
Design and characterization of EBOV GPΔMuc-presenting nanoparticles. (**a**) Surface models of nanoparticle (NP) carriers and GPΔMuc-presenting NPs. The three NP carriers shown here are 24-meric ferritin (FR) and 60-meric E2p and I3-01. The NP size is indicated by diameter (in nm). **(b)** SEC profiles of wildtype GPΔMuc (black) and WL^2^P^2^ (magenta)-presenting FR, E2p, and I3-01 NPs obtained from a Superose 6 10/300 GL column after mAb100 purification. The particle fraction is indicated by a dotted-line box for FR. **(c)** Negative stain EM images of SEC-purified GPΔMuc-WL^2^P^2^-presenting FR, E2p, and I3-01 NPs. **(d)** Schematic representation of multilayered NP design, in which a dimeric locking domain (LD) is fused to the C-terminus of an NP subunit, and a helper T-cell epitope (PADRE) is fused to the C-terminus of an LD. **(e)** SEC profiles of GPΔMuc-WL^2^P^2^-presenting E2p NPs with LDs 1-7 and I3-02 NPs with five LDs (4-5 and 7-9) after mAb100 purification. **(f)** SEC profiles of GPΔMuc-WL^2^P^2^-presenting E2p NP with LD4 and PADRE, or E2p-L4P (left), and I3-01/v9 NP with LD7 and PADRE, or I3-01/v9-L7P (right). I3-01v9 is a variant of I3-01 with a redesigned NP-forming interface. **(g)** BN-PAGE of GPΔMuc-WL^2^P^2^-presenting FR, E2p, and I3-01/v9 NPs, with LD and PADRE variants included for E2p and I3-01/v9. Low molecular weight (m.w.) bands are circled with red dotted lines. Black line indicates the gels on the left and right were generated from two separate experiments. **(h)** Negative-stain EM images of SEC-purified FR, E2p-L4P, and I3-01v9-L7P NPs that present the GPΔMuc-WL^2^P^2^ trimer. Samples are shown as a composite of two panels, each representing a different micrograph. **(i)** EC_50_ (μg/ml) values of GPΔMuc-WL^2^P^2^-foldon and GPΔMuc-WL^2^P^2^-presenting NPs binding to 10 respective antibodies. Four pan-Ebolavirus NAbs are colored in red. Antibody binding was measured by ELISA in duplicates, with mean value and standard deviation (SD) shown as black and red lines, respectively. **(j)** Antigenic profiles of GPΔMuc-WL^2^P^2^-foldon and GPΔMuc-WL^2^P^2^-presenting NPs against 10 antibodies. Two BLI experiments were performed with three replicates tested for the highest antigen concentration. Sensorgrams were obtained from an Octet RED96 using an antigen titration series of six concentrations (400-12.5 nM by twofold dilution for trimer, 25-0.78 nM by twofold dilution for FR, and 10-0.31 nM for multilayered E2p and I3-01v9) and quantitation (AHQ) biosensors. The average peak signals (nm) at the highest antigen concentration are listed in the matrix with the standard deviation (SD) shown in Fig. S6f. A higher color intensity indicates greater binding signal. Source data are provided as a Source Data file.

Previously, we demonstrated the use of a pan-reactive T cell epitope both as a linker and as a built-in T-cell help in an HIV-1 Env-I3-01 NP construct^55^, suggesting that additional structural and functional components can be incorporated into such large 60-meric NPs. Here, we sought to reengineer the E2p and I3-01 NPs by fusing a dimeric locking domain (LD) to the C-terminus of an NP subunit and then a T-helper epitope to the C terminus of an LD (**Fig. 4d**). We hypothesized that each LD dimer can stabilize a non-covalent NP-forming interface from inside, and the T-cell epitopes can form a hydrophobic core at the center of a fully assembled NP. To test this hypothesis, we manually inspected 815 homodimers in the PDB and selected nine LDs of 100 residues or less (**Fig. S6a**). Based on structural compatibility, LDs 1-7 were tested for E2p, and five LDs (4-5 and 7-9) were tested for I3-01, all displaying GPΔmuc-WL^2^P^2^. Following transient expression in 100-ml ExpiCHO cells and mAb100 purification, 12 LD-containing NP samples were characterized by SEC (**Fig. 4e**). Notably, LD4 and LD7 increased the NP peak (UV_280_ value) by 5- and 2.5-fold for E2p and I3-01, respectively, with substantially improved NP purity. The further incorporation of a T-cell epitope, PADRE, did not alter E2p properties, but negatively impacted I3-01 (**Fig. 4f**). An I3-01 variant, termed I3-01v9 (or 1VLW-v9 in the previous study^55^), was found to retain the NP yield and purity (**Fig. 4f**). Seven GP-NP samples, with three variants for each 60-meric NP, were further analyzed by BN-PAGE (**Fig. 4g**). The FR and two E2p variants displayed a single high m.w. band corresponding to well-formed NPs, whereas the wildtype E2p and all three I3-01 samples showed additional low m.w. bands at 232-440 kD on the BN gel, indicating unassembled GP-NP species. Lastly, the mAb100/SEC-purified FR, E2p-LD4-PADRE (E2p-L4P), and I3-01v9-LD7-PADRE (I3-01v9-L7P) NPs that present the GPΔmuc-WL^2^P^2^ trimer were analyzed by negative-stain EM (**Fig. 4h**). In addition to well-formed particles in all three samples, an array of GPΔmuc spikes could be clearly recognized on the surface of FR and E2p-L4P NPs.

Antigenicity of the three GPΔmuc-WL^2^P^2^-presenting NPs was assessed by ELISA using the same panel of 10 antibodies (**Fig. 4i, Fig. S6b** and **S6c**). Compared with the WL^2^P^2^ trimer, the three NPs exhibited an epitope-specific binding pattern. Overall, multivalent display improved antibody recognition of the RBS and glycan cap in GP1, but reduced binding for bNAbs that target the base and IFL at the GP1/GP2 interface (e.g., CA45) and the GP2 stalk (e.g., BDBV223). This finding raised concerns that some conserved bNAb epitopes on the NP-displayed GP trimers may not be as accessible as on the soluble GP trimers. Two BLI experiments, with a total of three replicates for the highest antigen concentration, were performed to further characterize the effect of multivalent display on the antibody recognition of various GP epitopes (**Fig. 4j, Fig. S6d**-**S6f**). Using comparable GP molar concentrations, the three NPs showed higher binding signals than the soluble trimer, with the most notable differences for NAbs mAb114 and mAb100 and a non-NAb c13C6. Based on these results, the FR, E2p-L4P, and I3-01v9-L7P NPs that present the redesigned GPΔmuc-WL^2^P^2^ trimer were selected for animal immunization.

Our results indicate that EBOV GP can be displayed on self-assembling NPs through gene fusion, which requires the optimization of both GP and NP. In addition to GPΔmuc, GP_ECTO_ was also tested but found unsuitable for NP display, as confirmed by the EM analysis of a GP_ECTO_-10GS-FR construct (**Fig. S6h**). The multilayered NP design exploits the inner space of large, cage-like NPs to increase their stability and deliver additional T-cell signals within a single-component system, representing a distinct strategy compared with the two-component NP design^89,90^.

### Immunogenicity of EBOV GP trimers and NPs in BALB/c mice

Following the protocol in our previous HIV-1 and HCV studies^55,65^, we immunized BALB/c mice to assess the immunogenicity of six representative constructs, including three EBOV GP/GPΔmuc trimers and three NPs (**Fig. 5a**). A GP_ECTO_-foldon trimer was included as a soluble version of the wildtype GP. Mice in the GPΔmuc-WL^2^P^2^-10GS-I3-01v9-L7P group were immunized with 20 μg mAb100-purified protein instead of 50 μg mAb100/SEC-purified protein due to the low yield of this NP. We first determined the GP-specific plasma antibody response in the ELISA using GPΔmuc-WL^2^P^2^-1TD0 as a probe, which utilized trimerization motif 1TD0 (PDB ID: 1TD0) (**Fig. 5b, Fig. S7a**-**S7c**). The two GPΔmuc groups significantly (*P* < 0.0064) outperformed the GP_ECTO_ group throughout immunization (**Fig. 5b**, top), suggesting that the MLD can shield GP from antibody recognition. However, little difference was found between the two GPΔmuc groups, with WL^2^P^2^ showing a slightly higher average EC_50_ titer at w2 and w5 that was reversed at later time points. Compared with the WL^2^P^2^ trimer group, all NP groups showed lower EC_50_ titers except for the E2p-L4P NP group, which yielded a modest *P* value of 0.0381 at w2 (**Fig. 5b**, bottom). In our recent HCV study, two NPs elicited higher antibody titers than the soluble E2 core at w2 (*P* < 0.0001) and w5 (*P* ≤ 0.0223)^65^. The stark difference between these two studies suggests that antibody titers in response to such NP vaccines may be influenced by antigen size, structure, and epitope distribution. NP display may occlude antibody access to the base and stalk epitopes, which are targets of many bNAbs^9^. This result may also be attributed to other factors, such as dosage, as noted in our recent study^65^. The NP carrier accounts for 21-33% of the total mass of an NP vaccine, and the same antigen dose (50 μg) has been used for all groups except the I3-01v9-L7P NP group. Thus, mice in the NP groups would receive less GP antigen than mice in the trimer-only group.

**Fig. 5.**
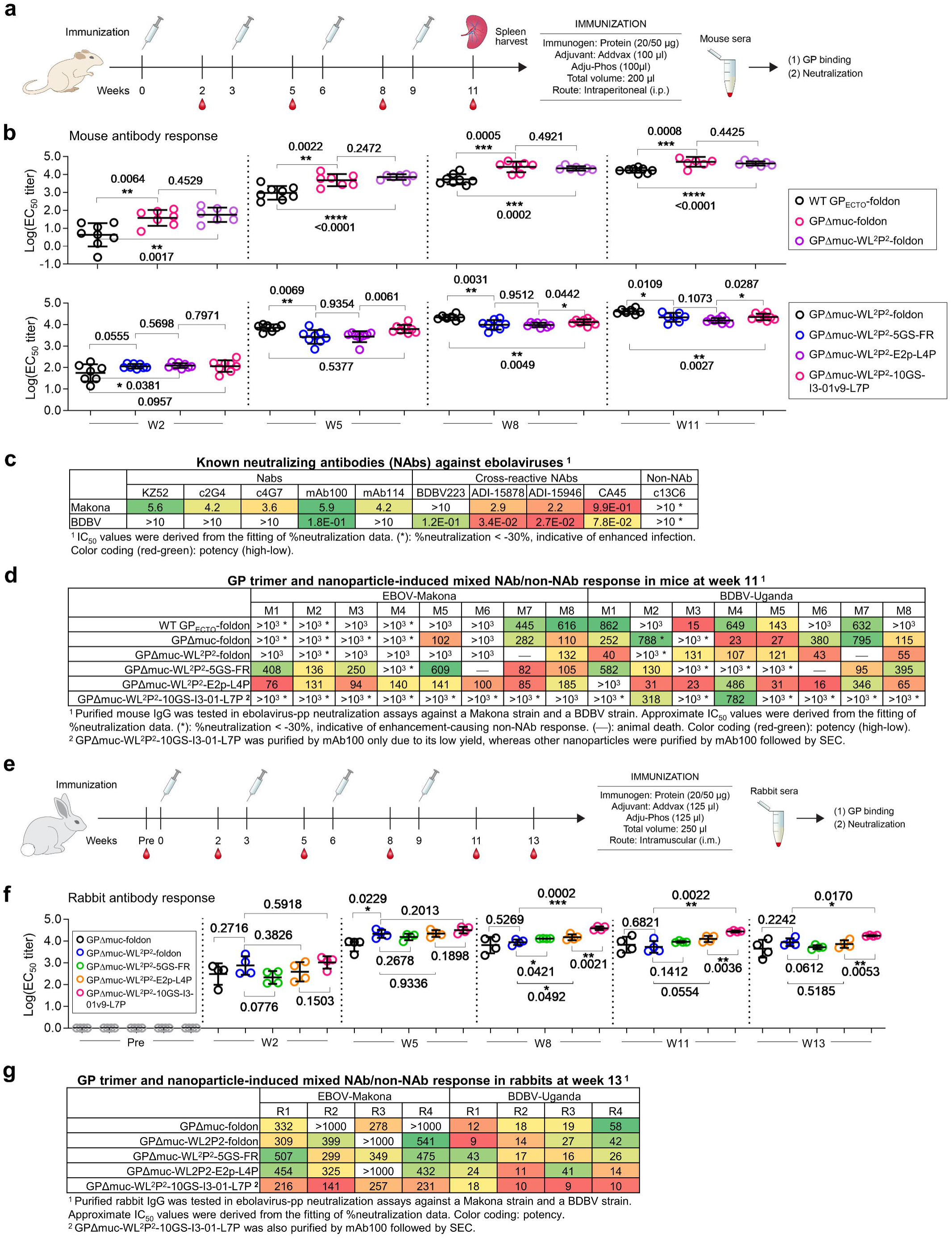
Immunogenicity of EBOV GPΔMuc trimers and nanoparticles in mice. **(a)** Schematic representation of the mouse immunization protocol. **(b)** Longitudinal analysis of GP-specific antibody titers in immunized mouse sera (n=8) at w2, w5, w8, and w11. Top panel: EC_50_ titers (fold of dilution) calculated from ELISA binding of mouse sera from three GP trimer groups to the coating antigen, GPΔMuc-WL^2^P^2^. Bottom panel: EC_50_ titers calculated from ELISA binding of mouse sera from three NP groups to the coating antigen, GPΔMuc-WL^2^P^2^, with the GPΔMuc-WL^2^P^2^-foldon group included for comparison. Mean values and standard deviation (SD) are shown as black lines. *P*-values were determined by an unpaired two-tailed *t* test in GraphPad Prism 8.4.3 and are labeled on the plots. The asterisk symbol (*) indicates the level of statistical significance: *, *P* < 0.05; **, *P* < 0.01; ***, *P* < 0.001; ****, *p* < 0.0001. **(c)** Ebolavirus-pp (EBOV-Makona and BDBV-Uganda) neutralization by 10 antibodies in 293 T cells. The neutralization was performed in duplicates. The half maximal inhibitory concentration (IC_50_) values are listed with the enhanced pseudovirus infection (<-30%) indicated by an asterisk (*). **(d)** Ebolavirus-pp (EBOV-Makona and BDBV-Uganda) neutralization by purified mouse IgGs from six vaccine groups in 293 T cells. The neutralization was performed without duplicates due to limited sample availability. **(e)** Schematic representation of the rabbit immunization protocol. **(f)** Longitudinal analysis of GP-specific antibody titers in immunized rabbit sera (n=4) at weeks 0, 2, 5, 8, 11, and 13. Mean values and standard deviation (SD) are shown as black lines. **(g)** Ebolavirus-pp (EBOV-Makona and BDBV-Uganda) neutralization by purified rabbit IgGs from five vaccine groups in 293 T cells. The neutralization was performed in duplicates. In (d) and (g), due to the presence of enhancement-causing non-NAbs in the purified IGs, approximate IC_50_ values derived from the fitting of % neutralization data are listed for comparison, with the enhanced pseudovirus infection (< −30%) indicated by an asterisk (*). Source data are provided as a Source Data file.

Before analyzing the mouse plasma NAb response, we validated the pseudoparticle (pp) neutralization assay^12^ by testing 10 antibodies against two ebolavirus strains, EBOV-Makona and BDBV-Uganda, in 293 T and TZM-bl^72^ cells (**Fig. 5c, Fig. S7d**-**S7f**). mAb114^12^ and early EBOV NAbs KZ52^76^, c2G4, and c4G7^10^ only neutralized EBOV-Makona, whereas mAb100^12^ and four bNAbs, except BDBV223^9^, blocked both ebolavirus-pps. ADI-15946 was the most potent bNAb, as indicated by the half maximal inhibitory concentration (IC_50_). Non-NAb c13C6, which is part of the ZMapp cocktail^10^ and binds the glycan cap^45^, enhanced ebolavirus-pp infection of both cell types. When tested against pseudoparticles bearing the murine leukemia virus Env, MLV-pps, all antibodies were non-reactive, except for c13C6, which enhanced MLV-pp infection in 293 T cells (**Fig. S7g**). Overall, the enhancement observed for non-NAb c13C6 in the pseudovirus assays appeared to be consistent with ADE observed for human mAbs targeting the same epitope^25^.

We next performed neutralization assays using purified mouse immunoglobulin G (IgG) from the last time point, w11 (**Fig. 5d, Fig. S7h** and **S7i**). Two distinct types of antibody response were observed: NAbs and non-NAbs that enhanced ebolavirus-pp infection. Among the three GP trimers, GP_ECTO_ elicited a moderate NAb response with signs of enhancement noted for three mice, whereas an increase in both types of antibody response was observed for GPΔmuc, suggesting that the removal of MLD exposes both NAb epitopes and the glycan cap, which is a main target for ADE-causing human mAbs^25^. The stalk/HR1_C_ mutation WL^2^P^2^ appeared to have largely reversed the enhancement caused by MLD removal. Among the three NPs, E2p-L4P elicited primarily NAb responses that blocked both ebolavirus-pps, whereas enhanced pseudoviral infection was observed for one mouse in the FR group and for all mice in the I3-01v9-L7P group. Because only non-NAb c13C6 and not any of the (b)NAbs reacted with MLV-pps (**Fig. S7g**), here we utilized the MLV-pp assay to gauge glycan cap-directed non-NAb responses induced by different vaccine constructs (**Fig. S7j**). Indeed, the patterns of enhanced MLV-pp infection correlated nicely with the patterns of enhanced ebolavirus-pp infection (**Fig. S7h-S7j**). In the MLV-pp assay, E2p-L4P NP induced a minimum enhancement-causing non-NAb response at a similar level to GP_ECTO_, in which MLD shields the glycan cap and other GP epitopes from the humoral response.

Our mouse study thus revealed some salient features of the GP-specific antibody response in the context of various GP forms and NP carriers. The c13C6-like non-NAbs that bind the glycan cap and cross-react with small secreted GP (ssGP)^45^ need to be minimized in vaccine design. The high level of enhancement-causing non-NAb responses observed for GPΔmuc and I3-01v9-L7P may be explained by less trimeric GP and unassembled GP-NP species, respectively. Nonetheless, a multilayered E2p NP displaying 20 GPΔmuc trimers elicited a robust bNAb response.

### Immunogenicity of EBOV GP trimers and NPs in rabbits

Following a similar protocol, we assessed two GPΔmuc trimers, wildtype and WL^2^P^2^, and three NPs presenting the WL^2^P^2^ trimer in rabbits (**Fig. 5e**). Rabbit sera collected at six timepoints during immunization were analyzed by ELISA using the same trimer probe (**Fig. 5f, Fig. S8a** and **S8b**). Notably, rabbits in the I3-01v9-L7P NP group were immunized with 20 μg of mAb100/SEC-purified material to reduce the enhancement-causing non-NAbs. Between the two trimer groups, WL^2^P^2^ showed higher average EC_50_ titers for all time points except w11, with a modest *P* value of 0.0229 at w5. Among the three NP groups, the I3-01v9 and FR groups yielded the highest and lowest EC_50_ titers, respectively, throughout immunization. A significant difference was found between the I3-01v9-L7P and E2p-L4P groups at w8, w11, and w13, with *P* values in the range of 0.0021 to 0.0053. Compared with the GPΔmuc-WL^2^P^2^ group, the I3-01v9-L7P NP group showed higher EC_50_ titers at all six time points, with significant *P* values at w8, w11, and w13. In contrast, the FR and E2p-L4P groups yielded lower EC_50_ titers than the GPΔmuc-WL^2^P^2^ group at w2 and w5, but this pattern was reversed at w8 and w11 with modest *P* values at w8. However, these two NP groups ended with lower EC_50_ titers than the trimer group at the last time point, w13.

We then performed ebolavirus-pp and MLV-pp assays using purified rabbit IgG from w11 (**Fig. 5g, Fig. S8c**). At this time point, all vaccine groups showed NAb responses with no sign of enhanced pseudovirus infection, in contrast to the pattern of mixed antibody responses in mice (**Fig. 5d**). Notably, the I3-01v9-L7P NP group yielded higher average IC_50_ titers than the other groups, 211.3 μg/ml and 11.72 μg/ml for EBOV-Makona and BDBV-Uganda, respectively, supporting the notion that unassembled GP-NP species and not the I3-01v9 NP carrier were responsible for eliciting enhancement-causing non-NAbs in mice. All vaccine groups showed no sign of enhanced MLV-pp infection at w11 (**Fig. S8c**). Therefore, enhancement-causing non-NAbs appeared to be absent in rabbit plasma toward the end of immunization. We next analyzed rabbit IgG from earlier time points day 0 (Pre), w2, w5, and w8 (**Fig. S8d-S8g**), which revealed a unique temporal pattern of an increasing NAb response tailing a transient enhancement-causing non-NAb response. Specifically, enhanced Makona-pp infection was observed for the two trimer groups, FR group, and two multilayered NP groups at w2, w5, and w8, which then disappeared at w5, w8, and w11, respectively. Our longitudinal analysis suggests that vaccine-induced enhancement-causing non-NAbs may shift epitopes and gain neutralizing activity through gene conversion, a mechanism employed by the rabbit immune system to rapidly develop functional antibodies^91^.

### B cell response profiles associated with EBOV GP trimers and NPs

Previously, we combined antigen-specific B cell sorting and NGS to obtain a quantitative readout of the B-cell response induced by an HCV E2 core and its E2p NP^65^. A more diverse usage of heavy-chain variable genes (V_H_), a higher degree of V_H_ mutations, and a broader range of heavy chain complementarity determining region 3 (HCDR3) length were observed for E2p^65^. In this study, we applied the same strategy to obtain GP-specific B cell profiles (**Fig. 6a**). Using an Avi-tagged GPΔmuc-WL^2^P^2^-1TD0 probe (**Fig. S9a**), we sorted GP-specific splenic B cells from 25 mice (**Fig. S9b**), which were sequenced on Ion GeneStudio S5. The NGS data were analyzed using a mouse Antibodyomics pipeline^92^ (**Fig. S9c**), with quantitative B cell profiles derived for different vaccine groups (**Fig. 6b, Fig. S9d-S9f**). We mainly focused on the GPΔmuc-WL^2^P^2^-foldon group and multilayered E2p group to compare B-cell responses to GPΔmuc in the trimeric versus NP forms (**Fig. 6b**). In terms of germline gene usage, similar patterns were observed for V_H_ and V_K_ genes (**Fig. 6b**, panels 1 and 2). The redesigned GPΔmuc trimer activated more V_H_/V_L_ genes (9.4/9.4) than its NP form (6/7), with *P* values of 0.0163 and 0.0076 for V_H_ and V_L_ genes, respectively. In contrast, the E2p NP decorated with 60 HCV E2 cores activated more V_H_ but not V_L_ genes than the E2 core^65^. In terms of somatic hypermutation (SHM), no significant difference was found between the two groups (**Fig. 6b**, panel 3). However, we observed a visible shift in the SHM distribution for the E2p-L4P NP group, which showed higher germline V_H_/V_K_ divergence (on average 6.4%/2.9%) than the trimer group (on average 5.3%/2.6%). In the HCDR3 analysis, both average loop length and the RMS fluctuation (RMSF) of loop length were calculated (**Fig. 6b**, panel 4). Unlike in the HCV study, in which the RMSF of HCDR3 length yielded a *P* <0.0001 between the E2 core and E2p NP groups^65^, no significant difference was found between the EBOV GPΔmuc and E2p NP groups. Overall, EBOV and HCV NPs exhibited distinct patterns of the B-cell response with respect to their individual antigens. Notably, there were no apparent correlations between B-cell profiles and vaccine-induced NAb and enhancement-causing non-NAb responses. Our results thus suggest that antigen size, structure, glycosylation, and epitope distribution, other than the NP carrier, may contribute critically to the NP vaccine-induced B-cell response.

**Fig. 6.**
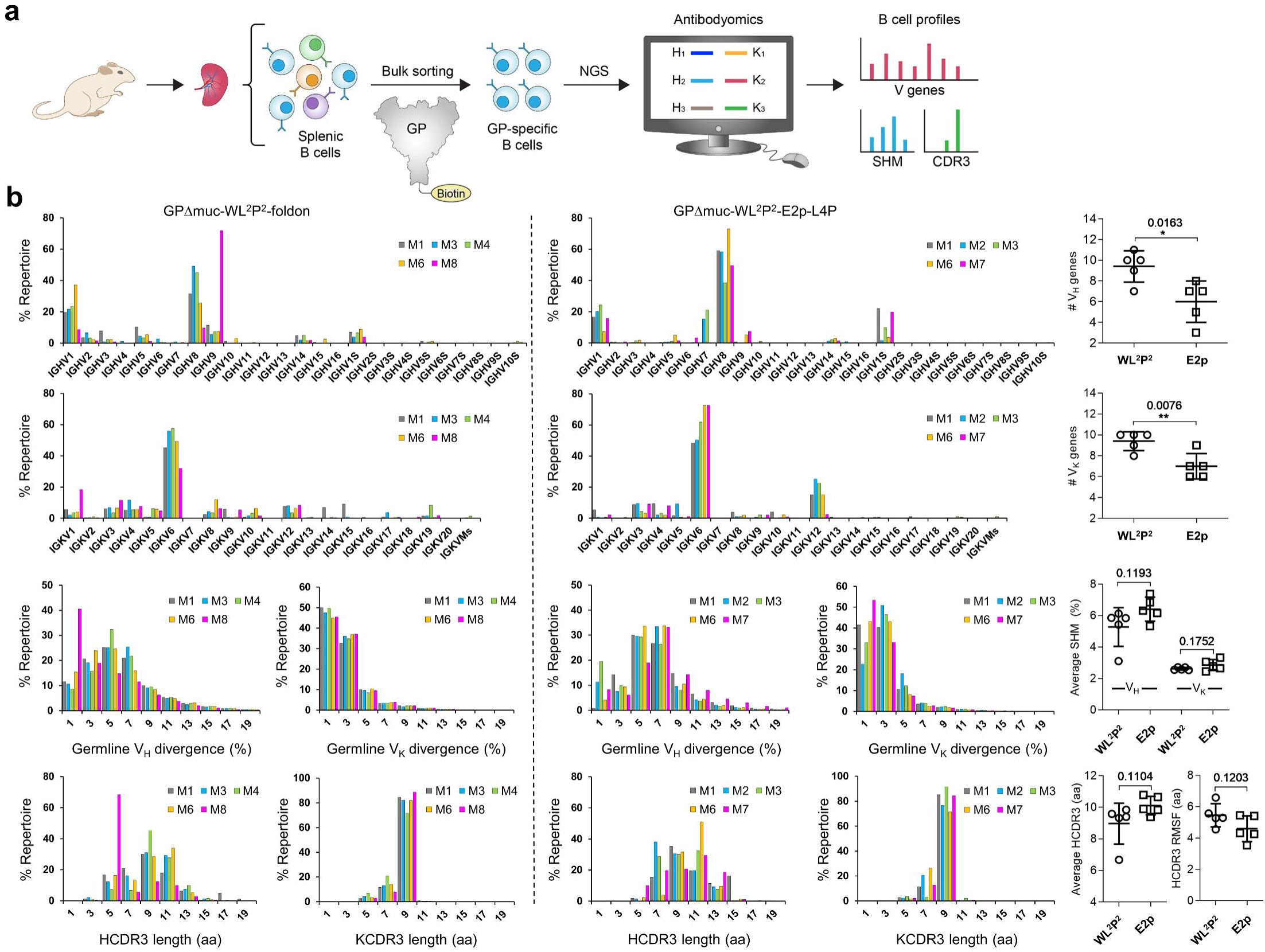
Quantitative assessment of GP-specific B-cell response. **(a)** Schematic representation of the strategy to analyze the GP-specific murine B-cell response that combines the antigen-specific bulk sorting of splenic B cells with next-generation sequencing (NGS) and antibodyomics analysis. **(b)** B cell profiles for the GPΔMuc-WL^2^P^2^-foldon (left) and GPΔMuc-WL^2^P^2^-E2p-L4P groups (right) and statistical comparison of key parameters derived from the profile analysis (far right). For each vaccine group (n=8), 5 mice were selected for NGS analysis. Panel 1/2: Distribution of germline V_H_/V_K_ genes and statistical comparison of the number of activated V_H_/V_K_ genes (≥ 1% of the total population). Panel 3: Distribution of V_H_/V_K_ somatic hypermutation (SHM) with percent (%) mutation calculated at the nucleotide level, and statistical comparison of the average V_H_/V_K_ SHM rate. Panel 4: Distribution of H/KCDR3 loop length and statistical comparison of two parameters, HCDR3 loop length and root-mean-square fluctuation (RMSF) of HCDR3 loop length. The RMSF value is used as an indicator of how much HCDR3 loop length varies within the EBOV GP-specific antibodies from each animal. In statistical comparison (far right), mean value and standard deviation (SD) are shown as black lines. *P*-values were determined by an unpaired two-tailed *t* test in GraphPad Prism 8.4.3 and are labeled on the plots. The asterisk symbol (*) indicates the level of statistical significance: *, *P* < 0.05; **, *P* < 0.01; ***, *P* < 0.001. Source data are provided as a Source Data file.

## Discussion

With a mortality rate of up to 90%, EBOV has caused significant humanitarian crises, calling for action across many sectors to develop effective interventions^3^. ZMapp^10,11^ established the use of NAbs as a treatment for patients with Ebola virus disease (EVD) and propelled a community-wide campaign to identify NAbs and bNAbs^8,9,30,48^. Several vaccine candidates have been tested in human trials^63,93^. Of these, rVSV-ZEBOV demonstrated an efficacy of 100% in an open-label, cluster-randomized ring trial^16^ and was recently approved for human use. In a Phase 2 placebo-controlled trial of two vectored vaccines (including rVSV-ZEBOV), the antibody titer remained similar between the vaccine and placebo groups at one week post-vaccination and peaked after one month^15^. A recent analysis of human B cell responses to EBOV infection revealed hurdles in the GP-specific NAb response^28,29^. NAbs from vaccinated humans showed low levels of SHM^94^, suggesting a suboptimal B cell response elicited by an otherwise effective vectored vaccine^95^. The immune correlates of vaccine protection may not be universal and are likely determined by vaccine platforms, in addition to other factors^96^. As most EBOV vaccines are based on viral vectors^63,93^, protein subunit vaccines remain a promising approach to combat this deadly virus.

Here, we approached EBOV vaccine design with an antibody-based strategy focusing on GP optimization and multivalent display. We applied a similar strategy to develop HIV-1 and HCV vaccine candidates for in vitro and in vivo characterization^55,56,64,65^, which provided a context for interpreting the findings for EBOV. Previously, we identified an HR1_N_ bend as the cause of HIV-1 Env metastability^55,56^ and optimized an HCV E2 core^65^. EBOV GP belongs to the class-I fusion protein family^53,54^ and is inherently metastable, as is HIV-1 Env. In this study, we probed the cause of EBOV GP metastability by testing various designs that target the HR2 stalk, HR1_C_ bend, and GP/GP interface (via inter-protomer disulfide bonds). The detailed characterization revealed a stabilizing effect of the W615L mutation and stalk extension, in addition to the unexpected sensitivity of HR1_C_ (equivalent to HIV-1 HR1_N_) to specific proline mutations, which increased the trimer yield but caused complex unfolding behaviors in DSC. Because this pattern was not reported for EBOV-Makona GPΔmuc that contained the T577P mutation^84^, GP metastability may thus be a strain-specific feature and warrant further investigation. Multivalent NP display proved to be challenging for EBOV GP because of its tendency toward dissociation. Although two-component NPs^89,90^ and the SpyTag/SpyCatcher system^97,98^ have been used to develop VLP-type vaccines, their inherent complexity in production, assembly, and purification, structural instability in vivo, and off-target response may dampen their potential as human vaccines. Here, we designed single-component, multilayered, and self-assembling protein NPs based on E2p and I3-01v9. Encoded within a single plasmid, such NP vaccines can be readily produced in good manufacturing practice (GMP)-compatible CHO cells followed by IAC and SEC purification, providing a simple and robust manufacturing process. Our immunogenicity studies in mice and rabbits revealed some salient features of GP that need to be addressed in EBOV vaccine development, regardless of the delivery platform. The choice of GP_ECTO_ or GPΔmuc as a vaccine antigen may lead to antibody responses that target different GP epitopes. Antibody access to GP epitopes at the IFL, base, and HR2 stalk may differ in the trimeric and NP forms. In animal immunization, we observed an ADE-like non-NAb response, which may be associated with non-trimeric GP and unassembled GP-NP species. Antibody isolation, structural epitope mapping, and live EBOV neutralization assays may be required to determine the biological relevance of these findings. Furthermore, EBOV challenge in rodents may help determine protective NAb titers elicited by GP-presenting NPs with respect to recombinant VLPs^60–62^. Nonetheless, the E2p-L4P NP elicited a minimum non-NAb response in mice and the highly purified I3-01v9-L7P NP induced the strongest NAb response in rabbits, providing two promising constructs for further optimization and in vivo evaluation.

Having assessed various GP trimer and NP constructs, future investigation may be directed toward assessing other GPΔmuc designs, such as GPΔmuc-WL^2^ and GPΔmuc-SS2, as well as their NPs, to further improve NAb responses and reduce glycan cap-directed non-NAb responses. The structural characterization of NAbs and non-NAbs isolated from immunized animals will provide critical insights into epitope recognition and guide future vaccine design. The strategy described in this study may find applications in vaccine development for other filoviruses.

## Methods

### Design, expression, and purification of EBOV GPΔmuc and GPΔmuc-presenting NPs

The glycoprotein sequence of Zaire EBOV (Mayinga-76 strain) with a T42A substitution was used to design all GP constructs in this study (UniProt ID: Q05320), with the primers summarized in **Table S1**. Logo analysis of EBOV and MARV GP sequences was performed using the WebLoGo v2.8 software to facilitate the design of the W615L mutation. Structural modeling was performed using the UCSF Chimera v1.13 software to facilitate the design of HR1_C_-proline and inter-protomer disulfide bond mutations. Wildtype and redesigned GPΔmuc constructs were transiently expressed in HEK293F cells (Thermo Fisher) for biochemical, biophysical, and antigenic analyses. Briefly, HEK293F cells were thawed and incubated with FreeStyle^TM^ 293 Expression Medium (Life Technologies, CA) in a shaker incubator at 37 °C at 135 rpm with 8% CO_2_. When the cells reached a density of 2.0 × 10^6^/ml, expression medium was added to reduce cell density to 1.0 × 10^6^ ml^−1^ for transfection with polyethyleneimine (PEI) (Polysciences, Inc). Next, 900 μg of plasmid in 25 ml of Opti-MEM transfection medium (Life Technologies, CA) was mixed with 5 ml of PEI-MAX (1.0 mg/ml) in 25 ml of Opti-MEM. After 30 min of incubation, the DNA-PEI-MAX complex was added to 1L 293 F cells. Culture supernatants were harvested 5 days after transfection, clarified by centrifugation at 1126 ×g for 22 min, and filtered using a 0.45 μm filter (Thermo Scientific). GPΔmuc proteins were extracted from the supernatants using an mAb114 antibody column or mAb100 antibody column. Bound proteins were eluted three times, each with 5 ml of 0.2 M glycine (pH 2.2) and neutralized with 0.5 ml of Tris-Base (pH 9.0), and buffer-exchanged into phosphate-buffered saline (PBS; pH 7.2). Proteins were further purified by SEC on a Superdex 200 Increase 10/300 GL column or HiLoad Superdex 200 16/600 column (GE Healthcare). GPΔmuc-presenting NPs were produced in ExpiCHO cells (Thermo Fisher). Briefly, ExpiCHO cells were thawed and incubated with ExpiCHO^TM^ Expression Medium (Thermo Fisher) in a shaker incubator at 37 °C at 135 rpm with 8% CO_2_. When the cells reached a density of 10×10^6^ ml^−1^, ExpiCHO^TM^ Expression Medium was added to reduce cell density to 6×10^6^ ml^−1^ for transfection. The ExpiFectamine^TM^ CHO/plasmid DNA complexes were prepared for 100-ml transfection in ExpiCHO cells following the manufacturer’s instructions. For these NP constructs, 100 µg of plasmid and 320 μl of ExpiFectamine^TM^ CHO reagent were mixed in 7.7 ml of cold OptiPRO™ medium (Thermo Fisher). After the first feed on day 1, ExpiCHO cells were cultured in a shaker incubator at 33 °C at 115 rpm with 8% CO_2_ according to the Max Titer protocol with an additional feed on day 5 (Thermo Fisher). Culture supernatants were harvested 13 to 14 days after transfection, clarified by centrifugation at 3724 ×g for 25 min, and filtered using a 0.45 μm filter (Thermo Fisher). The mAb100 antibody column was used to extract nanoparticles from the supernatants, followed by SEC on a Superose 6 10/300 GL column. All SEC data were collected using the Unicorn 7.5 software. For GPΔmuc and GPΔmuc-presenting NPs, the concentration was determined using UV_280_ absorbance with theoretical extinction coefficients.

### Blue native polyacrylamide gel electrophoresis

EBOV GPΔmuc and GPΔmuc-presenting NPs were analyzed by blue native polyacrylamide gel electrophoresis (BN*-*PAGE) and stained with Coomassie blue. The proteins were mixed with sample buffer and G250 loading dye and added to a 4-12% Bis-Tris NativePAGE^TM^ gel (Life Technologies). BN-PAGE gels were run for 2 to 2.5 h at 150 V using NativePAGE^TM^ running buffer (Life Technologies) according to the manufacturer’s instructions. BN-PAGE images were collected using the Image Lab v6.0 software.

### Enzyme-linked immunosorbent assay

Each well of a Costar^TM^ 96-well assay plate (Corning) was first coated with 50 µl of PBS containing 0.2 μg of appropriate antigens. The plates were incubated overnight at 4 °C, and then washed five times with wash buffer containing PBS and 0.05% (v/v) Tween 20. Each well was then coated with 150 µl of blocking buffer consisting of PBS, 40 mg ml^−1^ blotting-grade blocker (Bio-Rad), and 5% (v/v) FBS. The plates were incubated with blocking buffer for 1 h at room temperature, and then washed five times with wash buffer. For antigen binding, antibodies were diluted in blocking buffer to a maximum concentration of 10 μg ml^−1^ followed by a 10-fold dilution series. For each antibody dilution, a total of 50 μl volume was added to the appropriate wells. For animal sample analysis, plasma was diluted 10-fold for mouse and 50-fold for rabbit in blocking buffer and subjected to a 10-fold dilution series. For each sample dilution, a total of 50 μl volume was added to the wells. Each plate was incubated for 1 h at room temperature, and then washed five times with PBS containing 0.05% Tween 20. For antibody binding, a 1:5000 dilution of goat anti-human IgG antibody (Jackson ImmunoResearch Laboratories, Inc), or for animal sample analysis, a 1:2000 dilution of horseradish peroxidase (HRP)-labeled goat anti-mouse or anti-rabbit IgG antibody (Jackson ImmunoResearch Laboratories), was then made in wash buffer (PBS containing 0.05% Tween 20), with 50 μl of this diluted secondary antibody added to each well. The plates were incubated with the secondary antibody for 1 h at room temperature, and then washed five times with PBS containing 0.05% Tween 20. Finally, the wells were developed with 50 μl of TMB (Life Sciences) for 3-5 min before stopping the reaction with 50 μl of 2 N sulfuric acid. The resulting readouts were measured on a plate reader (PerkinElmer) at a wavelength of 450 nm and collected using the PerkinElmer 2030 v4.0 software. Notably, the week 2 plasma binding did not reach the plateau (or saturation) to allow for the accurate determination of EC_50_ titers. Nonetheless, the EC_50_ values were calculated in GraphPad Prism 8.4.3 and used as a quantitative measure of antibody titers to facilitate comparisons of different vaccine groups at week 2.

### Bio-layer interferometry

The kinetics of GPΔmuc and GPΔmuc-presenting NP binding to a panel of 10 antibodies was measured using an Octet RED96 instrument (FortéBio, Pall Life Sciences). All assays were performed with agitation set to 1000 rpm in FortéBio 1× kinetic buffer. The final volume for all solutions was 200 μl per well. Assays were performed at 30 °C in solid black 96-well plates (Geiger Bio-One). Antibody (5 μg ml^−1^) in 1× kinetic buffer was loaded onto the surface of anti-human Fc Capture Biosensors (AHC) for GPΔmuc and of anti-human Fc Quantitation Biosensors (AHQ) for NPs for 300 s. A 60 s biosensor baseline step was applied prior to analyzing association of the antibody on the biosensor to the antigen in solution for 200 s. A two-fold concentration gradient of antigen, starting at 400 nM for GPΔmuc trimers, 25 nM for FR NPs, and 10 for E2p/I3-01v9 NPs, was used in a titration series of six. Dissociation of the interaction was followed for 300 s. The correction of baseline drift was performed by subtracting the mean value of shifts recorded for a sensor loaded with antibody but not incubated with antigen and for a sensor without antibody but incubated with antigen. The Octet data were processed by FortéBio’s data acquisition software v8.2. For GPΔmuc trimers, experimental data were fitted with the binding equations describing a 2:1 interaction to achieve the optimal fitting and determine the *K_D_* values. For GP-presenting nanoparticles, two BLI experiments, one testing six antigen concentrations and the other testing the highest antigen concentration in duplicates, were performed. Binding signals at the highest antigen concentration (mean and standard deviation calculated from three replicates) were used to quantify the effect of multivalent NP display on GP-antibody interactions. Notably, the GPΔmuc-WL^2^P^2^-foldon trimer was also measured using AHQ biosensors to facilitate comparisons with three nanoparticles that present the GPΔmuc-WL^2^P^2^ trimer multivalently.

### Differential scanning calorimetry

Thermal melting curves of wildtype and redesigned GPΔmuc trimers were obtained with a MicroCal VP-Capillary calorimeter (Malvern). The purified GPΔmuc protein produced from 293F cells was buffer exchanged into 1×PBS and concentrated to 27–50 μM before analysis by the instrument. Melting was probed at a scan rate of 90 °C·h^−1^ from 25 °C to 110 °C. Data processing, including buffer correction, normalization, and baseline subtraction, was conducted using the standardized protocol from Origin 7.0 software.

### Protein production, crystallization, and data collection

Two *Zaire* EBOV GPΔmuc-foldon constructs, one with the W615L mutation and the L extension (to residue 637) and the other with an additional T577P mutation, were expressed in HEK293S cells. The expressed GP was purified using an mAB100 antibody column followed by SEC on a HiLoad Superdex 200 16/600 column (GE Healthcare). PBS (pH 7.2) was used as the gel filtration buffer during the purification process. The freshly purified GP protein was used for crystallization experiments using the sitting drop vapor diffusion method on our automated CrystalMation^TM^ robotic system (Rigaku) at both 4 °C and 20 °C at The Scripps Research Institute (TSRI)^99^. EBOV GP was concentrated to ∼10 mg/ml in 50 mM Tris-HCl, pH 8.0. The reservoir solution contained 12% (w/v) PEG 6000 and 0.1 M sodium citrate, pH 4.5. Diffraction-quality crystals were obtained after 2 weeks at 20 °C. The EBOV GP crystals were cryoprotected with 25% glycerol, mounted in a nylon loop and flash cooled in liquid nitrogen. Diffraction data were collected for crystals of GPΔmuc-WL^2^-foldon and GPΔmuc-WL^2^P^2^-foldon at Advanced Photon Source (APS) beamline 23ID-D and Stanford Synchrotron Radiation Light-source (SSRL) beamline 12-2, at 2.3 Å and 3.2 Å resolution, respectively. The diffraction data sets were processed with HKL-2000^100^. The crystal data were indexed in rhombohedral R32 and tetragonal P321 space groups with cell dimensions of GPΔmuc-WL^2^-foldon *a* = *b* = 114.58 Å and *c* = 312.38 Å and GPΔmuc-WL^2^P^2^-foldon *a* = *b* = 114.06 Å and *c* = 136.22 Å, respectively (**Table S2**). The overall completeness of the two datasets was 96.4% and 99.9%.

### Structure determination and refinement

The structures of EBOV GP were determined by molecular replacement (MR) using Phaser ^101^ from the CCP4i suite ^102^ with the coordinates of *Zaire* Ebola GP (PDB ID: 5JQ3) and the program MOLREP ^103^. The polypeptide chains were manually adjusted into electron density using Coot ^104^, refined with Refmac 5.8^105^, and validated using MolProbity^106^. The final R_cryst_ and R_free_ values for the refined structures are 19.6% and 22.9%, and 28.6% and 33.6%, for GPΔmuc-WL^2^-foldon and GPΔmuc-WL^2^P^2^-foldon, respectively. The data processing and refinement statistics are compiled in **Table S2**. Structural images shown in Fig. 3 and Supplementary Figs. 3-5 were generated using the PyMOL v2.3.4 software.

### Electron microscopy (EM) assessment of nanoparticle constructs

The initial EM analysis of EBOV GPΔMuc NPs was conducted at the Core Microscopy Facility at The Scripps Research Institute. Briefly, nanoparticle samples were prepared at a concentration of 0.01 mg/ml. Carbon-coated copper grids (400 mesh) were glow-discharged, and 8 µl of each sample was adsorbed for 2 min. Excess sample was wicked away and grids were negatively stained with 2% uranyl formate for 2 min. Excess stain was wicked away, and the grids were allowed to dry. Samples were analyzed at 80 kV with a Talos L120C transmission electron microscope (Thermo Fisher), and images were acquired with a CETA 16M CMOS camera. Further EM analysis was conducted at the Hazen facility at The Scripps Research Institute. Nanoparticle samples were diluted to ∼0.02 mg/ml and added onto carbon-coated copper 400 mesh grids (Electron Microscopy Sciences) that had been plasma cleaned for 10 s with Ar/O2. After blotting to remove excess sample, grids were stained with 3 µl of 2% (w/v) uranyl formate for 60 s and blotted again to remove excess stain. Negative stain images were collected on a 120 KeV Tecnai Spirit equipped with an Eagle 4K charge-coupled device (CCD) camera (FEI). Micrographs were collected using Leginon^107^ and processed using cryoSPARC v2^108^. Micrographs were CTF corrected, and particles were picked manually and extracted for two-dimensional classification.

### Animal immunization and sample collection

Similar immunization protocols were reported in our previous HIV-1 and HCV studies^55,65^. Briefly, the Institutional Animal Care and Use Committee (IACUC) guidelines were followed with animal subjects tested in the immunization study. Six-to-eight-week-old female BALB/c mice were purchased from The Jackson Laboratory and housed in ventilated cages in environmentally controlled rooms at The Scripps Research Institute, in compliance with an approved IACUC protocol and AAALAC (Association for Assessment and Accreditation of Laboratory Animal Care) international guidelines. The vivarium was maintained at 22 °C with a 13-hour light/11-hour dark cycle (lights on at 6:00 am and off at 7:00 pm) and 40-50% humidity, which might be reduced to 30%-40% during the winter. The mice were immunized at weeks 0, 3, 6, and 9 with 200 μl of antigen/adjuvant mix containing 50 μg of vaccine antigen and 100 μl of adjuvant, AddaVax or Adju-Phos (InvivoGen), via the intraperitoneal (i.p.) route. Of note, 20 μg instead of 50 μg of mAb100-purified I3-01v9 protein was used in mouse immunization due to its low yield. Blood was collected two weeks after each immunization. All bleeds were performed through the retro-orbital sinus using heparinized capillary tubes into EDTA-coated tubes. Samples were diluted with an equal volume of PBS and then overlaid on 4.5 ml of Ficoll in a 15 ml SepMate^TM^ tube (STEMCELL Technologies) and spun at 300 ×g for 10 min at 20 °C to separate plasma and cells. The plasma was heat inactivated at 56 °C for 30 min, spun at 300 ×g for 10 min, and sterile filtered. The cells were washed once in PBS and then resuspended in 1 ml of ACK Red Blood Cell lysis buffer (Lonza). After washing with PBS, peripheral blood mononuclear cells (PBMCs) were resuspended in 2 ml of Bambanker Freezing Media (Lymphotec). Spleens were also harvested and ground against a 70-μm cell strainer (BD Falcon) to release splenocytes into a cell suspension. Splenocytes were centrifuged, washed in PBS, treated with 5 ml of ACK lysing buffer (Lonza), and frozen with 3ml of Bambanker freezing media. Purified mouse IgGs at w11 were obtained using a 0.2-ml protein G spin kit (Thermo Scientific) following the manufacturer’s instructions and assessed in pseudovirus neutralization assays. Rabbit immunization and blood sampling were performed under a subcontract at ProSci (San Diego, CA) under the IACUC protocol number APF-1A and related amendments (10/01/2018 through 10/01/2021). Five groups of female New Zealand White rabbits, four rabbits per group, were intramuscularly (i.m.) immunized with 50 μg of vaccine antigen formulated in 250 µl of adjuvant, AddaVax or Adju-Phos (InvivoGen), with a total volume of 500 μl, at w0, w3, w6, and w9. Of note, 20 μg of mAb100/SEC-purified I3-01v9 NP was used for rabbit immunization. Blood samples, 20 ml each time, were collected from the auricular artery at day 0 (Pre), w2, w5, w8, and w11. More than 100 ml of blood was taken at w13, via cardiac puncture, for PBMC isolation. Plasma samples were heat inactivated for ELISA binding assays, and purified rabbit IgGs were assessed in pseudovirus neutralization assays.

### Pseudovirus neutralization assay

The ebolavirus pseudoparticle (ebolavirus-pp) neutralization assay^12^ was performed to assess the neutralizing activity of previously reported mAbs and vaccine-induced antibody responses in mice and rabbits. Ebolavirus-pps were generated by the co-transfection of HEK293T cells with the HIV-1 pNL4-3.lucR-E-plasmid (NIH AIDS reagent program: https://www.aidsreagent.org/) and the expression plasmid encoding the GP gene of an EBOV Makona strain (GenBank accession no. KJ660346) or BDBV Uganda strain (GenBank accession no. KR063673) at a 4:1 ratio by lipofectamine 3000 (Thermo Fisher). After 48-72 h, ebolavirus-pps were collected from the supernatant by centrifugation at 3724 ×g for 10 min, aliquoted, and stored at −80 °C until use. The mAbs at a starting concentration of 10 μg/ml, or purified IgGs at a starting concentration of 300 μg/ml for mouse and 1000 μg/ml for rabbit, were mixed with the supernatant containing ebolavirus-pps and incubated for 1 h at 37°C in white solid-bottom 96-well plates (Corning). Based on recent studies on EBOV infectivity in various cell lines^72,109^, 293 T cells or TZM-bl cells were used for ebolavirus-pp neutralization assays. Briefly, HEK293T cells or TZM-bl cells at 1 × 10^4^ were added to each well and the plate was incubated at 37 °C for 48 h. After incubation, overlying media was removed, and cells were lysed. The firefly luciferase signal from infected cells was determined using the Bright-Glo Luciferase Assay System (Promega) according to the manufacturer’s instructions. Data were retrieved from a BioTek microplate reader with Gen 5 software. The average background luminescence from a series of uninfected wells was subtracted from each well, and neutralization curves were generated using GraphPad Prism 8.4.3, in which values from wells were compared against a well containing ebolavirus-pp only. The same HIV-1 vectors pseudotyped with the murine leukemia virus (MLV) Env gene, termed MLV-pps, were produced in 293 T cells and included in the neutralization assays as a negative control. Because non-NAb c13C6 exhibited enhanced MLV-pp infection, the MLV-pp assay was also used to detect and quantify the glycan cap-directed non-NAb response in immunized animal samples.

### Bulk sorting of EBOV GPΔmuc-specific mouse splenic B cells

Spleens were harvested from mice 15 days after the last immunization, and the cell suspension was prepared. Dead cells were excluded by staining with the Fixable Aqua Dead Cell Stain kit (Thermo Fisher L34957). FcγIII (CD16) and FcγII (CD32) receptors were blocked by adding 20 µl of 2.4G2 mAb (BD Pharmigen, catalog no. N553142). The cells were then incubated with 10 μg/ml of a biotinylated GPΔmuc-WL^2^P^2^-1TD0-Avi trimer. Briefly, the probe was generated by biotinylation of the GPΔmuc-WL^2^P^2^-1TD0-Avi trimer using biotin ligase BirA according to the manufacturer’s instructions (Avidity). Biotin excess was removed by SEC on a Superdex 200 10/300 column (GE Healthcare). In the SEC profile, the Avi-tagged GPΔmuc trimer peak was centered at 10.0-11.0 ml, whereas a broader peak of biotin ligase was found at 20 ml. Cells and biotinylated proteins were incubated for 5 min at 4 °C, followed by the addition of 2.5 µl of anti-mouse IgG fluorescently labeled with FITC (Jackson ImmunoResearch catalog no. 115-095-071) and incubated for 15 min at 4 °C. Finally, 5 µl of premium-grade allophycocyanin (APC)-labeled streptavidin was added to cells and incubated for 15 min at 4 °C. In each step, the cells were washed with PBS and the sorting buffer was 0.5 ml of FACS buffer. FITC^+^ APC^+^ GPΔmuc-specific B cells were sorted using MoFloAstrios into one well of a 96-well plate with 20μl of lysis buffer. Gating strategies used in antigen-specific mouse B cell sorting are exemplified by the flow-chart in **Fig. S9b**. Briefly, antigen-specific mouse splenic B cells were isolated by gating on single cells that were live/dead marker negative, mouse IgG positive, and biotinylated EBOV GP positive. Flow cytometry data were collected using the Summit v6.3 software.

### NGS and bioinformatics analysis of mouse B cells

Previously, a 5′-rapid amplification of cDNA ends (RACE)-polymerase chain reaction (PCR) protocol was reported for the unbiased sequencing of mouse B-cell repertoires^65^. In this study, this protocol was applied to analyze bulk-sorted, GP-specific mouse splenic B cells. Briefly, 5′-RACE cDNA was obtained from bulk-sorted splenic B cells of each mouse with the SMART-Seq v4 Ultra Low Input RNA Kit for Sequencing (TaKaRa). The IgG PCRs were set up with Platinum *Taq* High-Fidelity DNA Polymerase (Life Technologies) in a total volume of 50 µl, with 5 μl of cDNA as the template, 1 μl of 5′-RACE primer, and 1 μl of 10 μM reverse primer. The 5′-RACE primer contained a PGM/S5 P1 adaptor, while the reverse primer contained a PGM/S5 A adaptor. We adapted the mouse 3′-C_γ_1-3/3′-C_μ_ inner primers and 3′-mC_κ_ outer primer as reverse primers for the 5′-RACE PCR processing of heavy and light (κ) chains (**Table S1**). A total of 25 cycles of PCR was performed and the expected PCR products (500-600 bp) were gel purified (Qiagen). NGS was performed on the Ion S5 GeneStudio system. Briefly, heavy and light (κ) chain libraries from the same mouse were quantitated using a Qubit® 2.0 Fluorometer with Qubit® dsDNA HS Assay Kit and then mixed at a 3:1 ratio before being pooled with antibody libraries of other mice at an equal ratio for sequencing. Template preparation and (Ion 530) chip loading were performed on Ion Chef using the Ion 520/530 Ext Kit, followed by sequencing on the Ion S5 system with default settings. The mouse Antibodyomics pipeline^65^ was used to process raw NGS data and derive quantitative profiles for germline gene usage, degree of SHM, and H/KCDR3 loop length.

### Statistics and Reproducibility

SEC was performed for all GP/NP constructs at least once for *in vitro* characterization and multiple times during protein production for animal studies. Representative SEC profiles were selected for comparison. BN-PAGE was performed for all GP/NP constructs at least once during screening, with selected constructs run on the same gel to facilitate visual comparison. DSC was performed up to three times to validate key thermal parameters and thermographs. Negative-stain EM was performed routinely for all NP constructs during in vitro characterization and protein production for animal studies. All ELISA binding assays were performed in duplicates. Due to the limited availability of purified mouse IgG samples, ebolavirus-pp neutralization assays were performed without duplicates. For the panel of 10 representative antibodies and purified rabbit IgG samples, ebolavirus-pp neutralization assays were performed in duplicates. An unpaired two-tailed *t* test was performed in GraphPad Prism 8.4.3 to determine *P* values in the analysis of binding antibody response and mouse B-cell repertoires. The level of statistical significance is indicated as: *, *P* < 0.05; **, *P* < 0.01; ***, *P* < 0.001; ****, *p* < 0.0001.

## Supporting information

Supplementary figures and ables

## Data Availability

The X-ray crystallographic coordinates for two rationally redesigned GPΔmuc structures in this study have been deposited in the Protein Data Bank (PDB, https://www.rcsb.org/), under accession codes 7JPI (http://doi.org/10.2210/pdb7jpi/pdb) and 7JPH (http://doi.org/10.2210/pdb7jph/pdb). The mouse B-cell NGS datasets have been deposited in the NIH Sequence Read Archive (SRA, https://www.ncbi.nlm.nih.gov/sra), with the identifier PRJNA718964 (https://www.ncbi.nlm.nih.gov/bioproject/PRJNA718964/). The authors declare that the data supporting the findings of this study are available within the article and its Supplementary Information files. Source data are provided with this paper.

## Acknowledgments

We thank Mansun Law and Juan Carlos de la Torre at The Scripps Research Institute and Sujan Shresta at the La Jolla Institute for Immunology for helpful discussions. We thank Christiana Corbaci for creating images used in Figs. 5a, 5e, and 6a and Michael Arends for proofreading the manuscript. Diffraction data were collected at the Advanced Photon Source (APS) beamline 23-IDD, and Stanford Synchrotron Radiation Lightsource (SSRL) beamline 12-2. Use of the APS was supported by the DOE, Basic Energy Sciences, Office of Science, under contract no. DE-AC02-06CH11357. Use of the SSRL was supported by the US Department of Energy, Basic Energy Sciences, Office of Science, under contract no. DE-AC02-76SF00515. This work was funded in part by NIH grants AI129698 (to J.Z.), AI140844 (to J.Z. and I.A.W.), UfoVax/SFP-2018-0416, and UfoVax/SFP-2018-1013 (to J.Z.).

## Author contributions

Project design by L.H., A.C., X.L., I.A.W., and J.Z.; rational design of EBOV GPΔMuc trimers and nanoparticles by L.H. and J.Z.; plasmid design and processing by L.H. and C.S.; antigen production, purification, and biochemical characterization by A.C., L.H., X.L., E.K., and T.N.; EBOV antibody production by X.L., E.K., and T.N.; negative-stain EM by T.A., G.O., and A.B.W.; DSC measurement by S.K.; crystallography by A.C. and R.L.S.; BLI by L.H. and J.Z.; mouse plasma-antigen ELISA by L.H. and X.L.; plasma IgG purification by L.H. and C.S. ebolavirus-pp and MLV-pp neutralization assays by X.L.; antigen-specific mouse B cell sorting by C.S. and L.H.; mouse B cell library preparation and NGS by L.H. and C.S.; bioinformatics analysis by L.H. and J.Z.; Manuscript written by L.H., A.C., X.L., I.A.W., and J.Z. All authors were asked to comment on the manuscript. The TSRI manuscript number is 30009.

## Competing interests

The authors declare that they have no competing interests.

## Notes

### Competing Interest Statement

The authors have declared no competing interest.

