## Supplementary figures and ables for "Single-component multilayered self-assembling nanoparticles presenting rationally designed glycoprotein trimers as Ebola virus vaccines"

**a**

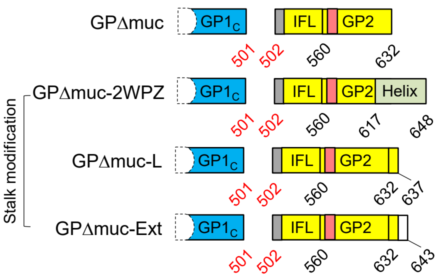


**b**

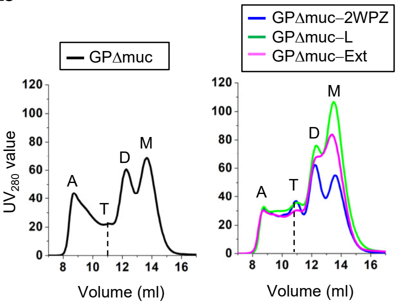


**c**

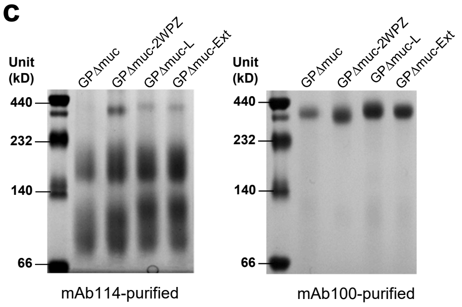


**d** Antigenic profiling of four mAb100/SEC-purified, stalk-modified EBOV GP/GPΔmuc-foldon trimers

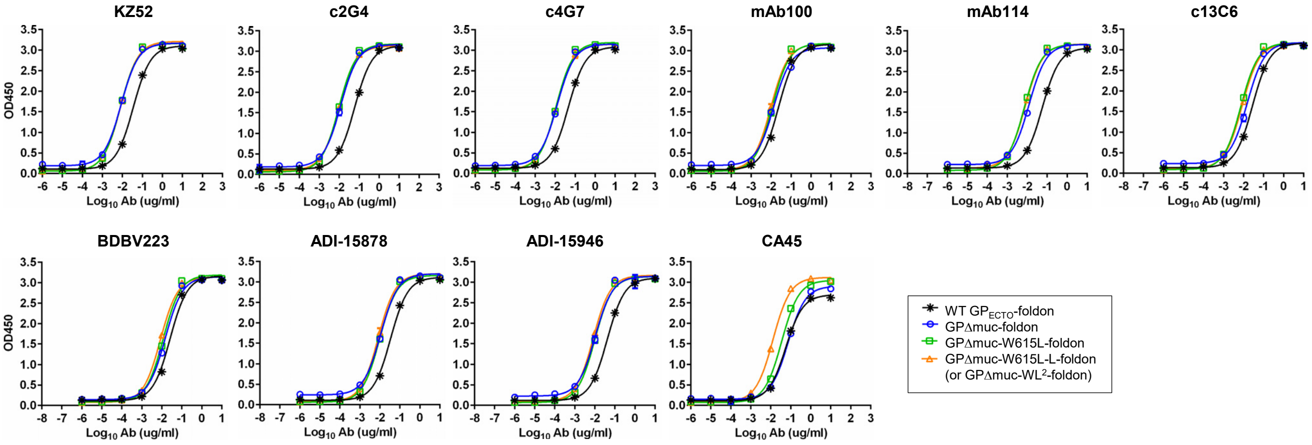


**e**

**$EC_{50}$ values (μg/ml) of four EBOV GP/GPΔmuc trimers binding to 10 antibodies [a].**

| | KZ52 | c2G4 | c4G7 | mAb100 | mAb114 | c13C6 | BDBV223 | ADI-15878 | ADI-15946 | CA45 |
|---|---|---|---|---|---|---|---|---|---|---|
| WT $GP_{ECTO}$-foldon | 0.035 | 0.059 | 0.045 | 0.024 | 0.056 | 0.028 | 0.026 | 0.034 | 0.042 | 0.056 |
| GPΔmuc-foldon | 0.009 | 0.012 | 0.013 | 0.013 | 0.012 | 0.015 | 0.015 | 0.011 | 0.010 | 0.073 |
| GPΔmuc-W615L-foldon | 0.008 | 0.009 | 0.011 | 0.011 | 0.007 | 0.007 | 0.012 | 0.010 | 0.010 | 0.035 |
| GPΔmuc-$WL^2$-foldon | 0.008 | 0.010 | 0.011 | 0.010 | 0.008 | 0.008 | 0.008 | 0.008 | 0.008 | 0.013 |

[a] $EC_{50}$ values were calculated from the besting fitting in GraphPad Prism v8.4.3.

**f** Antigenic profiling of an mAb100/SEC-purified, stalk/$HR1_C$-modified EBOV GPΔmuc-foldon trimer

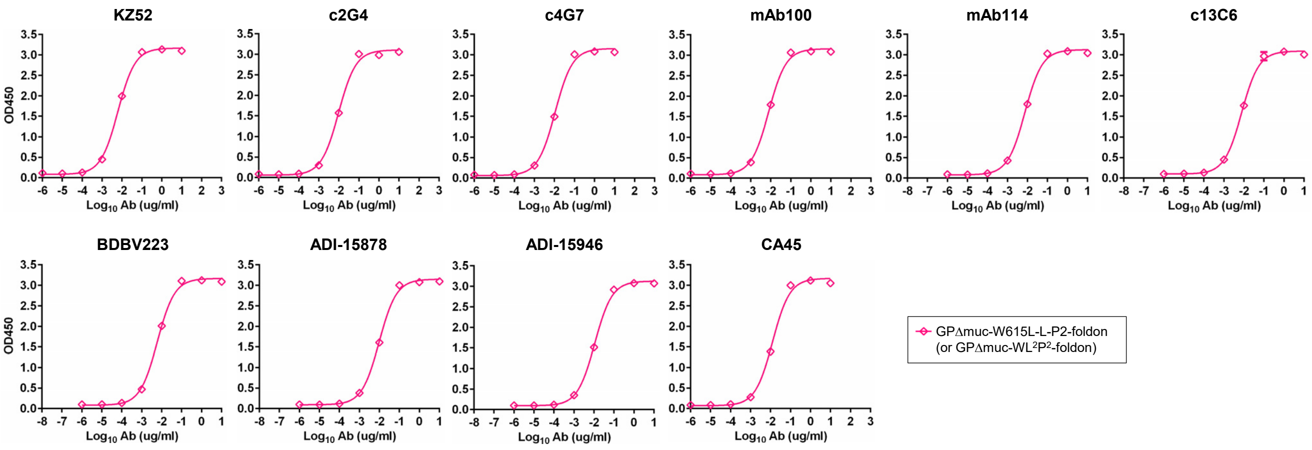


**g**

**$EC_{50}$ values (μg/ml) of a stalk/$HR1_C$-modified EBOV GPΔmuc trimer binding to 10 antibodies [a].**

| | KZ52 | c2G4 | c4G7 | mAb100 | mAb114 | c13C6 | BDBV223 | ADI-15878 | ADI-15946 | CA45 |
|---|---|---|---|---|---|---|---|---|---|---|
| GPΔmuc-$WL^2P^2$-foldon | 0.006 | 0.010 | 0.011 | 0.008 | 0.007 | 0.008 | 0.006 | 0.010 | 0.011 | 0.012 |

[a] $EC_{50}$ values were calculated from the besting fitting in GraphPad Prism v8.4.3.

h

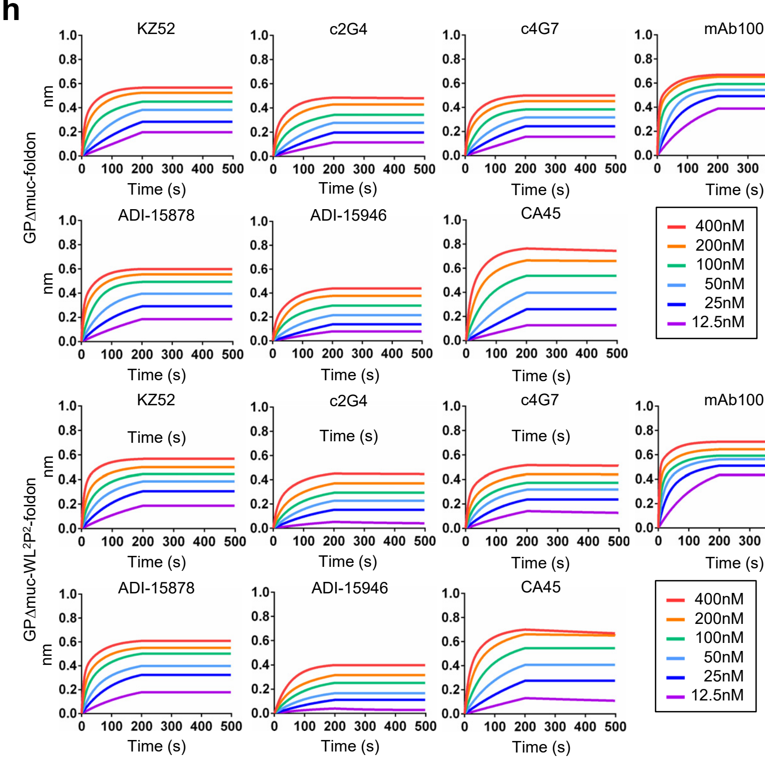


**Supplementary Fig. 1. Design, screening, and antigenic characterization of EBOV GPΔmuc trimers with modified stalk and HR1$_C$.** (**a**) Schematic representation of mucin-deleted GP (GPΔmuc) and three stalk designs (GPΔmuc-2WPZ, GPΔmuc-L, and GPΔmuc-Ext). In the GPΔmuc-2WPZ construct design, in addition to replacing aa 617-632 with the coiled-coil region (aa 3-33) of 2WPZ, a double mutation (P612G/W615F) was introduced to reduce structural strain within the mutated stalk. (**b**) SEC profiles of four GPΔmuc constructs obtained from a Superdex 200 10/300 column following transient expression in 250 ml HEK293 F cells and mAb114 purification. Left: the SEC curve of GPΔmuc shown in black line; Right: three stalk designs shown in blue, green, and magenta lines for GPΔmuc-2WPZ, -L, and -Ext, respectively. (**c**) BN-PAGE of WT GPΔmuc and three stalk designs (GPΔmuc-2WPZ, -L, and -Ext) purified by mAb114 (left) and mAb110 (right) columns. (**d**) ELISA curves of four mAb100/SEC-purified, stalk-modified EBOV GP/GPΔmuc-foldon trimers binding to 10 antibodies. (**e**) Summary of $EC_{50}$ values (μg/ml) of EBOV GP/GPΔmuc-foldon trimers binding to 10 antibodies. **(f)** ELISA curves of an mAb100/SEC-purified, stalk/HR1$_C$-modified EBOV GPΔmuc-foldon trimer binding to 10 antibodies. (**g**) Summary of $EC_{50}$ values (μg/ml) of a stalk/HR1$_C$-modified EBOV GPΔmuc-foldon trimer binding to 10 antibodies. In (e) and (g), $EC_{50}$ values were calculated for all ELISA data in Prism 8.4.3. (**h**) Antigenic evaluation of GPΔMuc-foldon and GPΔmuc-WL$^2$P$^2$-foldon trimers using BLI and 10 representative antibodies. Sensorgrams were obtained from an Octet RED96 instrument using six concentrations (400-12.5 nM by twofold dilution) and AHC biosensors (see Methods). Source data are provided as a Source Data file.

### Supplementary Fig. 2

**a**

**Inter-protomer disulfide bond (SS) mutations** [a]

| # | Res1 | Res2 | $d_{C\beta}$ | Comments |
|---|---|---|---|---|
| **SS1** | TYR534 | PHE153 | 3.82 | IFL-head |
| **SS2** | ALA538 | GLY91 | 3.99 | IFL-head |
| **SS3** | THR520 | ALA575 | 4.01 | IFL-NHR |
| — | TRP615 | ILE603 | 4.14 | Hydrophobic cluster |
| **SS4** | ILE532 | GLY157 | 4.20 | IFL-head |
| **SS5** | ASP522 | ALA575 | 4.41 | IFL-NHR |
| | GLU611 | HIS602 | 4.59 | SS cysteine |
| **SS6** | LYS56 | GLY599 | 4.60 | Stalk (HR2) |
| — | CYS609 | THR600 | 4.62 | SS cysteine |
| | TRP531 | GLN567 | 4.73 | $d_{C\beta}$ > **cutoff** |

[a] $C_\beta$-$C_\beta$ distance of 4.72 Å as the indicator of S-S bond formation

**b**

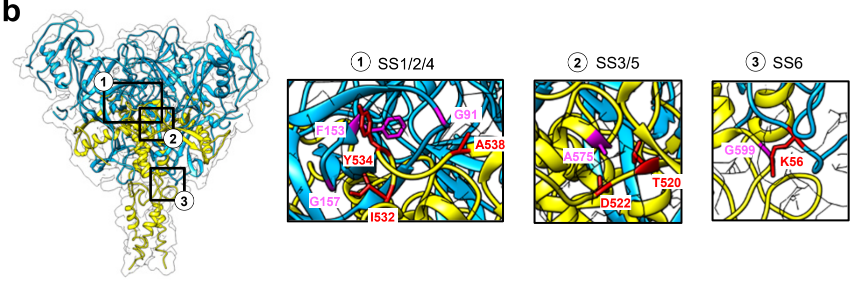


**c**

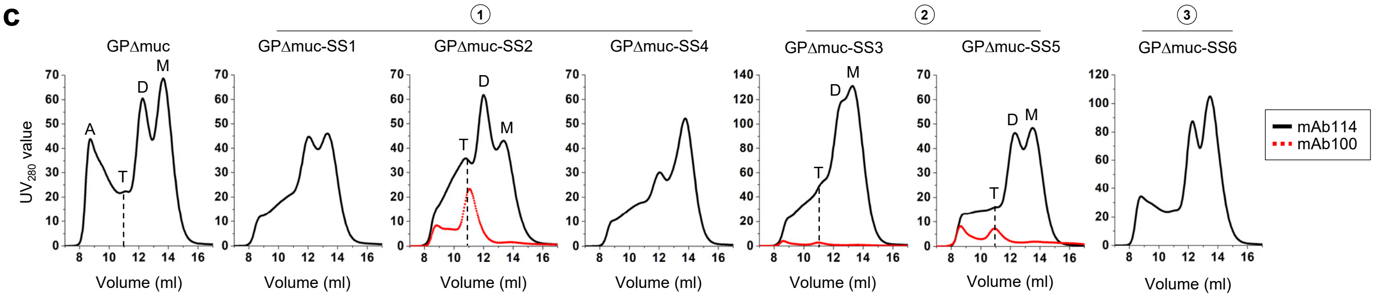


**d**

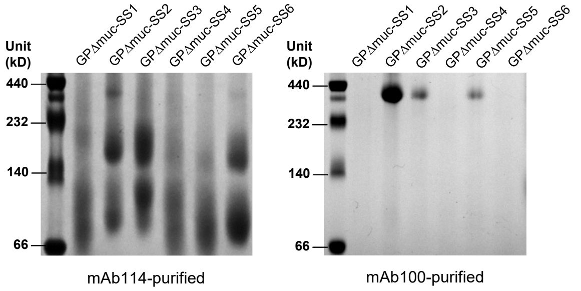


**e**

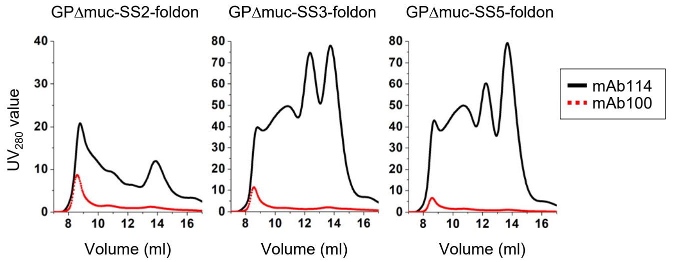


**f** **Antigenic profiling of mAb100/SEC-purified GPΔmuc-SS2 trimer (without foldon)**

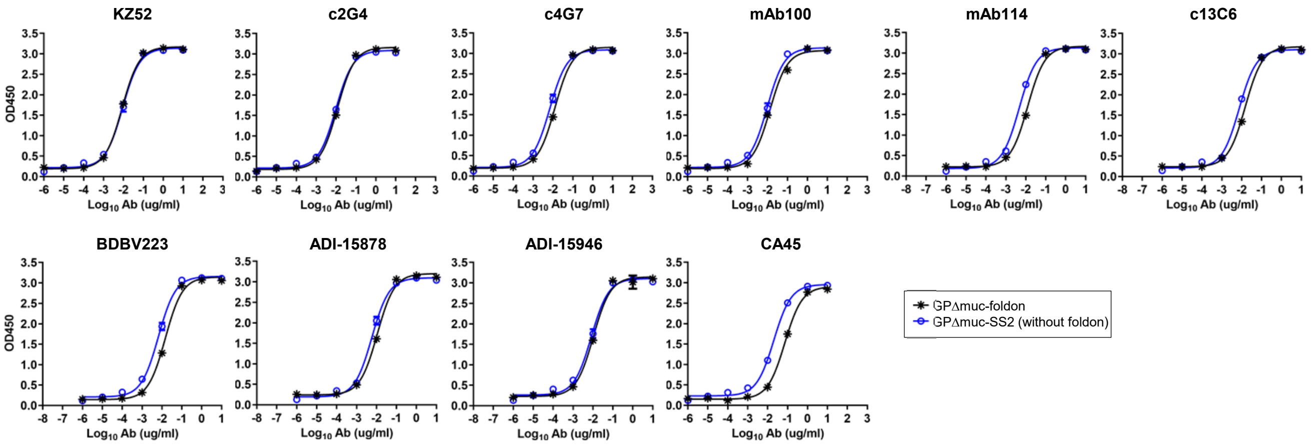


**g**

**$EC_{50}$ values (μg/ml) of EBOV GPΔmuc-SS2 trimer binding to 10 antibodies** [a]**.**

| | KZ52 | c2G4 | c4G7 | mAb100 | mAb114 | c13C6 | BDBV223 | ADI-15878 | ADI-15946 | CA45 |
|---|---|---|---|---|---|---|---|---|---|---|
| GPΔmuc-foldon | 0.009 | 0.012 | 0.013 | 0.013 | 0.012 | 0.015 | 0.015 | 0.011 | 0.010 | 0.073 |
| GPΔmuc-SS2 | 0.009 | 0.010 | 0.007 | 0.010 | 0.005 | 0.007 | 0.007 | 0.006 | 0.008 | 0.020 |

[a] $EC_{50}$ values were calculated from the besting fitting in GraphPad Prism v8.4.3. ELISA data for the GPΔmuc-foldon trimer is included for comparison.

**Supplementary Fig. 2. Design and characterization of EBOV GPΔmuc trimers with inter-protomer disulfide bonds.** (**a**) Inter-protomer disulfide bond (SS) mutations identified using the atomic structure of a WT GPΔmuc trimer (PDB: 5JQ3) with residue numbers, $C_\beta$-$C_\beta$ distance, and location listed as the key parameters for selection. (**b**) Structure and molecular surface of EBOV GPmMuc trimer (PDB: 5JQ3) in transparent molecular surface with the three SS classes labeled according to their locations. Zoomed-in views of the three SS classes, SS1/2/4, SS3/5, and SS6, are shown on the right. (**c**) SEC profiles of mAb114-purified GPΔmuc and six GPΔmuc-SS variants from a Superdex 200 10/300 column. SEC profiles of mAb100-purified SS2, SS3, and SS5 are shown in the red dotted line. Trimer (T), dimer (D), and monomer (M) peaks are labeled on the SEC profiles for SS2, SS3 and SS5, with the trimer peak marked with a black dashed line. (**d**) BN-PAGE of six GPΔmuc-SS variants after purification using an mAb114 column (left) and an mAb100 column (right). (**e**) SEC profiles of GPΔmuc-SS2/3/5-foldon constructs after purification using an mAb114 column (in the black line) and an mAb100 column (in the red dotted line) purification. No trimer peak was observed for any of the three GPΔmuc-SS-foldon constructs. (**f**) ELISA curves of the mAb100/SEC-purified GPΔmuc-SS2 trimer binding to 10 representative antibodies. (**g**) Summary of $EC_{50}$ values (μg/ml) of the mAb100/SEC-purified GPΔmuc-SS2 trimer binding to 10 representative antibodies. In (f) and (g), ELISA data for the GPΔmuc-foldon trimer are included for comparison. $EC_{50}$ values were calculated for all ELISA data in Prism 8.4.3. Source data are provided as a Source Data file.

**Supplementary Fig. 3**

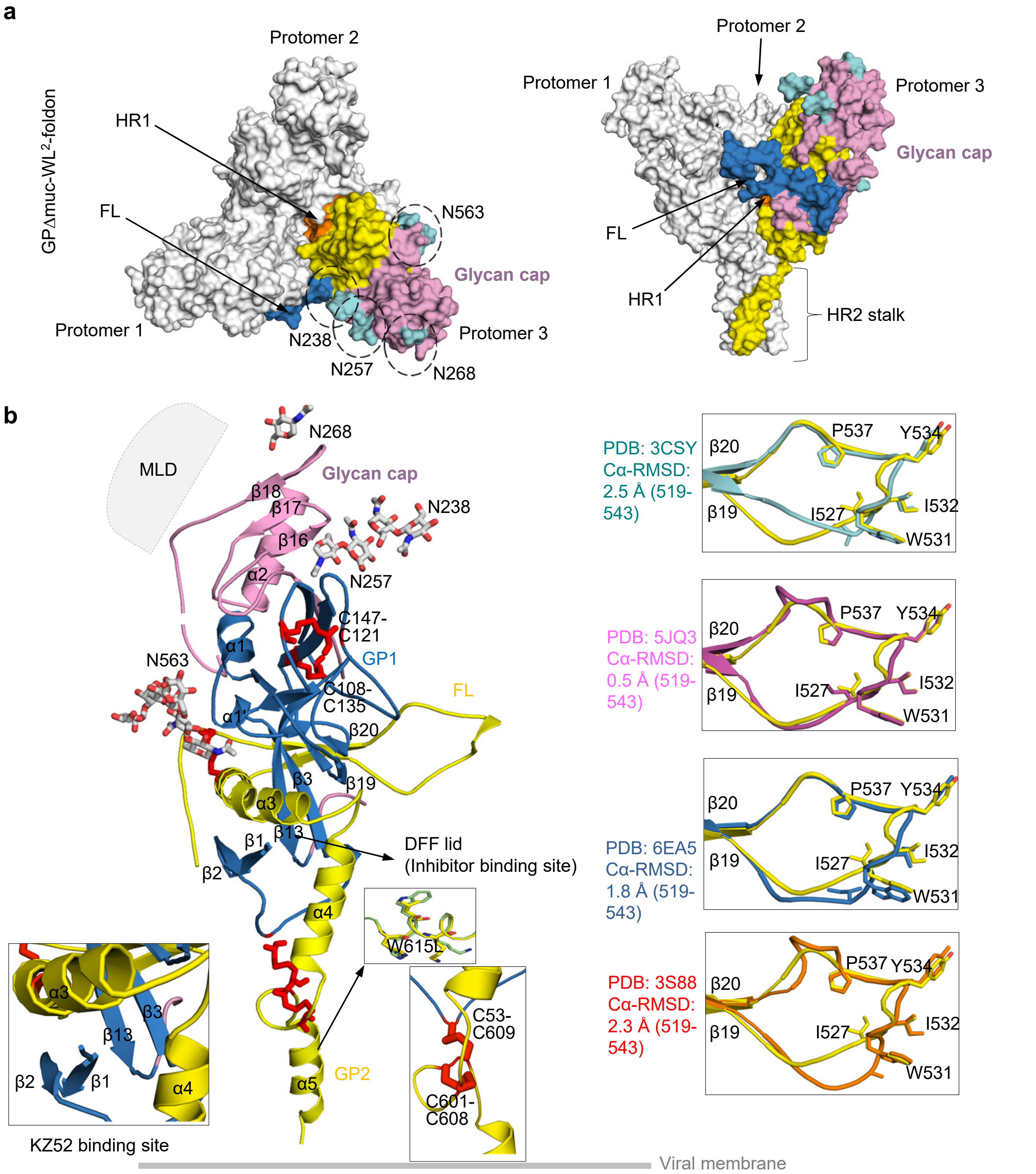


**Supplementary Fig. 3. Crystal structure of GPΔmuc-WL²-foldon at 2.3 Å resolution.** (**a**) Molecular surface representation of GPΔmuc-WL²-foldon trimer structure (Left: top view; Right: side view). Protomers 1 and 2 are shown in light grey and protomer 3 is colored as follows: HR1 (orange), IFL(blue), glycan cap (pink), and HR2 (yellow). *N*-linked glycans at N238, N257, N268 and N563 are highlighted in circles and colored in cyan. (**b**) Ribbon representation of a GPΔmuc-WL²-foldon protomer with the same color scheme as in (a). Glycans and disulfide bonds are shown as color-coded (by atom type) and red sticks, respectively. The mucin-like domain (MLD) is shown as a grey oval. Binding sites for KZ52 and small-molecule inhibitors are indicated. The W615L mutation and two disulfide bonds (C53-C609 and C601-C608) are shown as insets. Superposed IFLs (R519-E543) from WL² and known structures (PDB IDs: 3CSY, 5JQ3, 6EA5, and 3S88) are shown as insets (right), with Cα RMSDs of 0.5-2.5Å.

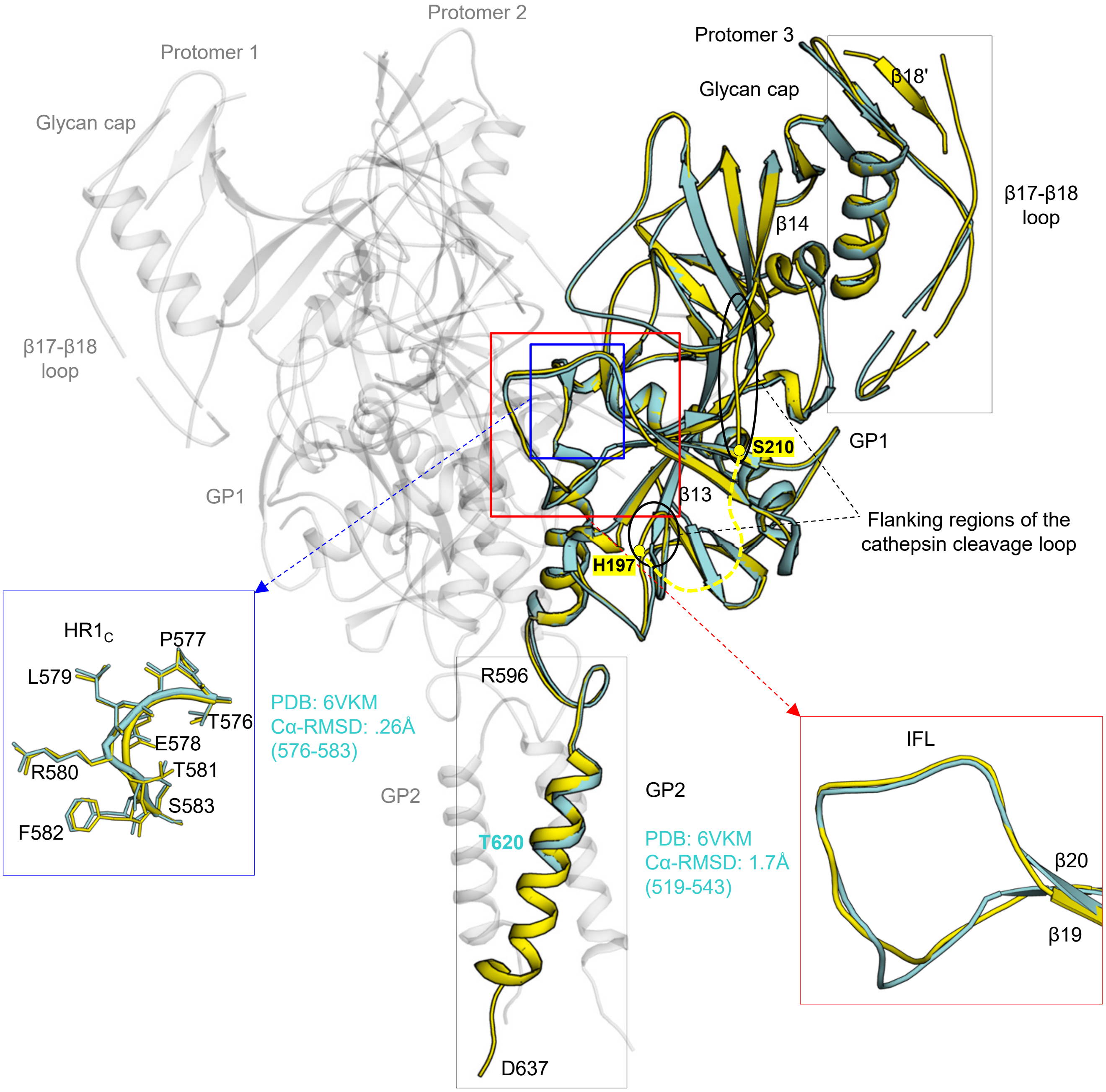


**Supplementary Fig. 4. Comparison of the Mayinga GPΔmuc-WL²-foldon structure and the previously reported Makona GPΔmuc structure containing a T577P/R588E double mutation**. Ribbon representation of the GPΔmuc-WL²-foldon trimer structure is shown, with protomers 1/2 and 3 colored in light grey and yellow, respectively. The structure of a Makona GPΔmuc protomer (PDB: 6VKM) is superimposed onto protomer 3 and shown in cyan. Superimposed $HR1_C$ regions and internal fusion loops (IFL) from the two structures are shown in the bottom left and right insets, respectively. The flanking regions of the cathepsin cleavage loop (between β13 and β14, aa 190-210) are indicated in black ovals. H197 and S210 are labeled. The unstructured cathepsin cleavage loop between H197 and S210 is indicated as a yellow dashed line.

#### Supplementary Fig. 5

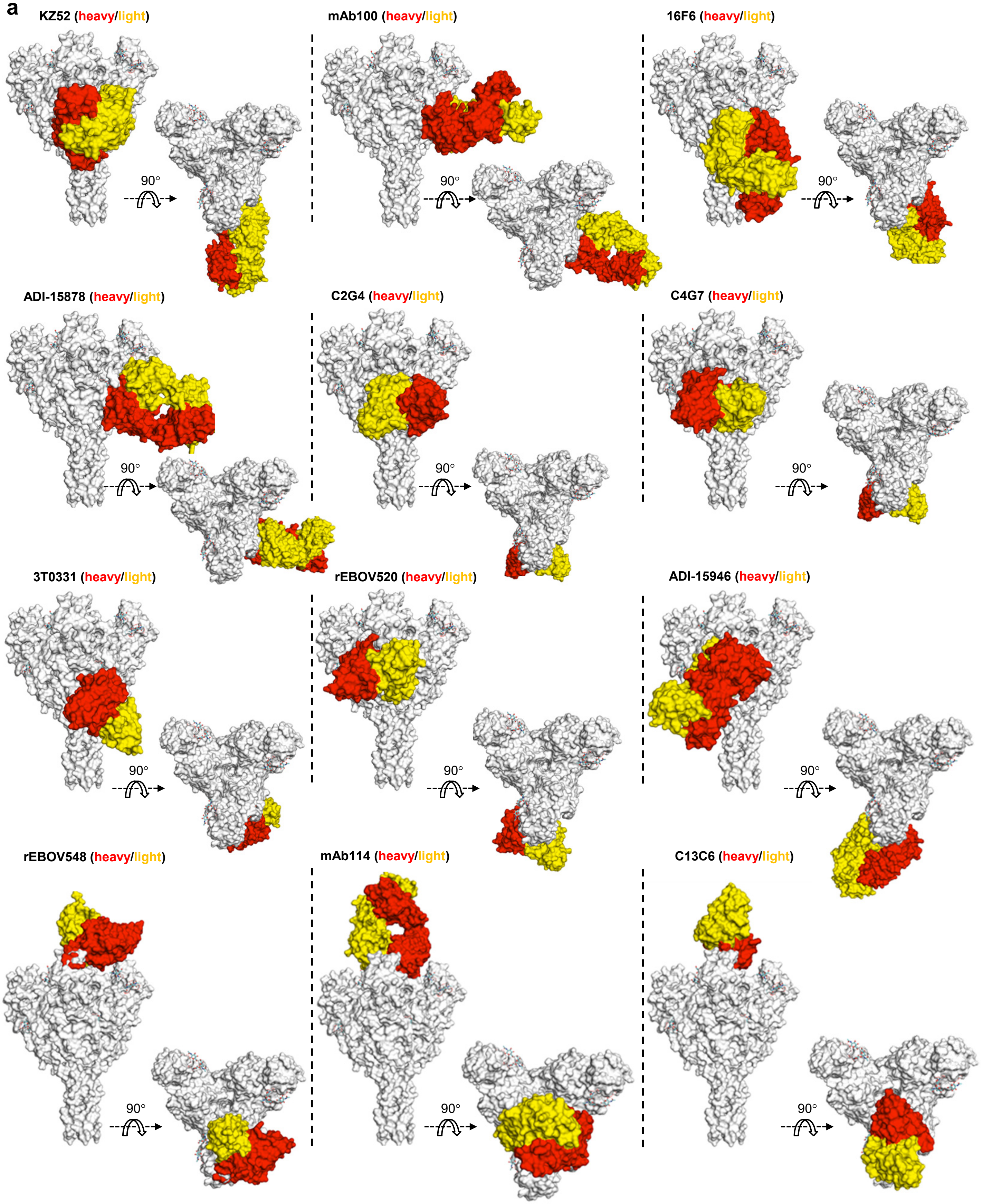

**Supplementary Fig. 5**

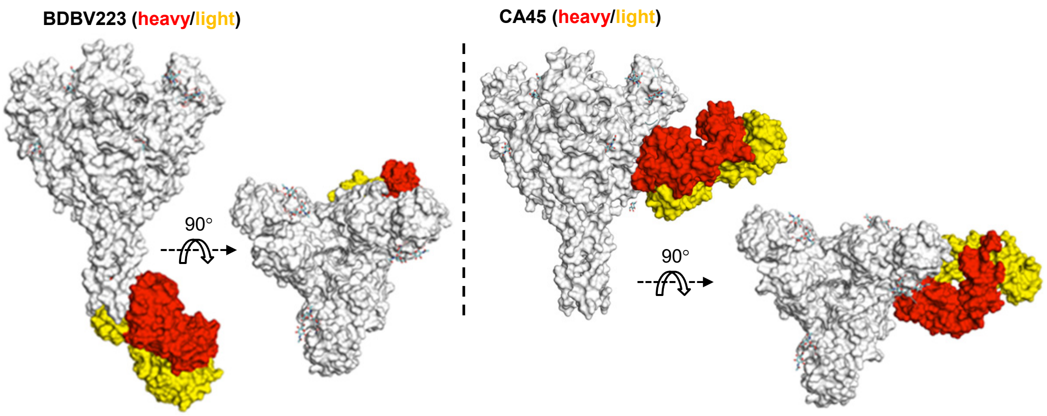


**b**

```
      32          47           60
5KEN -IPLGVIHNSTLQVSDVDKLVCRDKLSST C4G7
5KEL -IPLGVIHNSTLQVSDVDKLVCRDKLSST C13C6/C2G4
3CSY SIPLGVIHNSVLQVSDVDKLVCRDKLSST KZ52
3S88 SMPLGVVTNSTLEVTEIDQLVCKDHLAST 16F6
6EA5 -IPLGVVHNNTLQVSDIDKLVCRDKLSST ADI-15878
5FHC SIPLGVIHNSTLQVSDVDKLVCRDKLSST mAB114/mAB100
6PCI SIPLGVIHNSTLQVSDVDKLVCRDKLSST rEBOV548/rEBOV520
6MAM SIPLGVIHNSTLQVSDVDKLVCRDKLSST ADI-15946
6QD7 SIPLGVIHNSALQVSDVDKLVCRDKLSST 3T0331
6N7J ----------------------------- BDBV223
6EAY SIPLGVIHNSTLQVSDVDKLVCRDKLSST CA45
            70                             100                 120
5KEN NQLRSVGLNLEGNGVATDVPSATKRWGFRSGVPPKVVNYEAGEWAENCYNLEIKKPDGSE C4G7
5KEL NQLRSVGLNLEGNGVATDVPSATKRWGFRSGVPPKVVNYEAGEWAENCYNLEIKKPDGSE C13C6/C2G4
3CSY NQLRSVGLNLEGNGVATDVPSATKRWGFRSGVPPKVVNYEAGEWAENCYNLEIKKPDGSE KZ52
3S88 DQLKSVGLNLEGSGVSTDIPSATKRWGFRSGVPPKVVSYEAGEWAENCYNLEIKKPDGSE 16F6
6EA5 SQLASVGLNLEGNGVATDVPTATKRWGFRAGVPPKVVNYEAGEWAENCYNLDIKKADGSE ADI-15878
5FHC NQLRSVGLNLEGNGVATDVPSATKRWGFRSGVPPKVVNYEAGEWAENCYNLEIKKPDGSE mAB114/mAB100
6PCI NQLRSVGLNLEGNGVATDVPSVTKRWGFRSGVPPKVVNYEAGEWAENCYNLEIKKPDGSE rEBOV548/rEBOV520
6MAM NQLRSVGLNLEGNGVATDVPSATKRWGFRSGVPPKVVNYEAGEWAENCYNLEIKKPDGSE ADI-15946
6QD7 NQLRSVGLNLEGNGVATDVPSATKRWGFRSGVPPKVVNYEAGEWAENCYNLEIKKPDGSE 3T0331
6N7J ------------------------------------------------------------ BDBV223
6EAY NQLRSVGLNLEGNGVATDVPSATKRWGFRSGVPPKVVNYEAGEWAENCYNLEIKKPDGSE CA45
    136          186                      211                        240
5KEN RYVHKVSGTGPCALPQAKKDFFSSHPLREPVNATEDPSSGYYSTTIRYQATGFGTNETEYLFEVDNLT C4G7
5KEL RYVHKVSGTGPCALPQAKKDFFSSHPLREPVNATEDPSSGYYSTTIRYQATGFGTNETEYLFEVDNLT C13C6/C2G4
3CSY RYVHKVSGTGPCALPQAKKDFFSSHPLREPVNATEDPSSGYYSTTIRYQATGFGTNEVEYLFEVDNLT KZ52
3S88 RYVHKAQGTGPCPLAKPKETFLQSPPIREAVNYTENTSSYYATSYLEYEIENFGAQHSTTLFKIDNNT 16F6
6EA5 RYVHKVSGTGPCPLPETKKDFFEI-------------------------------------------- ADI-15878
5FHC RYVHKVSGTGPCALPQAKKDFFSSHPLREPVNATEDPSSGYYSTTIRYQATGFGTNETEYLFEVDNLT mAB114/mAB100
6PCI RYVHKVSGTGPCALPQAKKDFFSSHPLREPVNATEDPSSGYYSTTIRYQATGFGTNETEYLFEVDNLT rEBOV548/rEBOV520
6MAM RYVHKVSGTGPCALPQAKKDFFSSHPLREPVNATEDPSSGYYSTTIRYQATGFGN------------- ADI-15946
6QD7 RYVHKVSGTGPCALPQAKKDFFSSHPLREPVNATEDPSSGYYSTTIRYQATGFGTNETEYLFEVDNLT 3T0331
6N7J -------------------------------------------------------------------- BDBV223
6EAY RYVHKVSGTGPCALPQAKKDFFSSHPLREPVNATEDPSSGYYSTTIRYQATGFGTNETEYLFEVDNLT CA45
    241                         270                            302
5KEN YVQLESRFTPQFLLQLNETIYTSGKRSNTTGKLIWKVNPEIDTTIGEWAFWETKKNLIRSEELSFA-- C4G7
5KEL YVQLESRFTPQFLLQLNETIYTSGKRSNTTGKLIWKVNPEIDTTIGEWAFWETKKNLIRSEELSFTVV C13C6/C2G4
3CSY YVQLESRFTPQFLLQLNETIYTSGKRSNTTGKLIWKVNPEIDTTIGEWAFWETKKNLIRSEELSFTVV KZ52
3S88 FVRLDRPHTPQFLFQLNDTIHLHQQLSNTTGRLIWTLDANINADIGEWAFWENKKNLLRGEELSFEAL 16F6
6EA5 -------------------------------------------------------------------- ADI-15878
5FHC YVQLESRFTPQFLLQLNETIYTSGKRSNTTGKLIWKVNPEIDTTIGEWAFWETKKNLIRSEELSFSR- mAB114/mAB100
6PCI YVQLESRFTPQFLLQLNETIYASGKRSNTTGKLIWKVNPEIDTTIGEWAFWETKKNLIRSEELSFTN- rEBOV548/rEBOV520
6MAM -------------------------------------------------------------------- ADI-15946
6QD7 YVQLESRFTPQFLLQLNETIYTSGKRSNTTGKLIWKVNPEIDTTIGEWAFWETKKNLIRSEELSFTVV 3T0331
6N7J -------------------------------------------------------------------- BDBV223
6EAY YVQLESRFTPQFLLQLNETIYTSGKRSNTTGKLIWKVNPEIDTTIGEWAFWETKKNLIRSEELSFTVV CA45
    502                         545                        619
5KEN --IVNAQPKCNPNLHYWTTQDEGAAITEGLMHNQDGLICGLRQLANETTQAL---------------- C4G7
5KEL EAIVNAQPKCNPNLHYWTTQDEGAAITEGLMHNQDGLICGLRQLANETTQALTDKIDQIIHDFVDKTL C13C6/C2G4
3CSY EAIVNAQPKCNPNLHYWTTQDEGAAITEGLMHNQDGLICGLRQLANETTQALTDKIDQIIHDFVD--- KZ52
3S88 QTNTKATGKCNPNLHYWTAQEQHNAATEGLMHNQNALVCGLRQLANETTQAL--TSPIVKSFNRN--- 16F6
6EA5 --TLRTQAKCNPNLHYWTTQDEGAAITEGIMHNQNGLICGLRQLANETTQAL---------------- ADI-15878
5FHC EAIVNAQPKCNPNLHYWTTQDEGAAIIEGLMHNQDGLICGLRQLANETTQAL---------------- mAB114/Mab100
6PCI EVIVNAQPKCNPNLHFWTTQDEGAAITEGLMHNQDGLICGLRQLANETTQALTDKIDQIIHDDDDKAG rEBOV548/rEBOV520
6MAM EAIVNAQPKCNPNLHYWTTQDEGAAIIEGLMHNQDGLICGLRQLANETTQAL---------------- ADI-15946
6QD7 EAIVNAQPKCNPNLHYWTTQDEGAAIIEGLMHNQDGLICGLRQLANETTQALTDKIDQIIHDFVDGSG 3T0331
6N7J ----------------------------------------------------TDKIDQIIHDFIDKPL BDBV223
6EAY EAIVNAQPKCNPNLHYWTTQDEGAAITEGLMHNQDGLICGLRQLANETTQALTDKIDQIIHDFVDKTL CA45
```

GP1

GP2

**Supplementary Fig. 5. Structural analysis of known GP/antibody complexes.** (**a**) Molecular surface representation of the GPΔmuc-WL$^2$P$^2$ trimer bound to antibodies by docking the crystal structure of GPΔmuc-WL$^2$P$^2$-foldon (grey) into previously determined GP/antibody complex structures (Left: side view; Right: top view). Antibody heavy and light chains are shown in red and yellow, respectively. (**b**) Sequence analysis of antibody-contacting residues (≤ 4.0 Å) in GP1 and GP2. In the ClustalW alignment, antibody-contacting residues are colored in green except for 5KEL, 5FHC, and 6PCI, for which residues interacting with the first and second antibodies are colored in red and cyan, respectively, to define their respective epitopes.

**Supplementary Fig. 6**

**a**

**Dimeric locking domains (LD) for stabilizing the NP-forming interface**

>**LD1** 1NI8_A (**40aa** after C-terminal deletion)
SEALKILNNIRTLRAQARECTLETLEEMLEKLEVVVNERR~~**EEESAA**~~

>**LD2** 4AYA_B (**44aa** after N-/C-terminal deletion)
MNDCYSKLKELVPSIPQNKKVSKMEILQHVIDYILDLQIALDSH

>**LD3** 1OVX_A (**50aa** after N-terminal deletion)
~~**GHHHHHHTDKRKDGSGK**~~LLYCSFCGKSQHEVRKLIAGPSVYICDECVDLCNDIIREEIKEVAPHRER

>**LD4** 2MG4_A (**58aa** after N-terminal deletion)
~~**MEKRPRTE**~~FSEEQKKALDLAFYFDRRLTPEWRRYLSQRLGLNEEQIERWFRRKEQQIGWSHPQFEK

>**LD5** 2JV7_A (**78aa, without deletion**)
DQPSVGDAFDKYNEAVRVFTQLSSAANCDWAACLSSLSASSAACIAAVGELGLDVPLDLACAATATSSATEACKGCLW

>**LD6** 1JR5_A (88aa after N-terminal deletion)
~~**MN**~~KNIDTVREIITVASILIKFSREDIVENRANFIAFLNEIGVTHEGRKLNQNSFRKIVSELTQEDKKTLIDEFNEGFEGVYRYLEMYTNK

>**LD7** 1PZQ_A (46aa after N-/C-terminal deletion)
~~**GSAA**~~SPAVDIGDRLDELEKALEALSAEDGHDDVGQRLESLLRRWNSRRAD~~**APSTSAISED**~~

**>LD8** 1R2A_A (**39aa** after N-terminal deletion)
~~**HMGHIQI**~~PPGLTELLQGYTVEVLRQQPPDLVDFAVEYFTRLREARR

**>LD9** 2JRX_A (**65aa** after N-/C-terminal deletion and glycan deletion)
~~**MPQISR**~~YSDEQVEQLLAELLNVLEKHKAPTDLSLMVLGNMVTNLINT**A**IAPAQRQAIANSFARALQSSINE~~**DKAHLEHHHHHH**~~

**b**

**Antigenic evaluation of NPs presenting a stabilized GPΔmuc trimer**

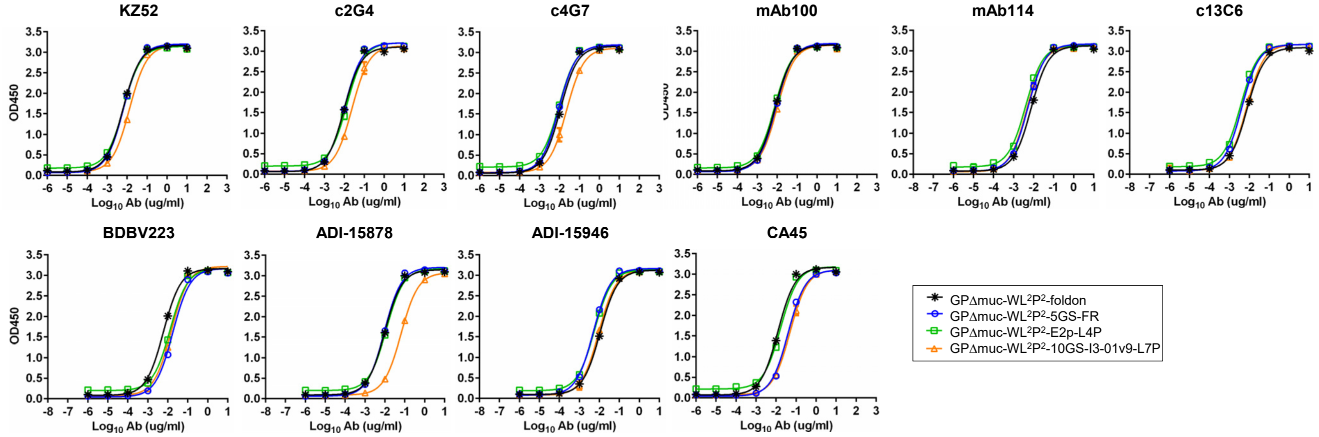


**c**

**$EC_{50}$ values (μg/ml) of EBOV GPΔmuc trimer and nanoparticles binding to 10 antibodies [a].**

| | KZ52 | c2G4 | c4G7 | mAb100 | mAb114 | c13C6 | BDBV223 | ADI-15878 | ADI-15946 | CA45 |
|---|---|---|---|---|---|---|---|---|---|---|
| GPΔmuc-WL$^2$P$^2$-foldon | 0.006 | 0.010 | 0.011 | 0.008 | 0.007 | 0.008 | 0.006 | 0.010 | 0.011 | 0.012 |
| GPΔmuc-WL$^2$P$^2$-5GS-FR | 0.007 | 0.010 | 0.009 | 0.009 | 0.005 | 0.004 | 0.018 | 0.009 | 0.005 | 0.040 |
| GPΔmuc-WL$^2$P$^2$-E2p-L4P | 0.007 | 0.013 | 0.009 | 0.008 | 0.004 | 0.004 | 0.013 | 0.012 | 0.006 | 0.017 |
| GPΔmuc-WL$^2$P$^2$-10GS-I3-01v9-L7P | 0.013 | 0.024 | 0.023 | 0.010 | 0.005 | 0.007 | 0.015 | 0.065 | 0.009 | 0.049 |

[a] $EC_{50}$ values were calculated from the besting fitting in GraphPad Prism 8.4.3.

#### Supplementary Fig. 6

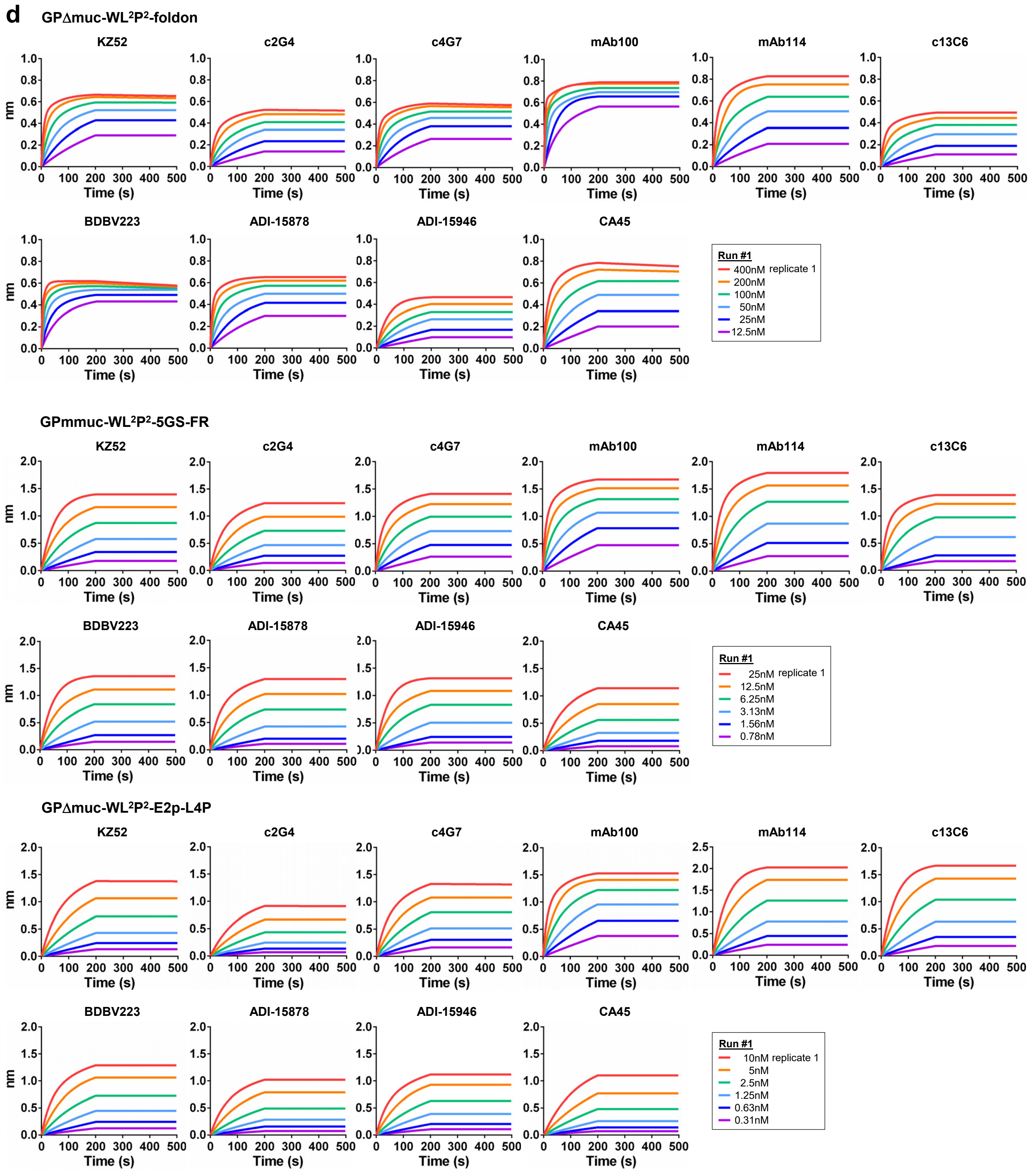

**Supplementary Fig. 6**

GPΔmuc-WL²P²-I3-01v9-L7P

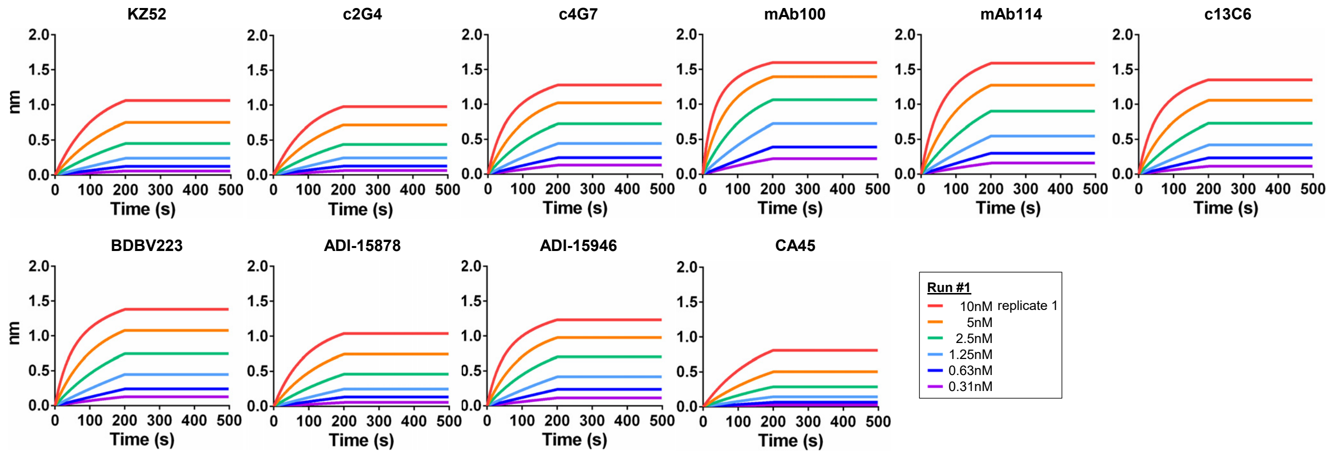


**e** GPΔmuc-WL²P²-foldon

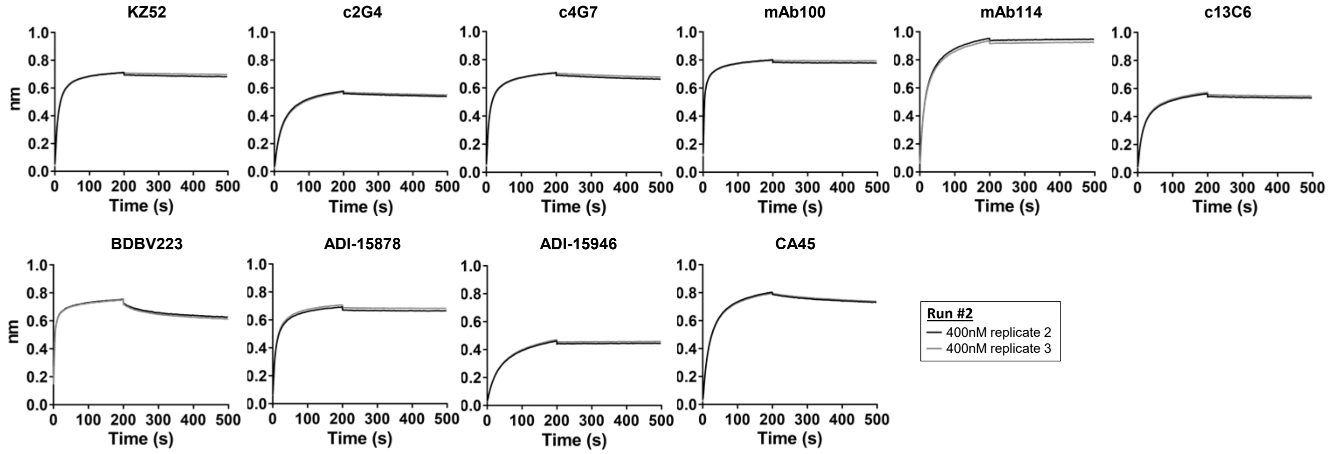


GPmmuc-WL²P²-5GS-FR

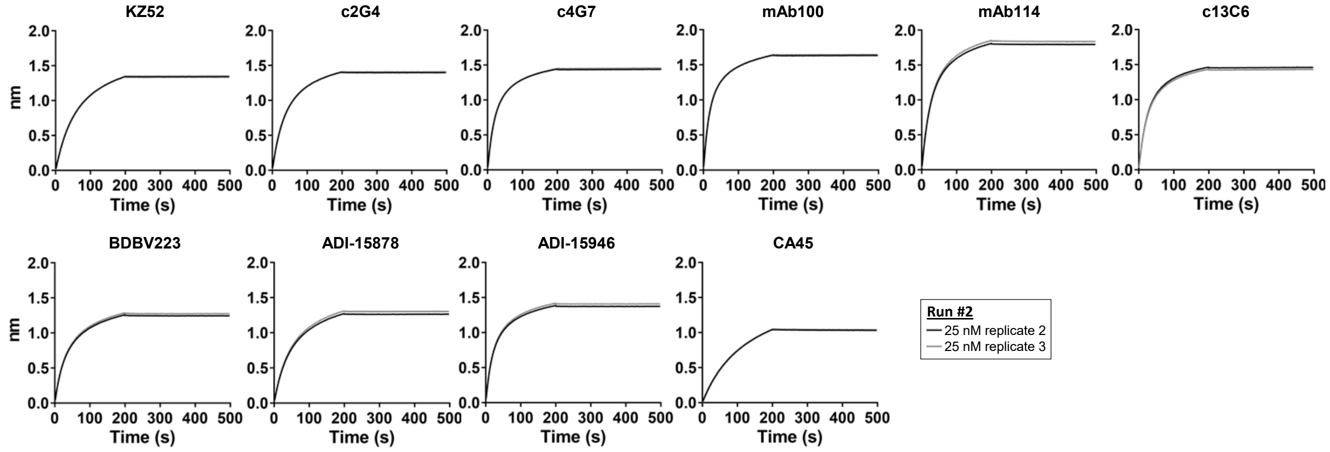

#### Supplementary Fig. 6

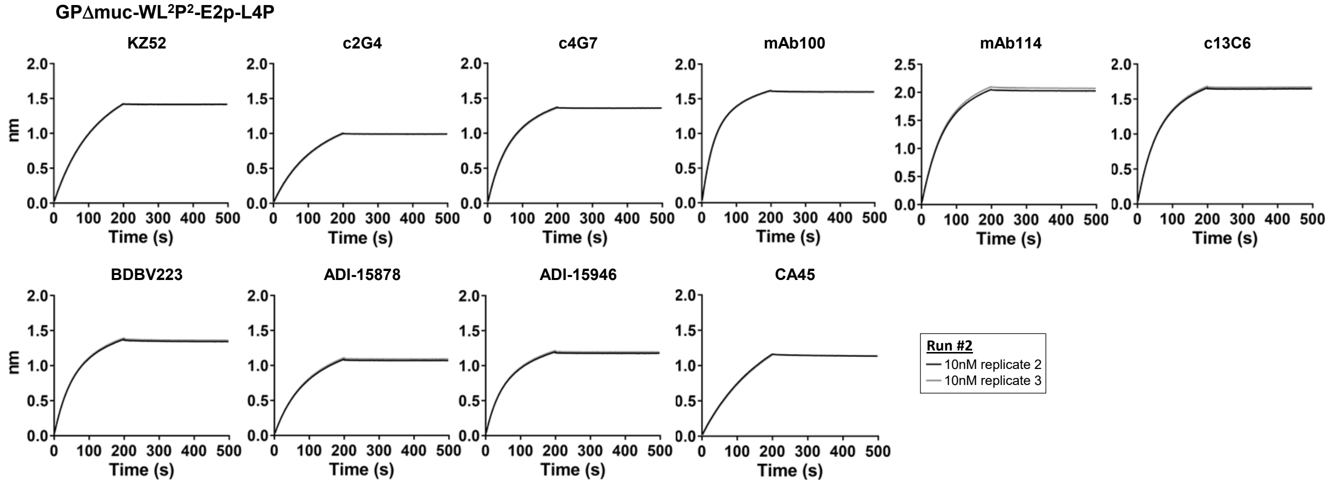


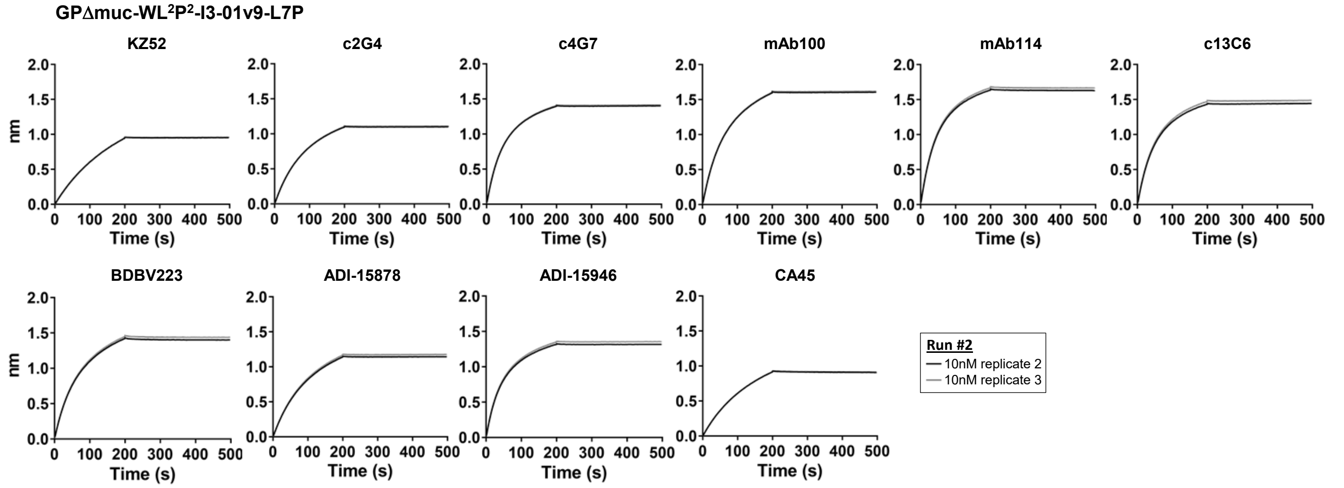


**f** Standard deviation of antibody binding signals at the highest antigen concentration (3 replicates)

| EBOV GP constructs: | KZ52 | c2G4 | c4G7 | mAb100 | mAb114 | c13C6 | BDBV223 | ADI-15878 | ADI-15946 | CA45 |
|---|---|---|---|---|---|---|---|---|---|---|
| GPΔmuc-WL$^2$P$^2$-foldon | 0.02 | 0.03 | 0.07 | 0.01 | 0.06 | 0.04 | 0.07 | 0.02 | 0.01 | 0.00 |
| GPΔmuc-WL$^2$P$^2$-5GS-FR | 0.04 | 0.09 | 0.02 | 0.03 | 0.03 | 0.03 | 0.06 | 0.02 | 0.04 | 0.06 |
| GPΔmuc-WL$^2$P$^2$-E2p-L4P | 0.02 | 0.04 | 0.02 | 0.05 | 0.03 | 0.02 | 0.05 | 0.03 | 0.04 | 0.02 |
| GPΔmuc-WL$^2$P$^2$-10GS-I3-01v9-L7P | 0.08 | 0.06 | 0.06 | 0.01 | 0.03 | 0.05 | 0.03 | 0.06 | 0.05 | 0.06 |

**g** mAb100-purified $GP_{ECTO}$-10GS-FR

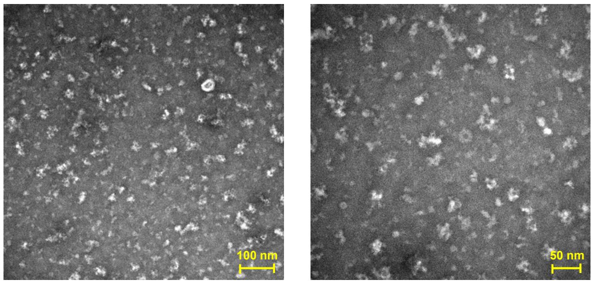

**Supplementary Fig. 6. Design and characterization of multilayered EBOV GPΔmuc-presenting NPs.** (**a**) Locking domains (LD) selected from 815 homodimers in the PDB for stabilizing the NP-forming interface. PDB IDs, number of residues after N- and C-terminal deletion, and amino acid sequences are provided. (**b**)-(**c**) Antigenic evaluation by ELISA. Binding of FR and reengineered 60-meric E2p (E2p-LD4-PADRE, or E2p-L4P) and I3-01v9 (I3-01v9-LD7-PADRE, or I3-01v9-L7P) NPs that present a stabilized GPΔmuc-WL$^2$P$^2$ trimer to 10 antibodies with ELISA curves shown in (**b**) and $EC_{50}$ values (μg/ml) summarized in (**c**). All GP-NPs were expressed transiently in ExpiCHO cells, purified by a mAb100 column, and further purified by SEC on a Superose 6 10/300 GL column. (**d**) Antigenic evaluation of GPΔmuc-WL$^2$P$^2$-foldon and three NPs (FR, E2p-L4P, and I3-01v9-L7P) that present GPΔmuc-WL$^2$P$^2$ using BLI and 10 representative antibodies. This experiment is designated "Run #1". Sensorgrams were obtained from an Octet RED96 instrument using six concentrations (400-12.5 nM, 25-0.78 nM, and 10-0.31 nM by twofold dilution for GPΔmuc, FR, and E2p-L4p/I3-01v9-L7P, respectively) and AHQ biosensors (see Methods). I3-01v9 is a variant of I3-01 with a redesigned NP-forming interface based on the PDB structure (1VLW). I3-01v9 was reported in our previous study (ref. 55) with a construct name "1VLW-v9". (**e**) Antigenic evaluation of GPΔmuc-WL$^2$P$^2$-foldon and three NPs (FR, E2p-L4P, and I3-01v9-L7P) that present GPΔmuc-WL$^2$P$^2$ using BLI and 10 representative antibodies at the highest antigen concentration in (**d**) with two replicates. This experiment is designated "Run #2". (**f**) Standard deviation of antibody binding signals at the highest concentration using 3 replicates from Run #1 and Run #2. Color coding is based on the range of binding signals shown in Fig. 4j. (**g**) Representative micrographs from the negative-stain EM of $GP_{ECTO}$-10GS-FR following ExpiCHO expression and mAb114 purification. The use of mAb114 instead of mAb100 for IAC purification was based on the consideration that the MLD may occlude antibody access to IFL on the FR NP surface. Source data are provided as a Source Data file.

a

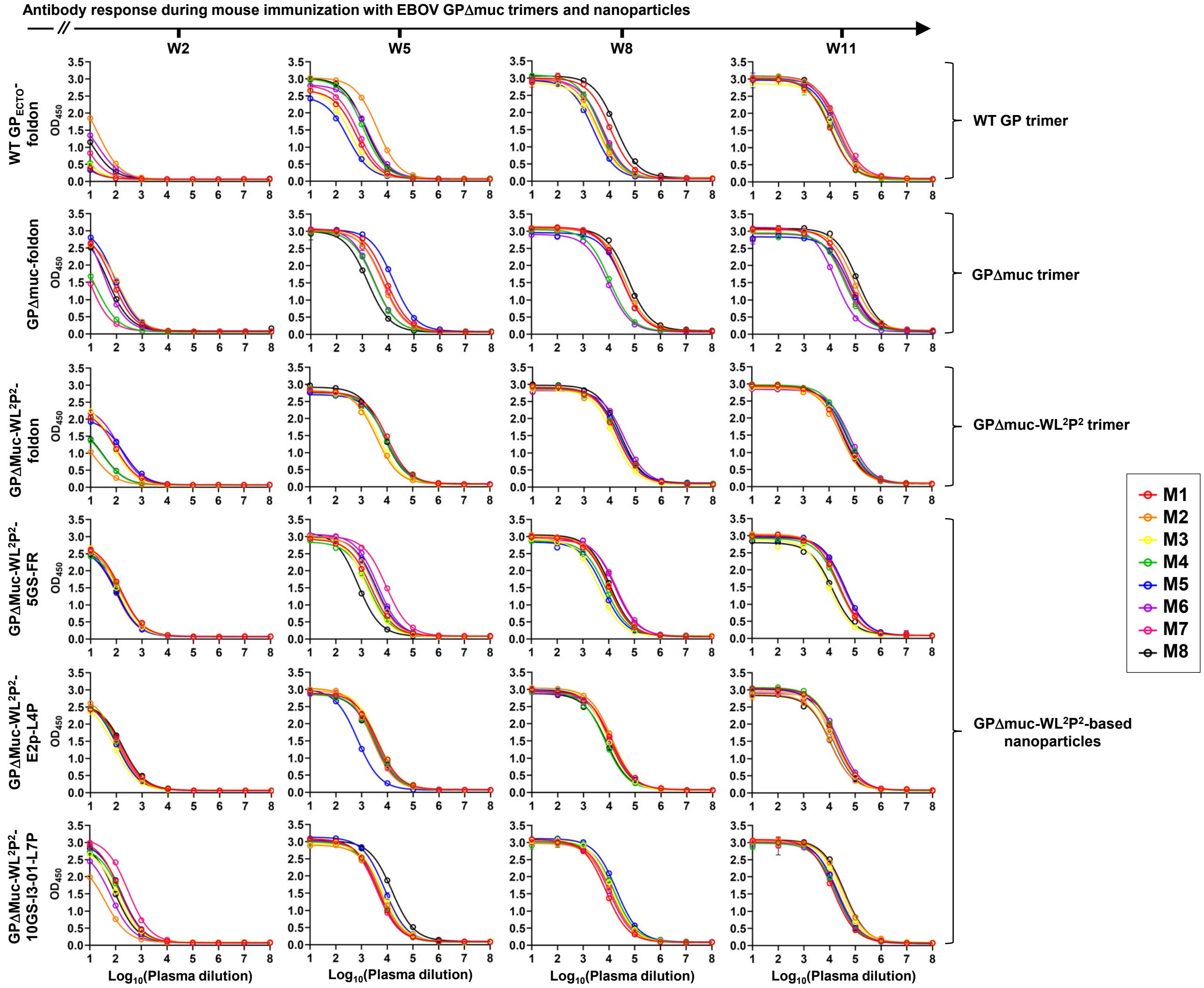


b

**EC$_{50}$ titers (fold of dilution) of mouse sera binding to EBOV GP trimers[a]**

| | W2 | | | W5 | | | W8 | | | W11 | | |
|---|---|---|---|---|---|---|---|---|---|---|---|---|
| | WT GP$_{ECTO}$-foldon | GPΔmuc-foldon | GPΔmuc-WL²P²-foldon | WT GP$_{ECTO}$-foldon | GPΔmuc-foldon | GPΔmuc-WL²P²-foldon | WT GP$_{ECTO}$-foldon | GPΔmuc-foldon | GPΔmuc-WL²P²-foldon | WT GP$_{ECTO}$-foldon | GPΔmuc-foldon | GPΔmuc-WL²P²-foldon |
| M1 | <1 | 71 | 93 | 607 | 8377 | 10280 | 10740 | 32034 | 18532 | 10467 | 56041 | 36300 |
| M2 | 21 | 119 | 14 | 4050 | 5879 | 3894 | 3663 | 40677 | 20082 | 20353 | 83513 | 28303 |
| M3 [b] | 5 | – | 67 | 376 | – | 4851 | 3323 | – | 14595 | 16418 | – | 37914 |
| M4 | 1 | 16 | 31 | 1183 | 2755 | 8470 | 4338 | 10690 | 19570 | 11262 | 37135 | 49211 |
| M5 | 3 | 89 | 187 | 289 | 15768 | 8994 | 2382 | 35240 | 22712 | 16546 | 56785 | 46695 |
| M6 | 20 | 31 | 127 | 1804 | 2752 | 8816 | 5651 | 9447 | 35455 | 18888 | 16351 | 64055 |
| M7 [b] | 5 | 7 | – | 740 | 6293 | – | 5066 | 33547 | – | 26293 | 42415 | – |
| M8 | 12 | 46 | 31 | 1449 | 1591 | 7248 | 17622 | 56265 | 25518 | 20626 | 117745 | 33217 |

[a] The EC$_{50}$ values were calculated from the besting fitting in GraphPad Prism 8.
[b] "-" indicates animal death during the immunization study.

**c**

**$EC_{50}$ titers (fold of dilution) of mouse sera binding to EBOV GPΔmuc-WL²P²-presenting nanoparticles [a]**

| | W2 | | | W5 | | | W8 | | | W11 | | |
|---|---|---|---|---|---|---|---|---|---|---|---|---|
| | GPΔmuc-WL²P²-5GS-FR | GPΔmuc-WL²P²-E2p-L4P | GPΔmuc-WL²P²-10GS-I3-01v9-L7P | GPΔmuc-WL²P²-5GS-FR | GPΔmuc-WL²P²-E2p-L4P | GPΔmuc-WL²P²-10GS-I3-01v9-L7P | GPΔmuc-WL²P²-5GS-FR | GPΔmuc-WL²P²-E2p-L4P | GPΔmuc-WL²P²-10GS-I3-01v9-L7P | GPΔmuc-WL²P²-5GS-FR | GPΔmuc-WL²P²-E2p-L4P | GPΔmuc-WL²P²-10GS-I3-01v9-L7P |
| M1 | 153 | 172 | 148 | 2367 | 4154 | 4389 | 9932 | 11715 | 7637 | 24325 | 17631 | 13984 |
| M2 | 146 | 112 | 41 | 3115 | 2559 | 5927 | 10324 | 12671 | 11689 | 22640 | 11150 | 34464 |
| M3 | 102 | 86 | 115 | 1691 | 4077 | 7196 | 4239 | 8701 | 14184 | 10166 | 15309 | 26621 |
| M4 | 159 | 121 | 159 | 2053 | 3426 | 4860 | 7750 | 6774 | 16144 | 23135 | 18605 | 18439 |
| M5 | 99 | 106 | 165 | 3598 | 692.1 | 8006 | 6227 | 9166 | 19698 | 38846 | 13793 | 19315 |
| M6 [b] | 84 | 105 | 67 | 4238 | 2973 | 5440 | 17428 | 12140 | 10331 | – | 24189 | 17649 |
| M7 | 101 | 142 | 305 | 9029 | 3443 | 3872 | 20184 | 10342 | 11975 | 34598 | 15778 | 21201 |
| M8 | 104 | 178 | 93 | 725.6 | 3022 | 15325 | 11421 | 7550 | 16234 | 13170 | 11012 | 37636 |

[a] The $EC_{50}$ values were calculated from the besting fitting in GraphPad Prism 8.
[b] "-" indicates animal death during the immunization study.

**d** Control antibody neutralization of EBOV-Makona in 293T cells

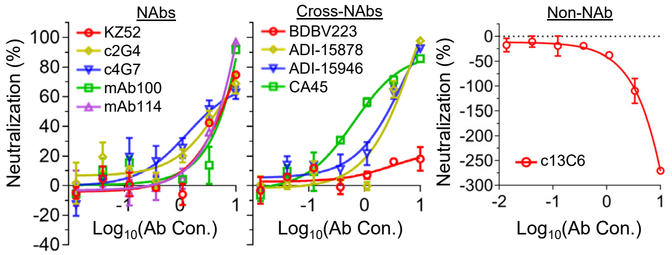


**e** Control antibody neutralization of BDBV-Uganda in 293T cells

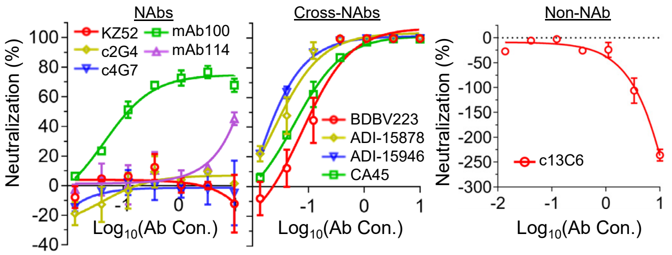


**f** Control antibody neutralization of EBOV-Makona in TZM-bl cells

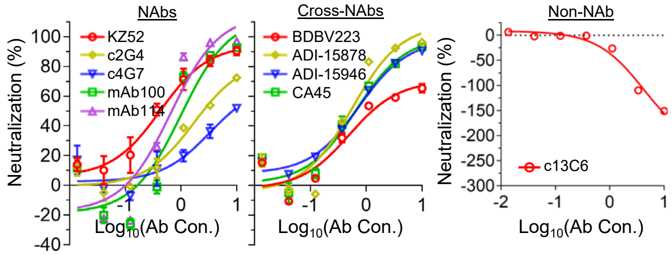


**g** Control antibody neutralization of MLV in 293T cells

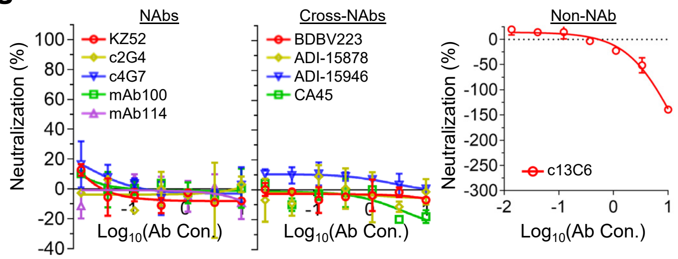


**h** Mouse IgG neutralization of EBOV-Makona in 293T cells

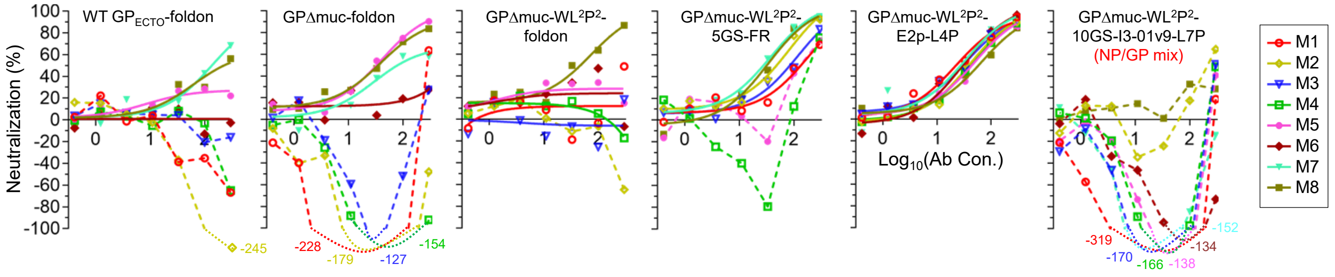


**i** Mouse IgG neutralization of BDBV-Uganda in 293T cells

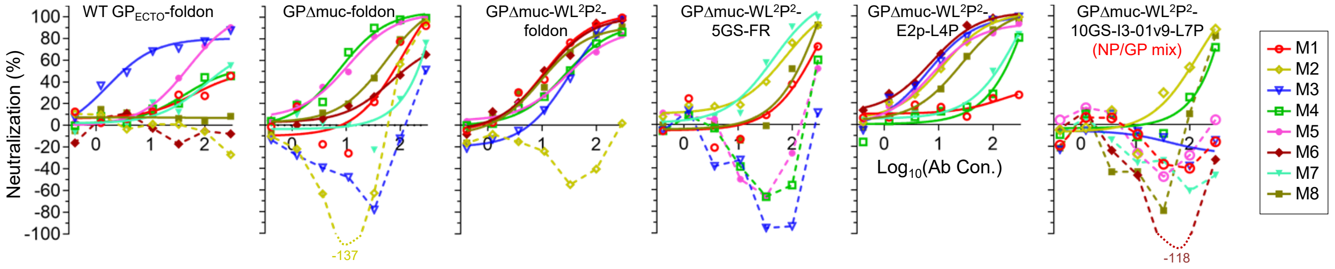

**j**

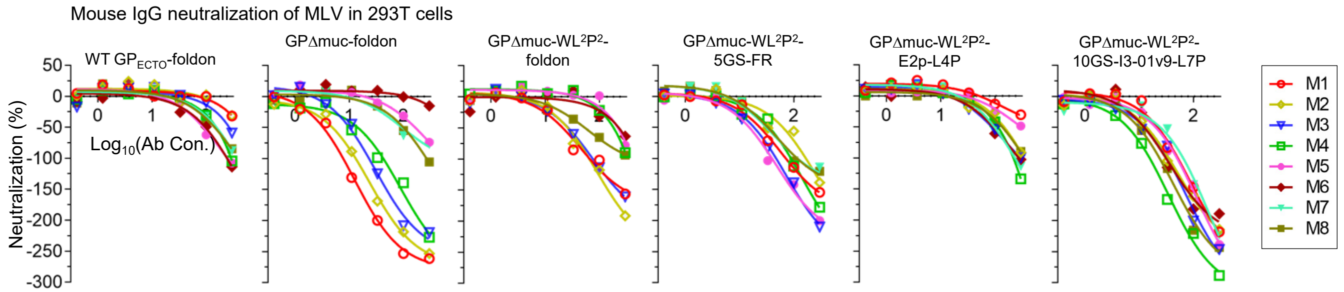


**Supplementary Fig. 7. Immunogenicity of EBOV GP/GPΔmuc trimers and GPΔmuc-presenting NPs in BALB/c mice.** (**a**) ELISA binding curves of mouse plasma from three trimer groups, in which mice were immunized with WT $GP_{ECTO}$-foldon, GPΔmuc-foldon, and GPΔmuc-$WL^2P^2$-fold trimers, and three NP groups, in which mice were immunized with FR, E2p-L4P, and I3-01v9-L7P NPs presenting GPΔmuc-$WL^2P^2$, to GPΔmuc-$WL^2P^2$ at w2, w5, w8, and w11. (**b**) $EC_{50}$ titers measured for the three trimer groups. (**c**) $EC_{50}$ titers measured for the three NP groups. The $EC_{50}$ titer was measured in the unit of fold of dilution. Of note, plasma binding at w2 did not reach the plateau (or saturation) to allow for accurate determination of $EC_{50}$ titers. Nonetheless, the $EC_{50}$ values calculated in Prism were used as a quantitative measure of binding antibody titers to facilitate the comparison of different vaccine groups at w2. (**d**) and (**e**) Ebolavirus-pp (EBOV-Makona and BDBV-Uganda) neutralization by 10 representative antibodies in HEK293T cells. (**f**) Ebolavirus-pp (EBOV-Makona) neutralization by 10 representative antibodies in TZM-bl cells. (**g**) MLV-pp neutralization by 10 representative antibodies in HEK293T cells as a negative control. In (**d**)-(**g**), the antibody neutralization was performed in duplicates, with mean value and standard deviation (SD) shown as symbol and solid line, respectively. (**h**) and (**i**) Ebolavirus-pp (EBOV-Makona and BDBV-Uganda) neutralization by purified mouse IgG from six vaccine groups in HEK293T cells, with (**j**) MLV-pp included as a negative control. The IgG concentration was started at 300μg/ml followed by a series of threefold dilutions. Due to the limited availability of mouse samples, ebolavirus-pp assays were performed only once without duplicate. For mice showing enhanced ebolavirus-pp infection (e.g. %neuralization<-30.0%), data points are shown as dashed lines instead of fitted curves. Data points below -100% are excluded with the lowest %neutralization values labeled on the plot and color-coded according to the mouse index. Mouse IgGs were purified from plasma collected at the last time point, w11. Source data are provided as a Source Data file.

### Supplementary Fig. 8

**a**

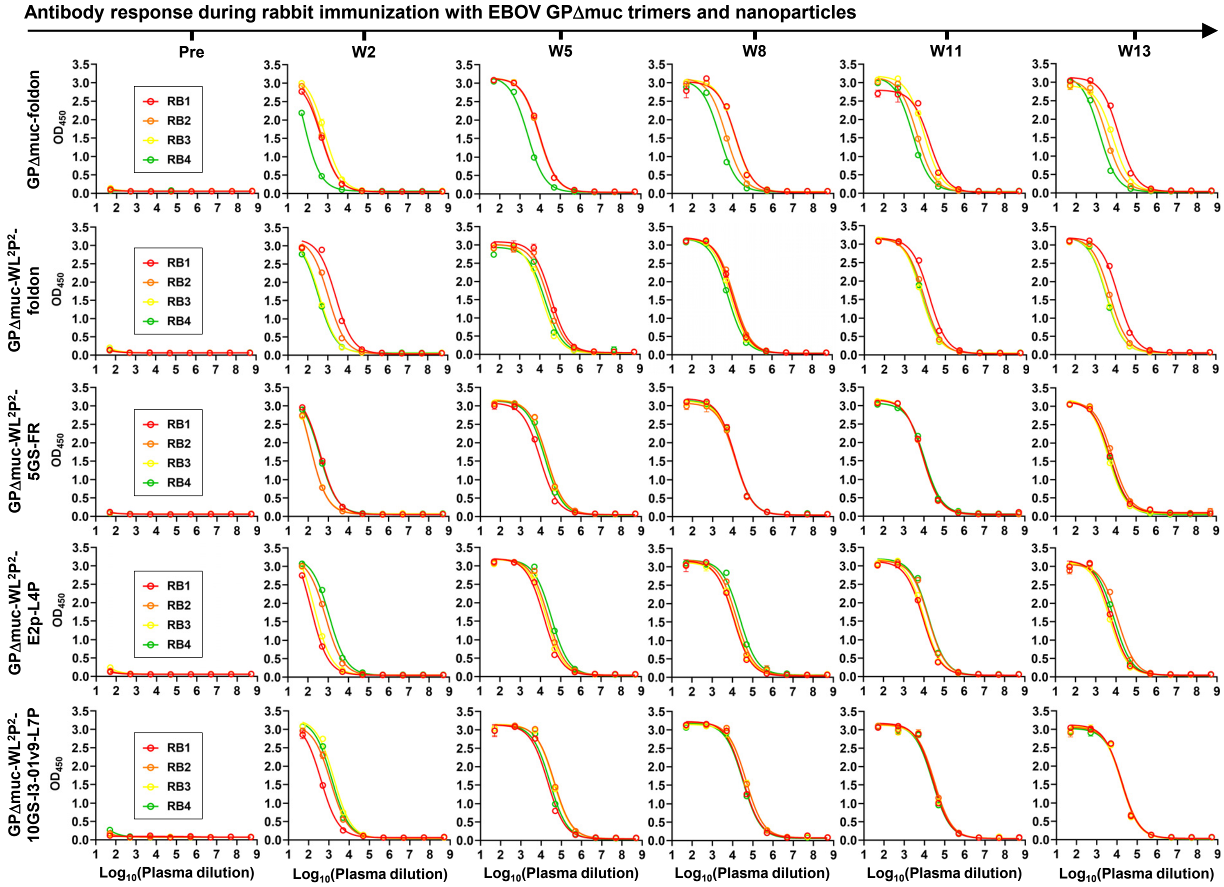


**b**

**EC$_{50}$ titers (fold of dilution) of rabbit sera binding to EBOV GP trimers[a]**

| | W2 | | W5 | | W8 | | W11 | | W13 | |
|---|---|---|---|---|---|---|---|---|---|---|
| | GPΔmuc-foldon | GPΔmuc-WL²P²-foldon | GPΔmuc-foldon | GPΔmuc-WL²P²-foldon | GPΔmuc-foldon | GPΔmuc-WL²P²-foldon | GPΔMuc-foldon | GPΔmuc-WL²P²-foldon | GPΔmuc-foldon | GPΔmuc-WL²P²-foldon |
| RB1 | 464 | 2428 | 9404 | 35115 | 13868 | 10110 | 18135 | 16116 | 12777 | 13338 |
| RB2 | 456 | 1044 | 8904 | 26144 | 4647 | 12073 | 4446 | 8694 | 3143 | 5177 |
| RB3 | 701 | 378 | 8943 | 13273 | 12652 | 8363 | 9807 | 6517 | 6681 | 3586 |
| RB4 | 57 | 336 | 2392 | 18008 | 2220 | 6092 | 2663 | 6901 | 1443 | 3424 |

[a] The EC$_{50}$ values were calculated from the besting fitting in GraphPad Prism 8.4.3.

**EC$_{50}$ titers (fold of dilution) of mouse sera binding to EBOV WT GPΔmuc-presenting nanoparticles [a]**

| | W2 | | | W5 | | | W8 | | | W11 | | |
|---|---|---|---|---|---|---|---|---|---|---|---|---|
| | GPΔmuc-5GS-FR | GPΔmuc-E2p-L4P | GPΔmuc-10GS-I3-01v9-L7P | GPΔmuc-5GS-FR | GPΔmuc-E2p-L4P | GPΔmuc-10GS-I3-01v9-L7P | GPΔmuc-5GS-FR | GPΔmuc-E2p-L4P | GPΔmuc-10GS-I3-01v9-L7P | GPΔmuc-5GS-FR | GPΔmuc-E2p-L4P | GPΔmuc-10GS-I3-01v9-L7P |
| RB1 | 393 | 133 | 404 | 9406 | 14736 | 21261 | 12961 | 11405 | 32895 | 9041 | 8972 | 26363 |
| RB2 | 117 | 735 | 1207 | 20093 | 23950 | 44607 | 12927 | 15609 | 45311 | 8893 | 16560 | 31073 |
| RB3 | 120 | 201 | 1774 | 18894 | 19727 | 41440 | 12770 | 12457 | 47664 | 8644 | 9954 | 28526 |
| RB4 | 368 | 1126 | 1382 | 16116 | 34695 | 27116 | 13140 | 22771 | 32291 | 10558 | 16803 | 25205 |

| | W13 | | |
|---|---|---|---|
| | GPΔmuc-5GS-FR | GPΔmuc-E2p-L4P | GPΔmuc-10GS-I3-01v9-L7P |
| RB1 | 5272 | 5795 | 16907 |
| RB2 | 6817 | 12387 | 18398 |
| RB3 | 4146 | 4889 | 16645 |
| RB4 | 5047 | 8066 | 18753 |

The EC$_{50}$ values were calculated from the besting fitting in GraphPad Prism 8.4.3.

**Supplementary Fig. 8**

**c** **Ebolavirus-pp neutralization by purified rabbit IgG at the 5th time point (w11)**

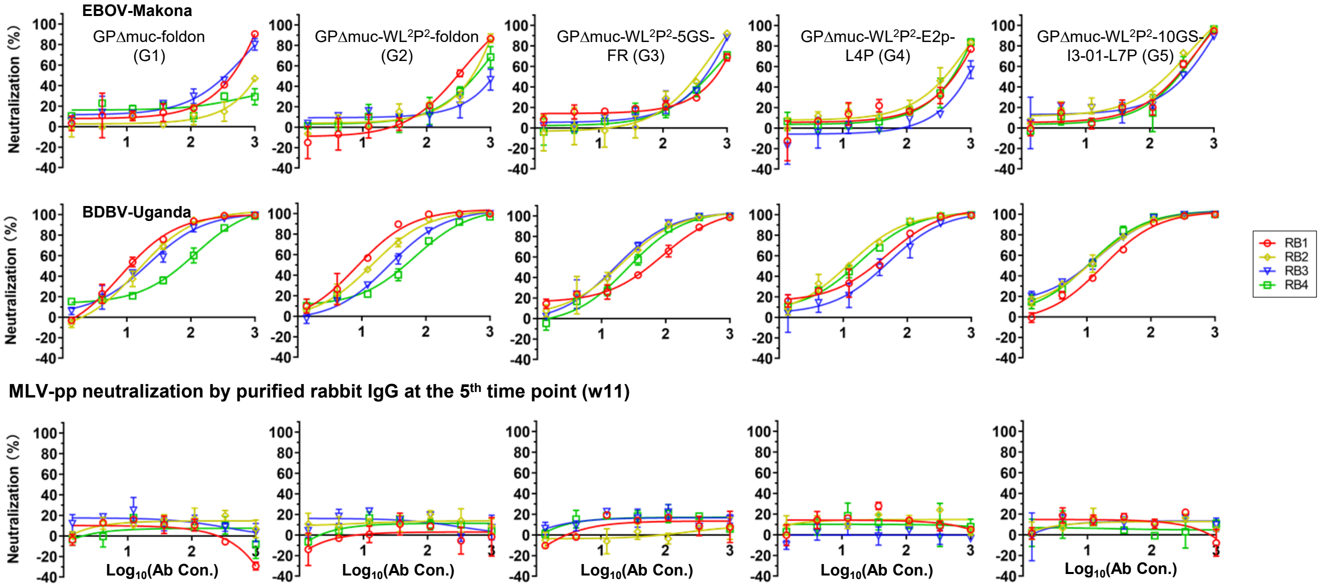


**d** **Ebolavirus-pp neutralization by purified rabbit IgG at the 1st time point (Pre)**

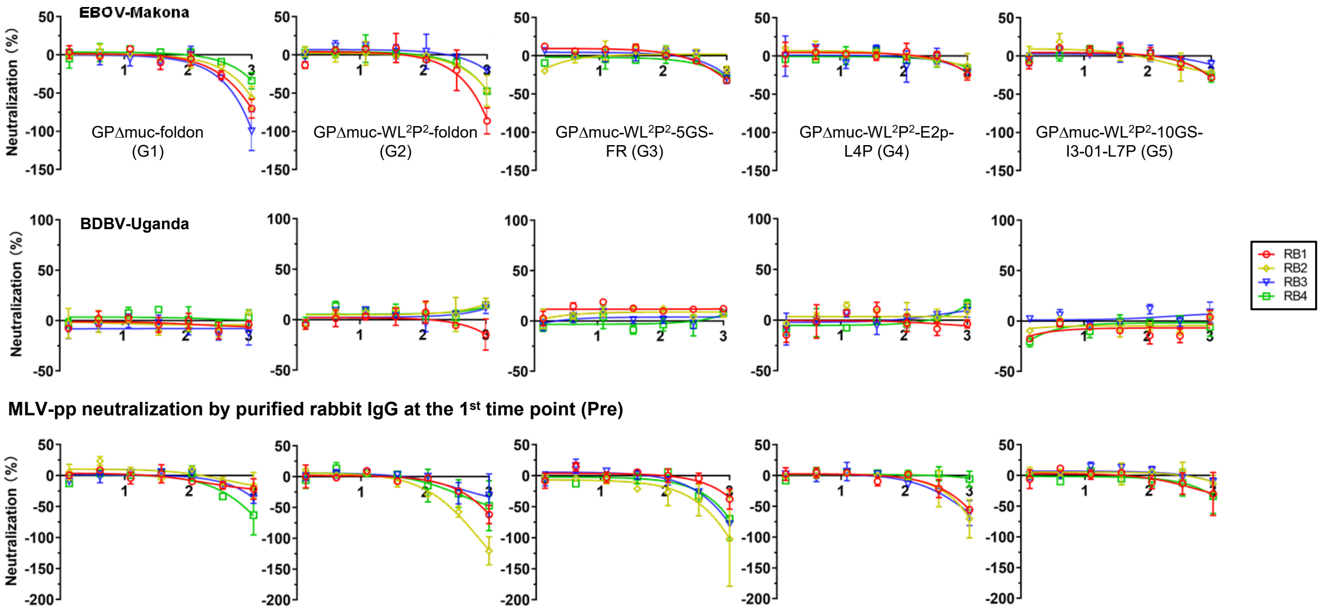

#### Supplementary Fig. 8

**e** **Ebolavirus-pp neutralization by purified rabbit IgG at the 2[nd] time point (w2)**

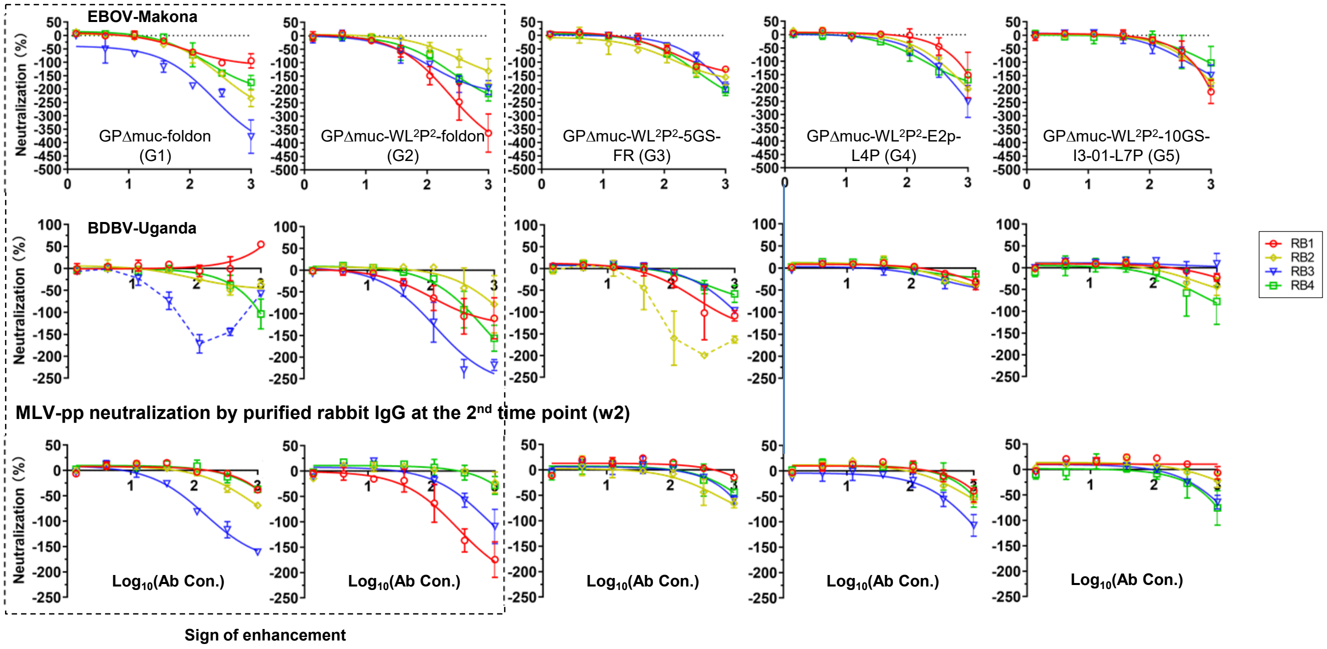


**f** **Ebolavirus-pp neutralization by purified rabbit IgG at the 3[rd] time point (w5)**

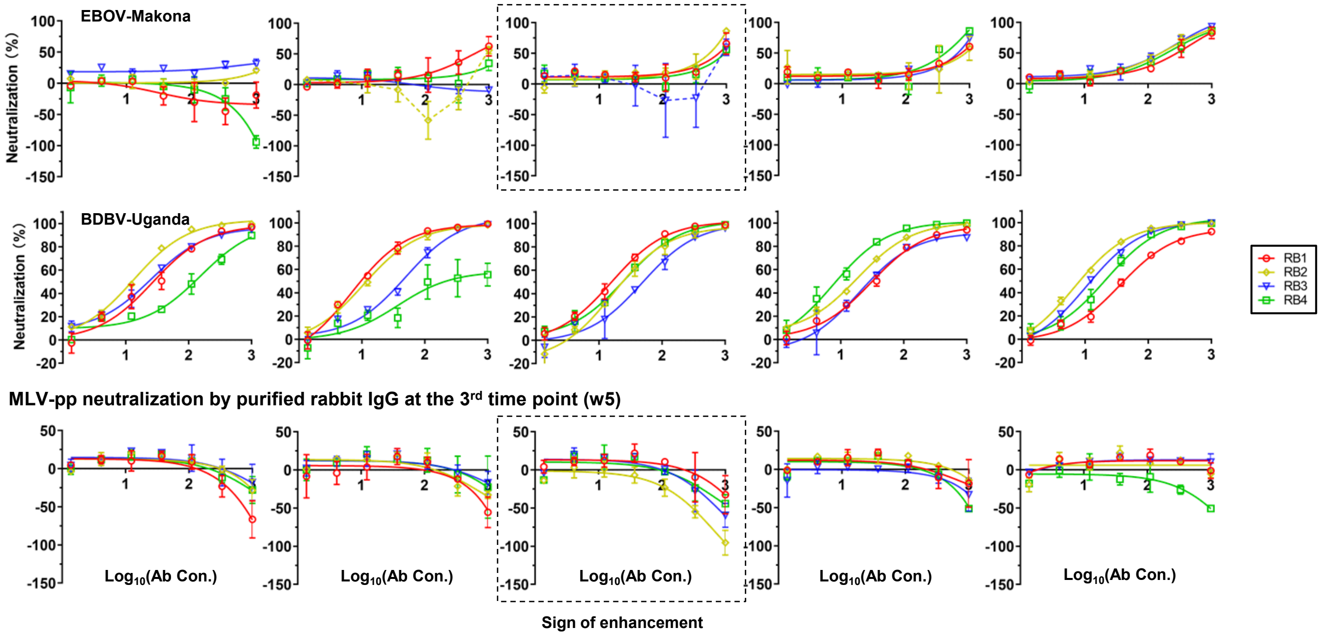

**g** Ebolavirus-pp neutralization by purified rabbit IgG at the 4th time point (w8)

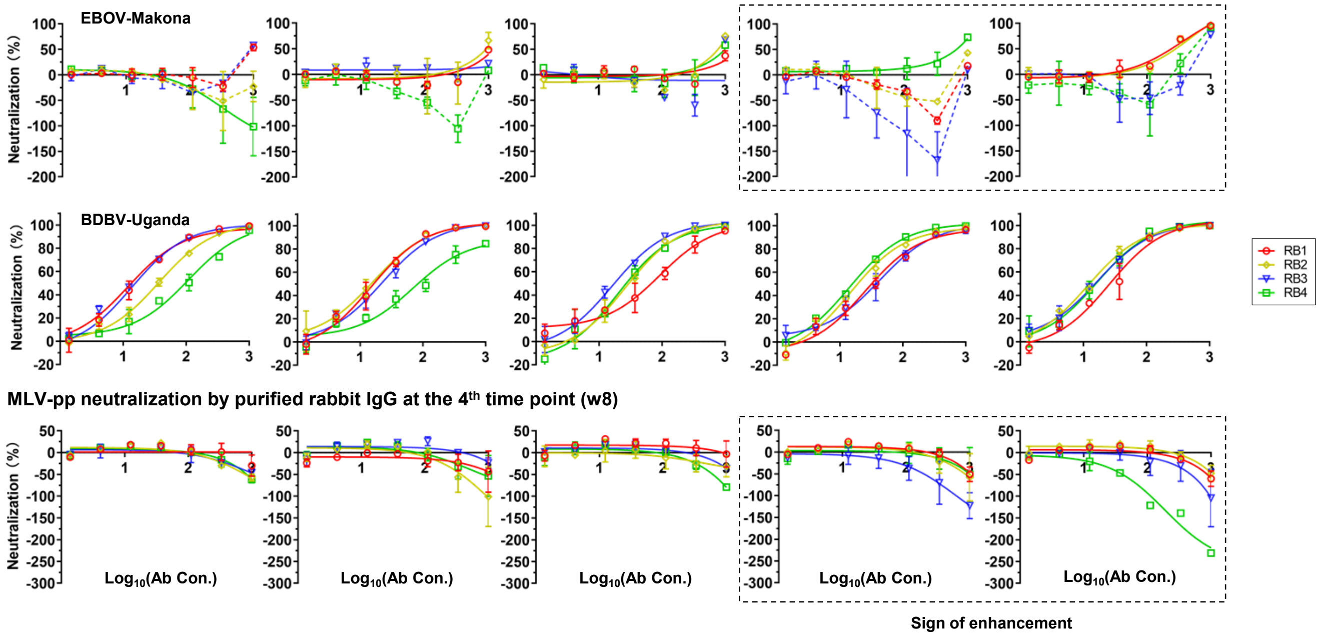


**Supplementary Fig. 8. Immunogenicity of EBOV GP/GPΔmuc trimers and GPΔmuc-presenting NPs in rabbits.** (**a**) ELISA binding curves of rabbit plasma from two trimer groups, in which rabbits were immunized with GPΔmuc-foldon and GPΔmuc-WL$^2$P$^2$-fold trimers, and three NP groups, in which rabbits were immunized with FR, E2p-L4P, and I3-01v9-L7P NPs presenting GPΔmuc-WL$^2$P$^2$, to GPΔmuc-WL$^2$P$^2$ at w0, w2, w5, w8, w11, and w13. (**b**) $EC_{50}$ titers measured for the two trimer groups and three NP groups. The $EC_{50}$ titer was measured in the unit of fold of dilution. Of note, plasma binding at w2 did not reach the plateau (or saturation), but the $OD_{450}$ values were approximately at the same level as those obtained at w5 (fully plateaued). Therefore, the $EC_{50}$ values calculated in Prism were considered sufficiently accurate to facilitate the comparison of different vaccine groups at w2. (**c**) Ebolavirus-pp (EBOV-Makona and BDBV) and MLV-pp neutralization by purified rabbit IgGs from w11 in HEK293T cells. No sign of enhanced pseudovirus infection is observed. Ebolavirus-pp (EBOV-Makona and BDBV) and MLV-pp neutralization by purified rabbit IgG from Pre, w2, w5, and w8 in HEK293T cells is shown in (**d**), (**e**), (**f**) and (**g**). In (**c**)-(**g**), the rabbit IgG neutralization was performed in duplicates, with mean value and standard deviation (SD) shown as symbol and solid line, respectively. Data points that do not fit a sigmodal dose response model are shown as dashed lines. Signs of enhanced pseudovirus infection for the two GPΔmuc trimer groups, the FR group, and the two multilayered 60-meric NP groups are observed at w2, w5, and w8, respectively, and are highlighted in dashed rectangles. Source data are provided as a Source Data file.

**a**

**GPΔmuc-WL²P²-1TD0-Avi-Biot**

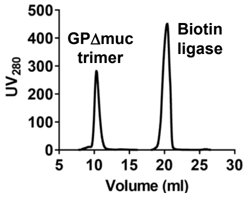


**b**

**Gating strategies in bulk B cell sorting**

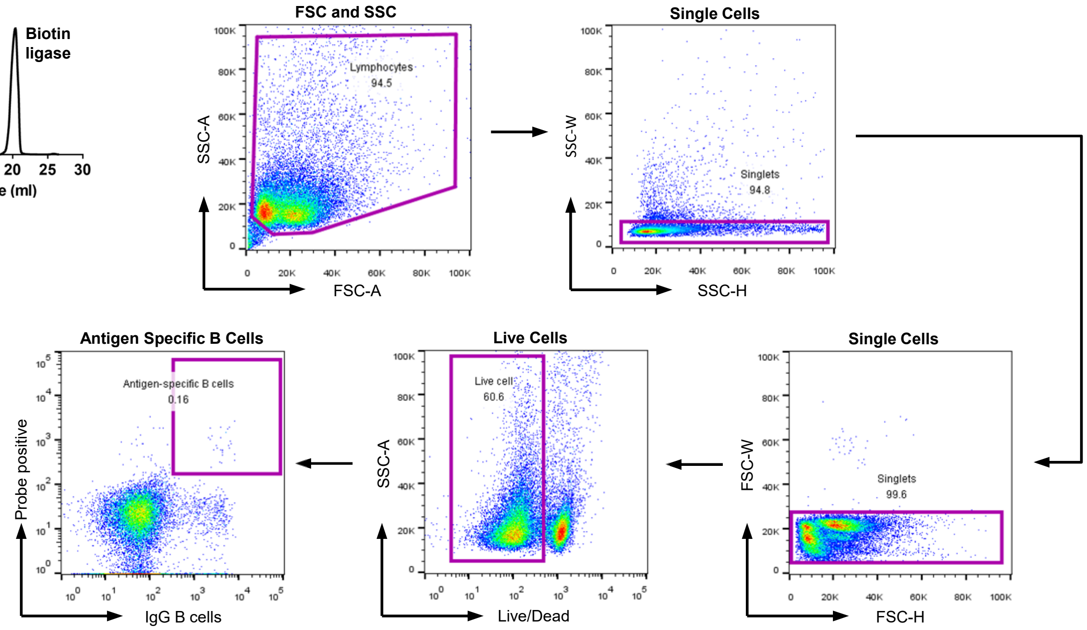


**EBOV GPΔmuc-specific sorting of mouse splenic B cells from five vaccine groups**

| GPΔmuc-Foldon | | | GPΔmuc-WL²P²-Foldon | | | GPΔmuc-WL²P²-5GS-FR | | | GPΔmuc-WL²P²-E2p-L4P | | | GPΔmuc-WL²P²-10GS-I3-01v9-L7P | | |
|---|---|---|---|---|---|---|---|---|---|---|---|---|---|---|
| Mouse | Sorted B cells | %Splenic B cells | Mouse | Sorted B cells | %Splenic B cells | Mouse | Sorted B cells | %Splenic B cells | Mouse | Sorted B cells | %Splenic B cells | Mouse | Sorted B cells | %Splenic B cells |
| G5-1 | 760 | 0.03579 | G7-1 | 1134 | 0.07849 | G3-1 | 1856 | 0.07481 | G4-1 | 1360 | 0.07285 | G10-2 | 401 | 0.07874 |
| G5-3 | 582 | 0.04646 | G7-3 | 585 | 0.18874 | G3-2 | 1146 | 0.04230 | G4-2 | 906 | 0.09027 | G10-3 | 602 | 0.10956 |
| G5-6 | 857 | 0.04580 | G7-4 | 1043 | 0.18983 | G3-5 | 1362 | 0.04844 | G4-3 | 1369 | 0.08957 | G10-4 | 443 | 0.06507 |
| G5-7 | 1065 | 0.04149 | G7-6 | 949 | 0.19321 | G3-7 | 2864 | 0.10775 | G4-6 | 1400 | 0.08291 | G10-5 | 871 | 0.04330 |
| G5-8 | 752 | 0.04904 | G7-8 | 982 | 0.09138 | G3-8 | 843 | 0.06591 | G4-7 | 1485 | 0.05536 | G10-8 | 372 | 0.05807 |

**c**

**Next-generation sequencing (NGS) data obtained for EBOV GP-specific mouse splenic B cells [a]**

| Vaccine antigen | Mouse | Chain | $N_{Chain}$ | $N_{Step5}$ | <Length> | $N_{Usable}$ |
|---|---|---|---|---|---|---|
| | G5-1 | H [b] | 241,888 | 152,926 | 369.7 | 83,074 |
| | | K | 320,999 | 271421 | 335.3 | 244,765 |
| | G5-3 | H [b] | 151,041 | 38,965 | 365.7 | 25,901 |
| | | K | 254,461 | 190,810 | 332.5 | 159,340 |
| GPΔmuc -foldon | G5-6 | H | 164,049 | 72,966 | 364.3 | 54,123 |
| | | K | 149,922 | 122,358 | 328.8 | 112,897 |
| | G5-7 | H [b] | 127,583 | 31,333 | 362.8 | 21,695 |
| | | K | 132,822 | 101,243 | 331.6 | 86,936 |
| | G5-8 | H [b] | 416,364 | 146,889 | 363.5 | 98,999 |
| | | K | 319,964 | 245,757 | 331.5 | 219,409 |
| | G7-1 | H [b] | 372,642 | 174,424 | 363.5 | 116,260 |
| | | K | 147,494 | 121,251 | 330.8 | 107536 |
| | G7-3 | H [b] | 225,662 | 109,976 | 363.4 | 75,064 |
| | | K | 207,512 | 158,440 | 332.6 | 142,203 |
| GPΔmuc-WL²P² -foldon | G7-4 | H [b] | 195,714 | 58,996 | 364.4 | 39,423 |
| | | K | 181,907 | 143,388 | 333.8 | 126,382 |
| | G7-6 | H [b] | 203,093 | 49,877 | 361.0 | 34,366 |
| | | K | 234,578 | 193,315 | 334.7 | 174,967 |
| | G7-8 | H [b] | 193,745 | 96,042 | 352.4 | 75,369 |
| | | K | 166,563 | 138,767 | 334.7 | 118,193 |
| | G3-1 | H [b] | 181,414 | 15,426 | 361.6 | 9,391 |
| | | K | 172,971 | 110,166 | 333.9 | 99,231 |
| | G3-2 | H [b] | 196,037 | 24,375 | 366.4 | 15,433 |
| | | K | 165,297 | 118,826 | 332.0 | 106,667 |
| GPΔmuc-WL²P² -5GS-FR [c] | G3-5 | H [b] | 171,116 | 13,134 | 367.4 | 8,154 |
| | | K | 202,059 | 149,466 | 335.5 | 133,378 |
| | G3-7 | H [b] | 149,991 | 10,289 | 365.4 | 6,028 |
| | | K | 192,304 | 132,601 | 329.7 | 120,184 |
| | G3-8 | H | 250,631 | 171,124 | 362.3 | 132,011 |
| | | K | 234,322 | 162,071 | 328.2 | 139,171 |

| Vaccine antigen | Mouse | Chain | $N_{Chain}$ | $N_{Usable}$ | <Length> | $Perc_{Usable}$ |
|---|---|---|---|---|---|---|
| | G4-1 | H | 132,044 | 55,538 | 362.8 | 42,292 |
| | | K | 173,895 | 138,355 | 331.8 | 118,788 |
| | G4-2 | H | 95,696 | 71,107 | 360.5 | 57,045 |
| GPΔmuc-WL²P² -E2p-L4P | | K | 155,727 | 132,125 | 335.0 | 117,275 |
| | G4-3 | H | 105,559 | 85,392 | 362.1 | 72,646 |
| | | K | 75,330 | 59,105 | 332.3 | 48,150 |
| | G4-6 | H | 139,269 | 106,616 | 365.5 | 84,238 |
| | | K | 146,092 | 113,590 | 335.2 | 105,666 |
| | G4-7 | H [b, d] | 672,503 | 542,016 | 365.2 | 369,668 |
| | | K | 519,784 | 405,118 | 333.6 | 362,987 |
| | G10-2 | H | 452,882 | 268,275 | 365.5 | 212,617 |
| | | K | 290,177 | 258,708 | 328.5 | 233,447 |
| | G10-3 | H | 470,021 | 278,422 | 362.0 | 210,314 |
| GPΔmuc-WL²P² -10GS-I3-01v9 -L7P | | K | 223,350 | 207,358 | 333.2 | 191,472 |
| | G10-4 | H | 297,056 | 261,706 | 363.5 | 215,196 |
| | | K | 226,568 | 203,176 | 336.7 | 187,416 |
| | G10-5 | H | 190,727 | 138,889 | 366.4 | 106,419 |
| | | K | 237,420 | 197,478 | 331.3 | 183,103 |
| | G10-8 | H | 260,502 | 90,086 | 364.6 | 72,427 |
| | | K | 350,497 | 270,707 | 334.3 | 229,614 |

[a] Listed items include the vaccine antigen, mouse sample ID, number of VH/VK chains prior to the antibodyomics pipeline processing, number of VH/VK chains at step 5 of the pipeline processing, average read length, and number of usable chains after removing fragments with a V-gene alignment of 250bp or shorter.

[b] Due to the difficulty in library preparation, a smaller heavy chain dataset was obtained for a number of mouse samples, especially those from the trimer and FR groups. Two additional Ion S5 sequencing runs using the Ion 530 chip was performed to obtain more heavy chain reads to facilitate the profile analysis.

[c] G3-1, 5, and 7 showed especially low read-out and the profiles were only used for qualitative comparison with other vaccine groups but not for statistical analysis.

[d] Due to the large size of the dataset obtained from the additional NGS run, pre-filtering with a read length cutoff of 400bp was performed before merging it with the heavy chain dataset obtained from the initial run.

#### Supplementary Fig. 9

**d** GPΔmuc-foldon group

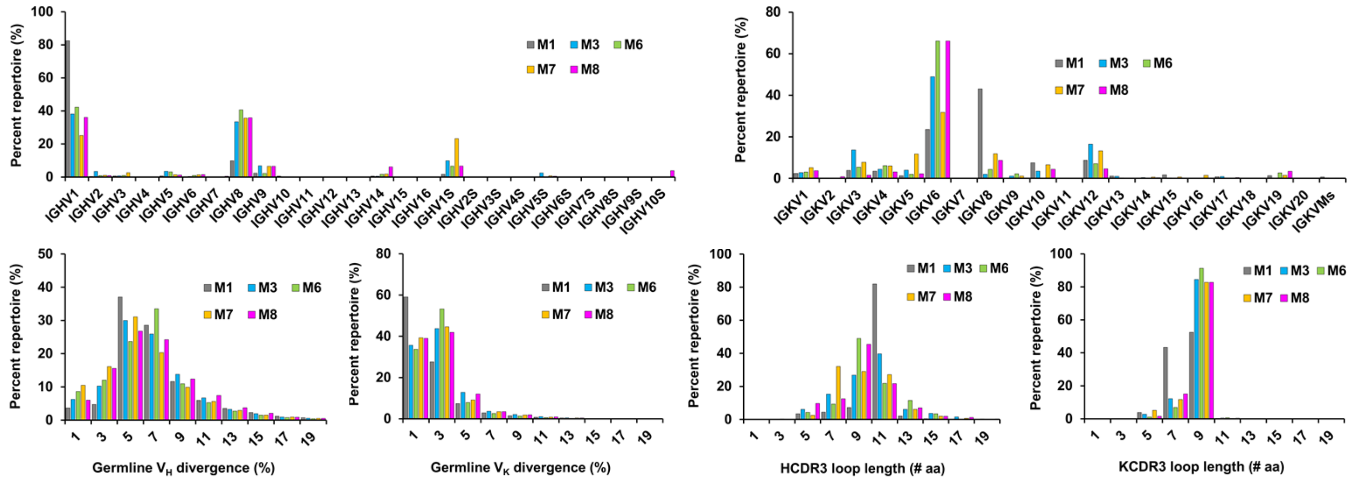


**e** GPΔmuc-WL²P²-5GS-FR group

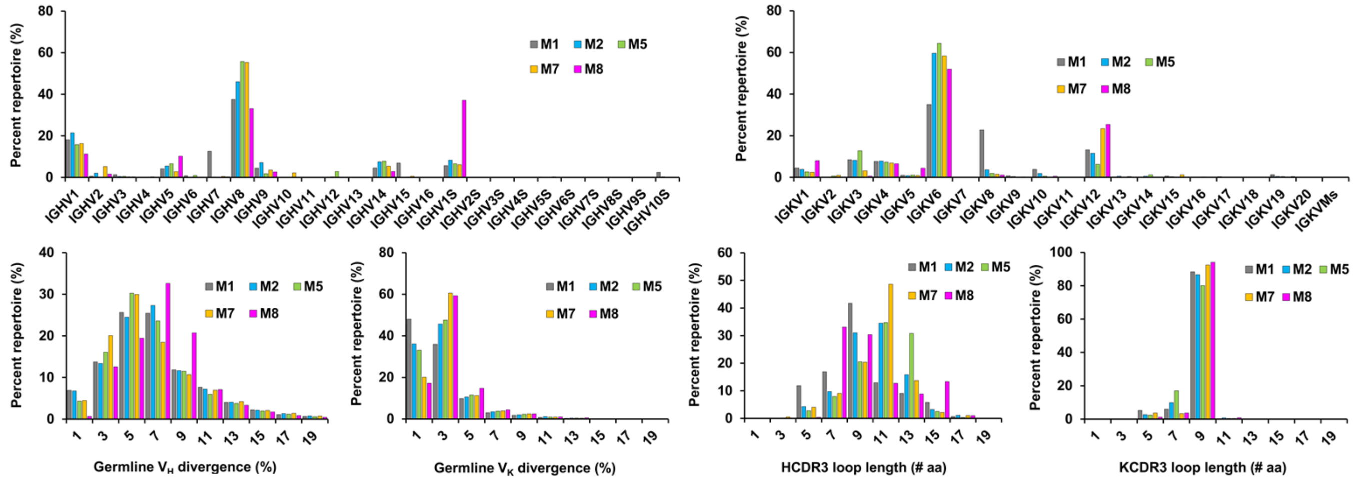


**f** GPΔmuc-WL²P²-10GS-I3-01v9-L7P group

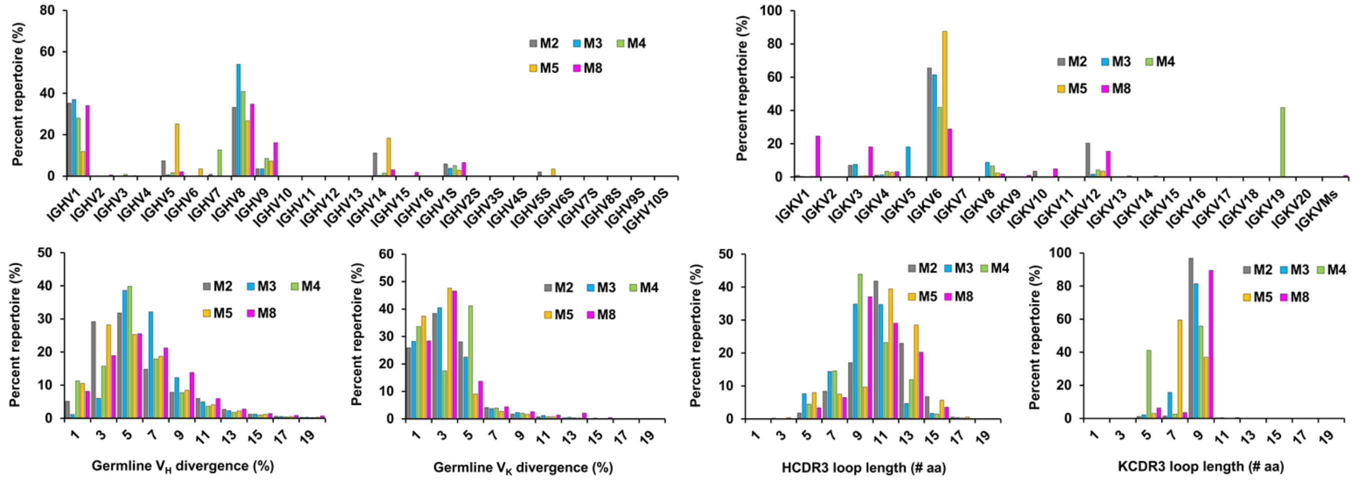

**Supplementary Fig. 9. Unbiased repertoire analysis of bulk-sorted EBOV GP-specific mouse splenic B cells.** (**a**) SEC profile of a biotinylated Avi-tagged EBOV GPΔmuc trimer, termed GPΔmuc-WL$^2$P$^2$-1TD0-Avi-Biot, obtained from a HiLoad Superdex 200 16/600 column. 1TD0 is a trimerization domain used to stabilize GPΔmuc and to mask foldon-specific B cells in the two trimer vaccine groups. (**b**) Top: gating strategies used in the antigen-specific mouse B cell sorting (Step 1: remove cell debris; Steps 2 and 3: exclude clumped or sticky cells to ensure that only single cells remain; Step 4: remove dead cells; Step 5: identify EBOV GP-specific B cells). Bottom: Summary of EBOV GPΔmuc-specific bulk sorting of mouse splenic B cells from five vaccine groups. (**c**) Antibodyomics analysis of repertoire NGS data obtained for GPΔmuc-specific mouse splenic B cells. Distribution of critical B-cell properties are shown for three vaccine groups, in which mice were immunized with (**d**) GPΔmuc-foldon, (**e**) GPΔmuc-WL$^2$P$^2$-5GS-FR, and (**f**) GPΔmuc-WL$^2$P$^2$-10GS-I3-01v9-L7P. For each group, distributions are shown for germline VH/VK gene usage (top), germline VH/VK divergence (bottom left) and CDRH/K3 loop length (bottom right). Source data are provided as a Source Data file.

**Supplementary Table 1. Primers used in EBOV GP construct design, nanoparticle construct design, and next-generation sequencing analysis of mouse splenic B cells.**

**a. Primers used in EBOV GP construct design**

| | |
|---|---|
| Leader-SalI-Forward | ACCGTCGTCGACGCCACCATGGGTGTCACCGGGATACTCCAG |
| GPΔmuc/$GP_{ECTO}$-NheI-BamHI-Reverse | ATGATGGGATCCCTATTAGCTAGCATCTACAAAGTCGTGGATGATCTGGTC |
| GPΔmuc-2WPZ-NheI-BamHI-Reverse | ATGATGGGATCCCTATTAGCTAGCTCGTTCGCCGACAAGTTTCTTGAG |
| GPΔmuc-L-NheI-BamHI-Reverse | ATGATGGGATCCCTATTAGCTAGCATCGGGCAATGTTTTATCTACAAA |
| GPΔmuc-Ext-NheI-BamHI-Reverse | ATGATGGGATCCCTATTAGCTAGCGTTATCGTTGTCTCCCTGATCGGGCAATGT |
| W615L-Forward | TGTTGCATAGAGCCGCACGATTTTACCAAGAATATCACCGATAAA |
| W615L-Reverse | TTTATCGGTGATATTCTTGGTGAGATCGTGCGGCTCTATGCAACA |
| GPΔmuc-P1-Forward | CTCCAGCTCTTCCTCAGGGCCCCTACGGAACTGCGAACGTTTAGC |
| GPΔmuc-P1-Reverse | GCTAAACGTTCGCAGTTCCGTAGGGGCCCTGAGGAAGAGCTGGAG |
| GPΔmuc-P2-Forward | CAGCTCTTCCTCAGGGCCACTCCGGAACTGCGAACGTTTAGCATT |
| GPΔmuc-P2-Reverse | AATGCTAAACGTTCGCAGTTCCGGAGTGGCCCTGAGGAAGAGCTG |
| GPΔmuc-P3-Forward | CAGCTCTTCCTCAGGGCCACTACGCCACTGCGAACGTTTAGCATTCTTAAT |
| GPΔmuc-P3-Reverse | ATTAAGAATGCTAAACGTTCGCAGTGGCGTAGTGGCCCTGAGGAAGAGCTG |
| GPΔmuc-P4-Forward | CTCTTCCTCAGGGCCACTACGGAACCGCGAACGTTTAGCATTCTTAATAGG |
| GPΔmuc-P4-Reverse | CCTATTAAGAATGCTAAACGTTCGCGGTTCCGTAGTGGCCCTGAGGAAGAG |
| GPΔmuc-P5-Forward | TTCCTCAGGGCCACTACGGAACTGCCAACGTTTAGCATTCTTAATAGGAAA |
| GPΔmuc-P5-Reverse | TTTCCTATTAAGAATGCTAAACGTTGGCAGTTCCGTAGTGGCCCTGAGGAA |
| GPΔmuc-P6-Forward | CTCAGGGCCACTACGGAACTGCGACCGTTTAGCATTCTTAATAGGAAAGCG |
| GPΔmuc-P6-Reverse | CGCTTTCCTATTAAGAATGCTAAACGGTCGCAGTTCCGTAGTGGCCCTGAG |
| GPΔmuc-P7-Forward | AGGGCCACTACGGAACTGCGAACGCCTAGCATTCTTAATAGGAAAGCGATA |
| GPΔmuc-P7-Reverse | TATCGCTTTCCTATTAAGAATGCTAGGCGTTCGCAGTTCCGTAGTGGCCCT |
| GPΔmuc-P8-Forward | GCCACTACGGAACTGCGAACGTTTCCCATTCTTAATAGGAAAGCGATAGAT |
| GPΔmuc-P8-Reverse | ATCTATCGCTTTCCTATTAAGAATGGGAAACGTTCGCAGTTCCGTAGTGGC |
| GPΔmuc-SS1a-Q5-Forward | TGCGCGGGTGATTTTGCTTGTCATAAAGAAGGCGCGTTTTTC |
| GPΔmuc-SS1a-Q5-Reverse | TGGTCCTGTCCCTGACACCTTGTG |
| GPΔmuc-SS1b-Q5-Forward | GGTCTTGCTTGGATACCCTGCTTCGGACCAGCCGCTGAAGGA |
| GPΔmuc-SS1b-Q5-Reverse | TATCGCAGCCCCTTCGTCTTGTGT |
| GPΔmuc-SS2a-Q5-Forward | AGGTGGGGTTTTAGGTCCTGTGTTCCCCCTAAGGTTGTTAAT |
| GPΔmuc-SS2a-Q5-Reverse | TTTGGTTGCTGATGGGACGTCAGT |
| GPΔmuc-SS2b-Q5-Forward | ATACCCTACTTCGGACCATGCGCTGAAGGAATCTACATA |
| GPΔmuc-SS2b-Q5-Reverse | CCAAGCAAGACCTATCGCAGCCCC |

| | |
|---|---|
| GPΔmuc-SS3a-Q5-Forward | AACTTGCATTATTGGACCTGTCAAGACGAAGGGGCTGCGATA |
| GPΔmuc-SS3a-Q5-Reverse | AGGGTTACATTTTGGCTGGGCATT |
| GPΔmuc-SS3b-Q5-Forward | CTCCAGCTCTTCCTCAGGTGCACTACGGAACTGCGAACGTTT |
| GPΔmuc-SS3b-Q5-Reverse | GGCTTGCGTGGTCTCATTGGCAAG |
| GPΔmuc-SS4a-Q5-Forward | TGCGCGGGTGATTTTGCTTTTCATAAAGAATGCGCGTTTTTCCTCTATG ACCGC |
| GPΔmuc-SS4a-Q5-Reverse | TGGTCCTGTCCCTGACACCTTGTG |
| GPΔmuc-SS4b-Q5-Forward | GCGATAGGTCTTGCTTGGTGTCCCTACTTCGGACCAGCCGCT |
| GPΔmuc-SS4b-Q5-Reverse | AGCCCCTTCGTCTTGTGTGGTCCA |
| GPΔmuc-SS5a-Q5-Forward | AACTTGCATTATTGGACCACACAATGCGAAGGGGCTGCGATAGGTCTT |
| GPΔmuc-SS5a-Q5-Reverse | AGGGTTACATTTTGGCTGGGCATT |
| GPΔmuc-SS5b-Q5-Forward | CTCCAGCTCTTCCTCAGGTGCACTACGGAACTGCGAACGTTT |
| GPΔmuc-SS5b-Q5-Reverse | GGCTTGCGTGGTCTCATTGGCAAG |
| GPΔmuc-SS6a-Q5-Forward | CAAGTTTCAGATGTGGACTGTCTTGTCTGTCGCGATAAACTT |
| GPΔmuc-SS6a-Q5-Reverse | AAGCGCGGAGTTGTGGATCACACC |
| GPΔmuc-SS6b-Q5-Forward | CTCCTCCAAAGGTGGGGCTGTACGTGTCATATATTGGGTCCC |
| GPΔmuc-SS6b-Q5-Reverse | GAAATCTATCGCTTTCCTATTAAG |

**b. Primers used in EBOV GP-presenting nanoparticle construct design**

| | |
|---|---|
| Ferritin-BamHI-Reverse | ATGATGGGATCCCTATTATGATTTGCGGCTCTTTGCGATACC |
| E2p-BamHI-Reverse | ATGATGGGATCCCTATTACATCAGAAGCAGCTCGGGGTCACT |
| I3-01-BamHI-Reverse | ATGATGGGATCCCTATTACTCAGTACATCCCCGTATTTT |
| LD1-BamHI-Reverse | ATGATGGGATCCCTATTACCGTCGCTCATTAACAACCACTTC |
| LD2-BamHI-Reverse | ATGATGGGATCCCTATTAGTGAGAGTCGAGGGCAATCTGCAA |
| LD3-BamHI-Reverse | ATGATGGGATCCCTATTACCGTTCCCTGTGTGGTGCGACTTC |
| LD4-BamHI-Reverse | ATGATGGGATCCCTATTACTTTTCAAACTGTGGGTGTGACCA |
| LD5-BamHI-Reverse | ATGATGGGATCCCTATTACCACAAGCACCCTTTGCATGCTTC |
| LD6-BamHI-Reverse | ATGATGGGATCCCTATTACTTATTCGTATACATTTCGAGGTA |
| LD7-BamHI-Reverse | ATGATGGGATCCCTATTAGTCTGCCCTGCGTGAGTTCCAGCG |
| LD8-BamHI-Reverse | ATGATGGGATCCCTATTATCGCCGTGCTTCGCGCAGTCGAGT |
| LD9-BamHI-Reverse | ATGATGGGATCCCTATTACTCATTTATGCTGCTCTGAAGTGC |
| PADRE-BamHI-Reverse | ATGATGGGATCCCTATTAGGCGGCGGCCTTCAGGGTCCAGGCGGC |

**c. Primers used in preparation of antibody chain libraries from mouse splenic B cells. [a]**

| | |
|---|---|
| 5'-RACE-P1-adaptor | CCACTACGCCTCCGCTTTCCTCTCTATGGGCAGTCGGTGATAAGCAGT GGTATCAACGCAGAGTAC |
| 3'-Cγ1-inner-A-adaptor [b] | CCATCTCATCCCTGCGTGTCTCCGACTCAG [barcode] GCTCAGGGAAATAGCCCTTGAC |
| 3'-Cγ2c-inner-A-adaptor [b] | CCATCTCATCCCTGCGTGTCTCCGACTCAG [barcode] GCTCAGGGAAATAACCCTTGAC |
| 3'-Cγ2b-inner-A-adaptor [b] | CCATCTCATCCCTGCGTGTCTCCGACTCAG [barcode] ACTCAGGGAAGTAGCCCTTGAC |
| 3'-Cγ3-inner-A-adaptor [b] | CCATCTCATCCCTGCGTGTCTCCGACTCAG [barcode] GCTCAGGGAAGTAGCCTTTGAC |
| 3'-Cμ-inner-A-adaptor [b] | CCATCTCATCCCTGCGTGTCTCCGACTCAG [barcode] AGGGGGAAGACATTTGGGAAGGAC |
| 3'-mCκ-outer-A-adaptor [b] | CCATCTCATCCCTGCGTGTCTCCGACTCAG [barcode] GATGGTGGGAAGATGGATACAGTT |

[a] The 3' primers are adapted from Tiller et al., *Journal of Immunological Methods* 350, 183-193 (2009).
[b] Ion Xpress Barcodes 1-20 [in blue] are used to differentiate antibody libraries from different mice.

**Supplementary Table 2. Data collection and refinement statistics for EBOV GPΔmuc trimers.**

| **Data Collection** | **GPΔmuc-WL$^2$-foldon** | **GPΔmuc-WL$^2$P$^2$-foldon** |
|---|---|---|
| Beamline | APS 23-ID-D | SSRL 12-2 |
| Wavelength (Å) | 1.0332 | 0.97946 |
| Resolution (Å)[a] | 49.05-2.28 (2.34-2.28) | 49.39-3.20 (3.26-3.20) |
| Space group | R32 | P321 |
| Unit cell (Å) | 114.58, 114.58, 312.38 | 114.06, 114.06, 136.22 |
| ( °) | 90 90 120 | 90 90 120 |
| Total reflections | 156,593 | 153,527 |
| Unique reflections | 34,904 (1725) | 17,343 (864) |
| Multiplicity | 4.5 (3.3) | 8.9 (8.7) |
| Completeness (%) | 96.4 (97.3) | 99.9 (99.8) |
| Mean (I)/ $\sigma_I$ | 8.0 (0.9) | 8.4 (0.45) |
| $R_{meas}$[c] (%) | 16.8 (147) | 31.9 (557) |
| $R_{pim}$[d] (%) | 7.4 (77.5) | 10.8 (187) |
| $CC_{1/2}$[e] (%) | 78.1 (32.5) | 77.6 (31.3) |
| **Refinement** | | |
| Refinement resolution (Å)[a] | 49.05-2.28 (2.34-2.28) | 49.39-3.20 (3.26-3.20) |
| # reflections in refinement (work/free) | 33188/1715 | 16245/837 |
| $R_{work}$ (%) | 19.7 (34.8) | 28.5 (51.3) |
| $R_{free}$ (%) | 23.4 (37.5) | 32.9 (48.2) |
| # Protein atoms | 2942 | 3123 |
| # Carbohydrate atoms | 153 | 139 |
| # Waters | 66 | 0 |
| # Protein residues | 374 | 398 |
| Bond r.m.s. deviation (Å) | 0.014 | 0.013 |
| Angle r.m.s. deviation (º) | 1.99 | 1.48 |
| Ramachandran favored, allowed, outliers (%)(*83*) | 96.9, 3.1, 0.0 | 92.1, 7.9, 0.0 |
| Clashscore[f] | 7.0 | 15.0 |
| Wilson B (Å$^2$) | 38 | 67 |
| Average B (Å$^2$) | 60 | 80 |
| Protein | 59 | 80 |
| PDB ID | 7JPI | 7JPH |

[a]Numbers in parentheses are for highest resolution shell

[b]$R_{merge} = \Sigma_{hkl} \Sigma_{i=1,n}|I_i(hkl) - \langle I(hkl)\rangle| / \Sigma_{hkl} \Sigma_{i=1,n} I_i(hkl)$

[c]$R_{meas} = \Sigma_{hkl} \sqrt{(n/n-1)} \Sigma_{i=1,n}|I_i(hkl) - \langle I(hkl)\rangle| / \Sigma_{hkl} \Sigma_{i=1,n} I_i(hkl)$

[d]$R_{pim} = \Sigma_{hkl} \sqrt{(1/n-1)} \Sigma_{i=1,n}|I_i(hkl) - \langle I(hkl)\rangle| / \Sigma_{hkl} \Sigma_{i=1,n} I_i(hkl)$

[e]$CC_{1/2}$ = Pearson Correlation Coefficient between two random half datasets

[f] Number of unfavorable all-atom steric overlaps ≥ 0.4Å per 1000 atoms
